# Archaeal *S*-adenosyl-l-homocysteine hydrolases: structure, function and substrate preferences

**DOI:** 10.1101/2023.05.12.540488

**Authors:** Désirée Popadić, Raspudin Saleem-Batcha, Lars-Hendrik Köppl, Philipp Germer, Jennifer N. Andexer

**Affiliations:** Institute of Pharmaceutical Sciences, University of Freiburg, Albertstr. 25, 79104 Freiburg, Germany

## Abstract

*S*-Adenosyl-l-homocysteine hydrolase (SAHH) reversibly cleaves *S*-adenosyl-l-homocysteine (SAH), the product of *S*-adenosyl-l-methionine (SAM)-dependent methylation reactions. The conversion of SAH into adenosine and l-homocysteine (Hcy) plays an important role in the regulation of the methyl cycle. An alternative metabolic route for SAM regeneration in the extremophiles *Methanocaldococcus jannaschii* and *Thermotoga maritima* was identified with the deamination of SAH to *S*-inosyl-l-homocysteine (SIH). Herein, we report the first structural characterisation of different archaeal SAHHs together with a biochemical analysis of various SAHHs from all three domains of life. We found that homologues deriving from the Euryarchaeota phylum show a higher conversion rate with SIH compared to SAH. Crystal structures of SAHH originating from *Pyrococcus furiosus* in complex with SIH and inosine as ligands, show architectural flexibility in the active site and offer deeper insights into the binding mode of hypoxanthine-containing substrates. Altogether, the findings presented in this study support the understanding of an alternative metabolic route for SAM and offer insights into the evolutionary progression and diversification of SAHHs involved in methyl and purine salvage pathways.

## Introduction

The methylation of small molecules as well as nucleic acids and proteins is an important modification in nature found in various biological processes such as drug metabolism, epigenetic regulation, and cancer development^1–3^. The methyl groups for such modifications are provided by a global metabolic pathway, the methyl cycle. The enzyme cofactor needed for this reaction is *S*-adenosyl-l-methionine (SAM), which is converted to *S*-adenosyl-l-homocysteine (SAH) by methyltransferases (MTs; EC 2.1.1.x) once the methyl group is transferred to a substrate, e.g. DNA, RNA, proteins, or small molecules^4,5^. MTs are a diverse group of enzymes installing the methyl group onto O, N, S, and C atoms, among others. In cells, the by-product SAH is salvaged as part of a complex regulation system, as SAH is known to act as a negative feedback inhibitor on most MTs, reducing the methylation rate in the cell^6^.

The SAM/SAH ratio is controlled by different enzymatic pathways (Fig. 1). 5’-Methylthioadenosine (MTA)/SAH nucleosidase (MTAN; EC 3.2.2.9) irreversibly cleaves the glycosidic bond of SAH to adenine and *S*-ribosyl-l-homocysteine (RibHcy), which is further transformed to l-homocysteine (Hcy) by the RibHcy lyase LuxS (EC 4.4.1.21)^7^. Alternatively, SAH hydrolase (SAHH; EC 3.3.1.1) reversibly converts SAH to adenosine and Hcy^7,8^. Hcy is re-methylated to l-methionine, a building block for SAM formation, by different enzymes^9–11^.

**Figure 1:**
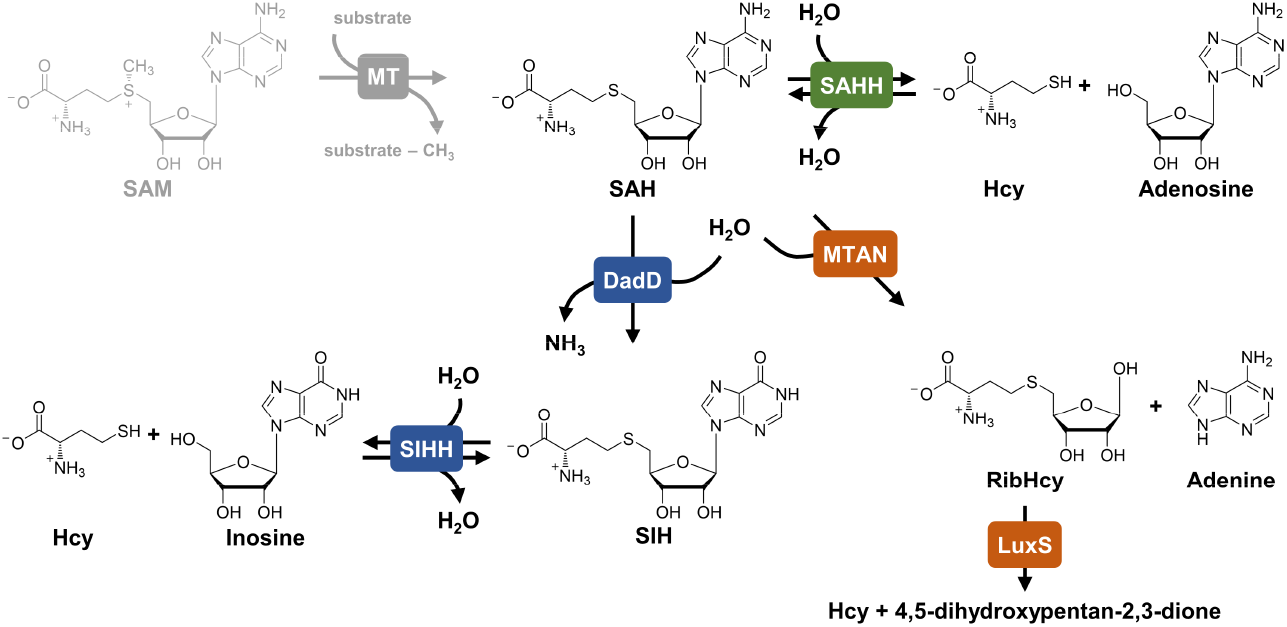
SAH degradation pathways. SAH is either reversibly cleaved to adenosine and Hcy by SAHH (green) or degraded in two steps catalysed by MTAN and LuxS (orange). The pathway discovered in *Methanocaldococcus jannaschii* containing the deamination step of SAH to SIH, which is subsequently cleaved to inosine and Hcy by SIHH (blue).

Some organisms feature both routes (MTAN/LuxS and SAHH) for SAH degradation, while others only possess one, or in rare cases, e.g. for *Mycoplasma genitalium*, neither pathway ^12^. A third pathway was identified in the archaeon *Methanocaldococcus jannaschii*. A 5’-deoxyadenosine deaminase (DadD; EC 3.5.4.41) catalyses the deamination of SAH to *S*-inosyl-l-homocysteine (SIH)^13^, which is subsequently hydrolysed to inosine (Ino) and Hcy by the SAHH homologue from *M. jannaschii*. For SAH, no activity was detected and therefore the enzyme was reclassified as an SIH hydrolase (SIHH; EC 3.13.1.9)^14^.

A homologue of DadD, an MTA/SAH deaminase (EC 3.5.4.31/28), was found in *Thermotoga maritima* in a structure-based activity prediction in 2007, indicating the same metabolic pathway as in the archaeon *M. jannaschii*. The SAHH from this thermophilic bacterium was shown to catalyse SIH hydrolysis in addition to SAH hydrolysis (with K_*M*_ values in the same order of magnitude)^15^. In 1988, a putative homologue of the deaminase was identified in *Streptomyces flocculus* (*Streptomyces albus* ATCC 13257) and proposed to be part of a major route in SAH metabolism, as SIH was isolated from the organism^16,17^. In addition to the SAHH homologues from *M. jannaschii*^14^ and *T. maritima*^15^ that were tested for SIH hydrolysis or synthesis activity, one bacterial representative from *Alcaligenes faecalis* was tested with nucleoside analogues in the synthesis reaction in 1984. With inosine, the enzyme showed almost no activity (0.5%) compared to adenosine as substrate^18^.

SAHHs are highly conserved in all domains of life and have been extensively characterised biochemically, as well as structurally, over the last century^19,20^. As the products of the SAHH-catalysed hydrolysis are rapidly removed *in vivo*, this reaction is preferred in cells, while SAH synthesis is the main reaction taking place *in vitro*^21,22^. SAHH depends on the cofactor nicotinamide adenine dinucleotide in its oxidised form (NAD^+^), which is self-regenerated during the catalytic cycle^23–26^. Briefly, the 3’-OH group of the adenosine ribose is oxidised to a ketone by reducing NAD^+^ to NADH. The now more acidic C4’ proton is abstracted to form a carbanion intermediate, followed by the release of Hcy. Water is added through a Michael-type addition to the C4’–C5’ double bond^23^. The last step is the reduction of the ketone to form adenosine under re-oxidation of NADH to NAD^+^; thus, the cofactor is ready for the next reaction cycle either in the hydrolysis or synthesis direction.

The protein structure of SAHH features three domains in a monomer, the active form is usually a homotetramer^19^ (Figs. 2A and 2B). The substrate-binding domain is located next to the cofactor-binding domain, each showing a Rossmann fold^19^. The C-terminus is a smaller dimerisation domain. The substrate-binding and the cofactor-binding domains are connected by a two-part hinge element. SAHHs have been shown to alternate between two conformations differing in the relative positions of the substrate- and cofactor-binding domains (Fig. 2C). In the “open” conformation, with no substrate bound, the domains are away from each other providing access to the active site (PDB IDs: 3X2F and 4LVC)^27,28^. In the “closed” conformation, the substrate-binding domain reorients by about 18° relative to the cofactor-binding domain and both domains form the active site interacting with a substrate or inhibitor^29,30^. In addition of the overall conformational state, a critical loop region comprising a pair of histidine (His) and phenylalanine (Phe) act as gate residues that provide a channel for the substrate to access the active site; it is highly conserved over SAHH sequences of all domains of life^31^. This so called “molecular gate” loop displays a plasticity that opens (His-OUT) and shuts (His-IN) upon different ligation states *via* a mechanism through a 180° flip of the peptide plane between Cα atoms of His and Phe (Fig. 2C)^31^.

**Figure 2:**
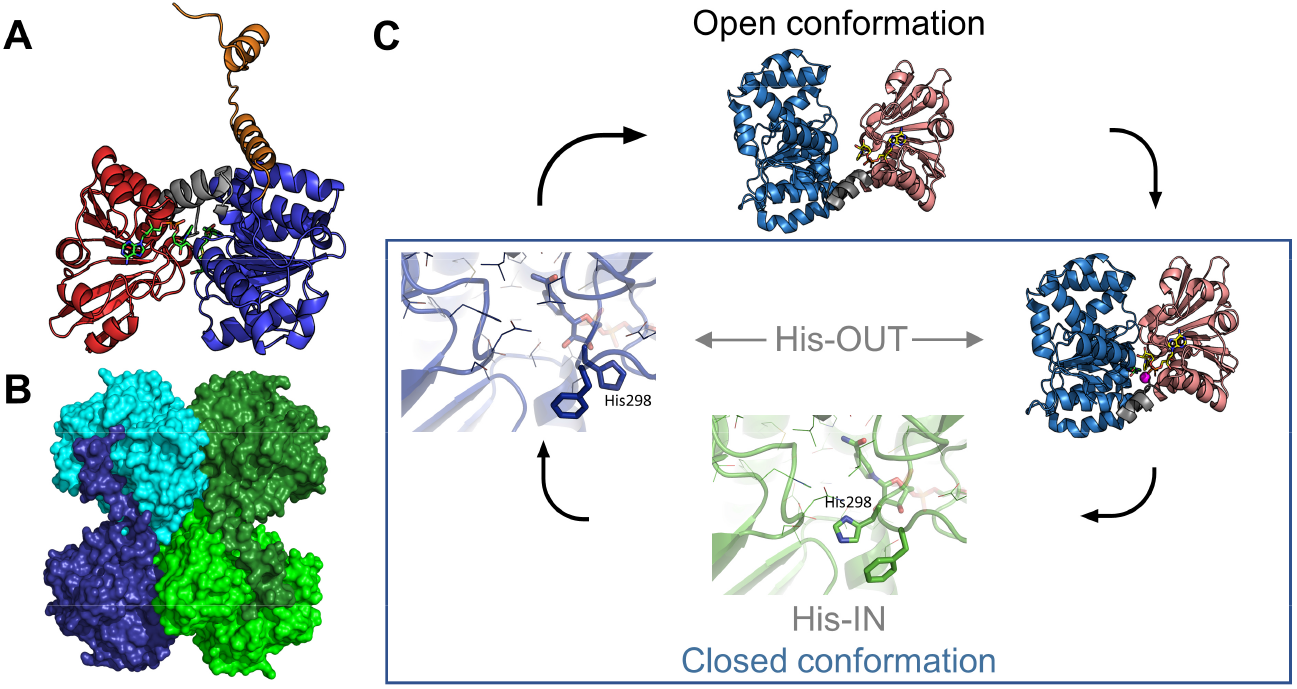
Structural features of SAHHs. A: An enzyme monomer consists of the C-terminal dimerisation domain (orange), the substrate-binding domain (blue) and the cofactor-binding domain (red) connected by a two-part hinge element (grey) (PDB ID: 7R38). B: The monomers oligomerise to form a dimer of a dimer (PDB ID: 7R38). C: The monomers alter their conformation between the open and closed state (PDB ID: 4LVC). Within the closed conformation, a His can act as gate residue (PDB IDs: 7R37 and 7R38).

Multiple protein structures from bacterial and eukaryotic SAHH enzymes have been determined^26,28,31,32^ (Supplementary Table S7 online); however, no protein structure originating from the domain of Archaea has been published. The archaeal enzymes are predicted to show similar architecture as their eukaryotic and bacterial relatives but display remote differences such as a shortened C-terminus and a missing 40 amino acid segment in the catalytic domain among other smaller deletions^12,33^. Further, an HxTxQ(E) sequence signature (with x corresponding to any amino acid) is found in eukaryotic and bacterial enzymes in their substrate-binding domain, while extremophile SAHHs have been suggested to feature a different motif, HxT(E)xK^19^. The His residue is important for catalysis (Supplementary Fig. S17 online), while Thr(Glu) and Gln(Glu) have been suggested to stabilise the nucleobase in the active site *via* hydrogen bonds^19,25^. The residue Gln(Glu) was also found to play a role in the conformational changes of the enzyme, regulating its activity^34^.

The discovery of SIHHs^14^ prompted us to study the tertiary structures and biochemical properties of SAHH enzymes from archaea and to identify their differences from bacterial and eukaryotic homologues characterised biochemically. Here, we show the first archaeal SAHH/SIHH crystal structures within the Euryarchaeota and Crenarchaeota phyla in the presence of hypoxanthine derivatives (inosine and SIH) of the recently proposed alternate SAM salvage pathway^13^. In addition, our investigation of the substrate ranges of different SAHHs led to the identification of more organisms potentially featuring alternative SAM regeneration pathways. Altogether these findings result in a substantial support of the alternate SAM salvage pathway and the importance of methyl metabolism in archaea.

## Results and Discussion

### Overview of investigated SAHHs/SIHHs

In this work, SAHHs/SIHHs from archaea, bacteria, and eukaryotes were characterised biochemically (Supplementary Table S4 online). The homologues from *M. jannaschii* (*Mj*SIHH) and *T. maritima* (*Tm*SAHH) were previously reported to have SIH hydrolysis activity ^14,15,27,35^. Structures of archaeal SAHHs were determined from three organisms: *Methanococcus maripaludis* (*Mma*SAHH), *Pyrococcus furiosus* (*Pfu*SAHH)^36^, and *Sulfolobus acidocaldarius* (*Sac*SAHH). In addition, the structure of SAHH from *Mus musculus* (*Mm*SAHH) was determined in complex with the alternative substrate inosine.

### The substrate range differs depending on the origin of the enzyme

All enzymes were tested in both reaction directions with the hypoxanthine- and adenine-containing substrates. As SAH/SIH hydrolysis is not preferred *in vitro* and consequently hard to follow, the reaction was coupled to Hcy *S*-methylation catalysed by Hcy *S*-MT (HSMT; EC 2.1.1.10)^9^ to drive the reaction forward by removing one of the products (Figs. 3B-D). This still allows the analysis with HPLC-DAD compared to previous coupled colorimetric and photometric assays for SAHHs^37,38^. As a side-product, the corresponding nucleobase (adenine or hypoxanthine) was detected in all set-ups, as previously described^39^ (Figs. 3B and 3C). In some reports, SAHH activity was only observed with externally added NAD^+40–42^, while there are also examples showing SAH hydrolysis and synthesis activity without the addition of extra NAD^+32,36,43^. All enzymes tested in this work were active without the addition of NAD^+^.

**Figure 3:**
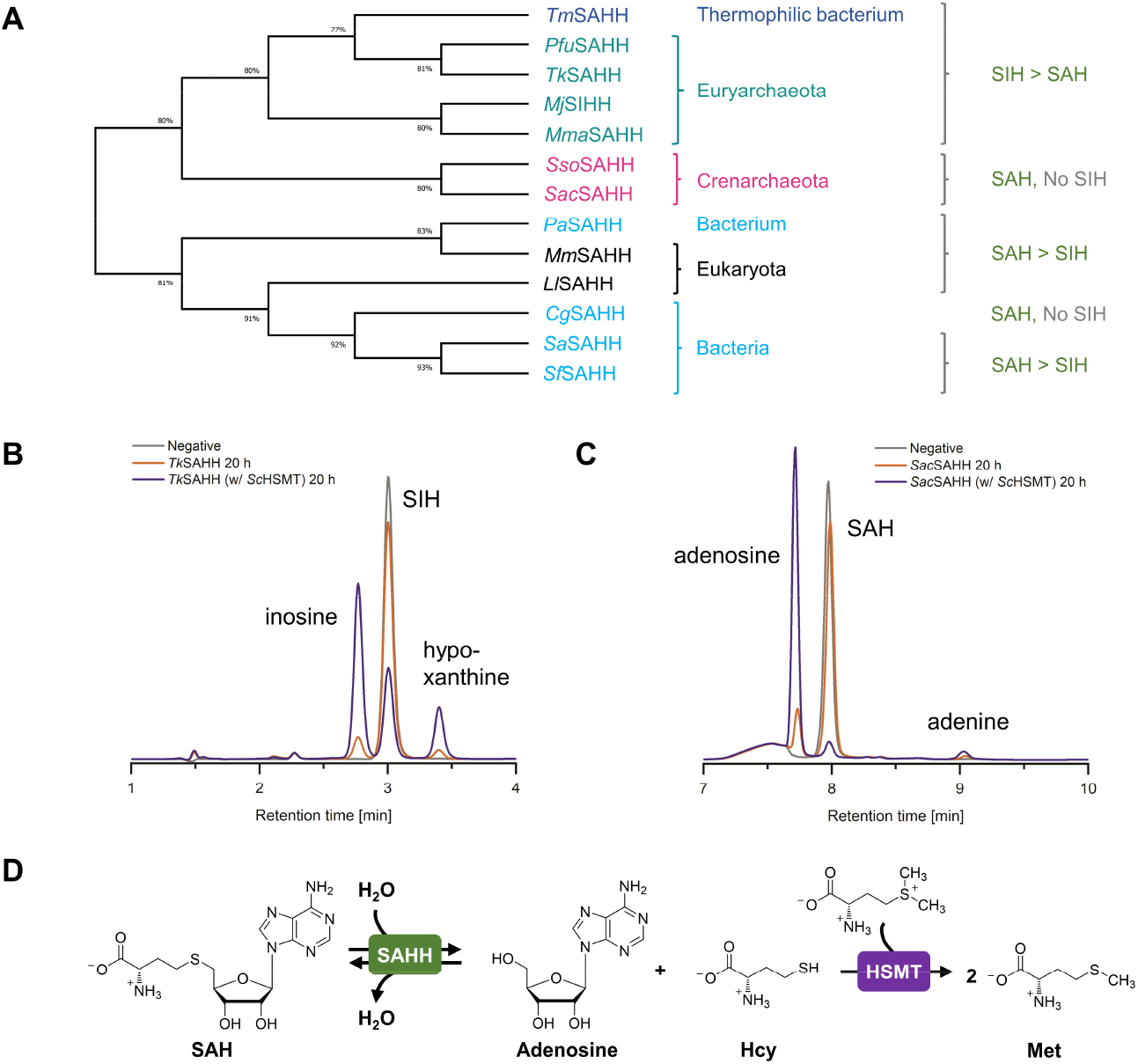
Phylogenetic tree of investigated SAHHs/SIHHs with substrate range and HPLC analysis of the hydrolysis reactions. A: Representatives from Crenarchaeota only accept SAH as substrate, while family members from the Euryarchaeota and a closely related thermophilic bacterium accept both SAH and SIH, while preferring SIH. Except for *Cg*SAHH, bacterial and eukaryotic homologues accept both substrates with a clear preference for SAH. B: *Tk*SAHH-catalysed cleavage of SIH, and C: *Sac*SAHH-catalysed cleavage of SAH, both with and without addition of *Sc*HSMT (negative control without enzymes). D: The addition of the second enzyme *Sc*HSMT increases the conversion of the hydrolysis reaction.

Under the chosen conditions, all SAHHs/SIHHs were active for SAH hydrolysis and synthesis (Table 1), including the homologue from *M. jannaschii* (*Mj*SIHH) previously described to be specific for SIH as substrate^14^ (Supplementary Fig. S4; all chromatograms in Figs. S4-S16 online). Nevertheless, this enzyme clearly prefers the hypoxanthine-containing compounds over the adenine-containing ones, the same results were observed for the other representatives from Euryarchaeota (*Mma*SAHH, *Pfu*SAHH, and *Tk*SAHH). In contrast, SAHHs from Crenarchaeota (*Sac*SAHH and *Sso*SAHH) showed no activity with SIH (Figs. S8 and S9 online). As described before, the bacterial *Tm*SAHH, which is closely related to euryarchaeal enzymes, catalysed SIH hydrolysis and synthesis^15^, in our hands also with a strong preference for the hypoxanthine derivatives. These findings strongly support the assumption of alternative routes for methyl metabolism and purine salvage within Euryarchaeota and closely related bacteria.

**Table 1:**
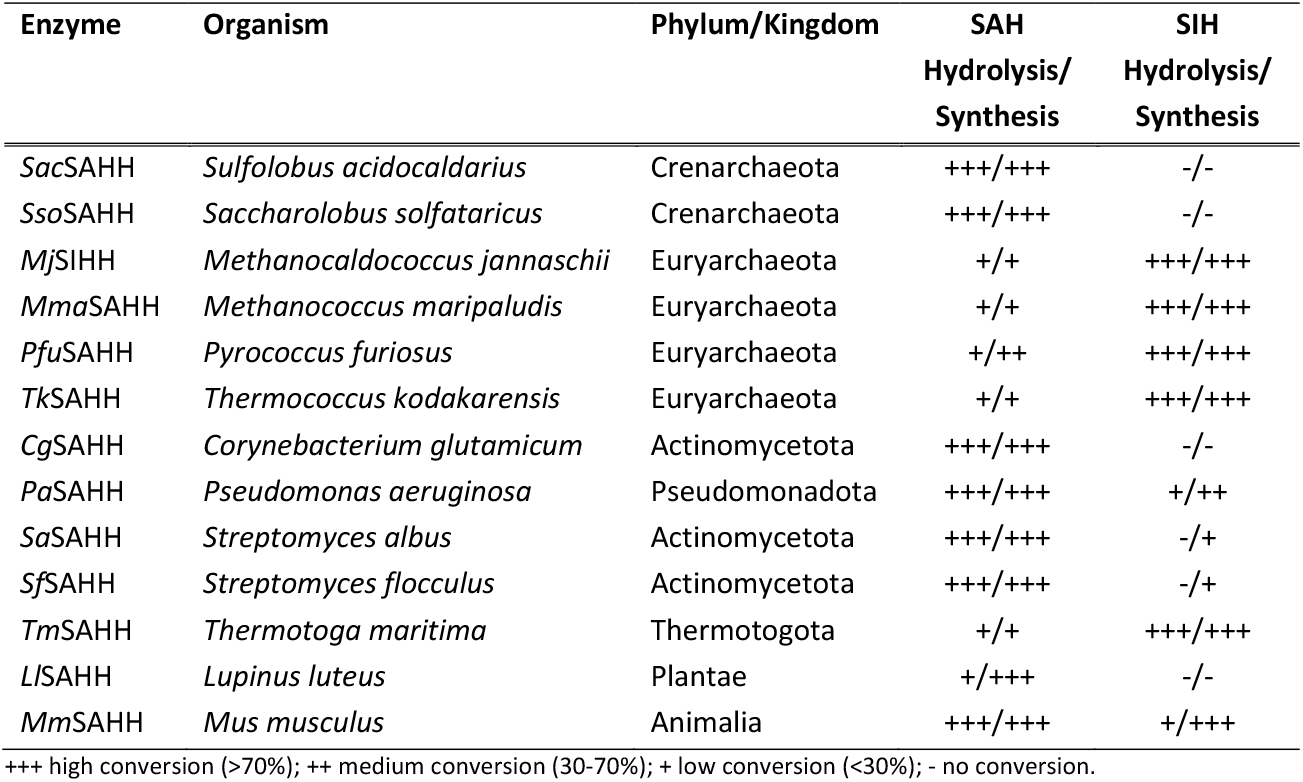
Overview of biochemically tested SAHHs in this work. Conversions indicated for the hydrolysis are based on reactions coupled to *Sc*HSMT.

In previous work, we successfully used the SAHH from mouse (*Mm*SAHH) for an *in vitro* SAM regeneration cycle with alternative nucleobases including hypoxanthine^44^. This activity was now confirmed in the individual hydrolysis and synthesis reactions; in contrast to the euryarchaeal enzyme, reactions with SAH were in this case preferred. A similar pattern was observed for the SAHH from the bacterium *Pseudomonas aeruginosa*, while neither the representative from *Corynebacterium glutamicum* nor from the plant *Lupinus luteus* accepted inosine or SIH as substrates. As SIH had been described as a metabolite in *S. flocculus*, along with the presence of a SAH deaminase^16,17^, we tested SAHHs from two *Streptomyces* species; in both cases a low activity for SIH synthesis was detected, but no activity for SIH hydrolysis (Figs. S15 and S16 online). It might be that SIH is metabolised by other enzymes in these bacteria, suggesting an alternative SAH metabolism pathway than the one described for Euryarchaeota and archaeal-type bacteria. It would be interesting to investigate this in detail by sequencing the strains of *Streptomyces albus* and *S. flocculus* in the future.

### Genomes from Euryarchaeota and archaeal-type bacteria encode a deaminase

In order to get more insight in the potential alternative SAH salvage pathway, a BLASTP^45^ search for a protein homologue of DadD from *Methanocaldococcus jannaschii* (*Mj*DadD) in the organisms producing the SAHHs/SIHHs analysed in this work was performed. The genomes from the Euryarchaeota, as well as from the archaeal-type bacterium *Thermotoga maritima*, were found to have a homologue encoded (Supplementary Table S5 online). This is consistent with the characterised archaeal SAHHs/SIHHs hydrolysing and synthesising SIH in addition to SAH. A more detailed BLASTP search including all available entries in the UniProt database^46^ resulted in the identification of further homologues of *Mj*DadD, exclusively within the domain of bacteria and the phylum of Euryarchaeota. This suggests that the alternative SAM regeneration pathway going through SIH is indeed not present in all archaea, but only the phylum of Euryarchaeota and closely related thermophilic bacteria. Thermophilic bacteria were shown to have obtained genes *via* horizontal transfer from archaea^47^. The organisms without an encoded SAH deaminase do not need their SAHH to degrade SIH in addition to SAH. The reason for substantial SIH degradation and synthesis by *Mm*SAHH and *Pa*SAHH remains yet unclear.

### Architecture and domains of archaeal SAHHs

So far, the molecular basis for the differences in substrate ranges among the investigated SAHHs/SIHHs remains elusive and suggested a detailed structural comparison. The archaeal enzymes were an ideal model system as they show a clear cut of substrate preference between the phyla of Crenarchaeota and Euryarchaeota. As no structures of archaeal SAHHs have been published to date, we present here the structures of three archaeal SAHHs (*Pfu*SAHH, *Mma*SAHH, *Sac*SAHH); in addition, the structure of the eukaryotic *Mm*SAHH was determined in complex with its unusual substrate inosine. In all SAHH structures obtained, the NAD^+^ cofactor was present indicating that it was co-purified with the enzyme from the host expression organism. The crystal structures of *Pfu*SAHH complexes belong to the *P*4_2_2_1_2 space group with one homodimer in the asymmetric unit. Both complexes were elucidated by the molecular replacement (MR) method using chain A of the PDB entry 5AXA as search model^26^. The *Pfu*SAHH•NAD•inosine complex (PDB ID: 7R37) and the *Pfu*SAHH•NAD•SIH complex (PDB ID: 7R38) were refined to resolutions up to 2.3 Å and 2.0 Å, respectively. The crystal structure of *Mma*SAHH•NAD•inosine complex (PDB ID: 7R3A) belongs to *P*2_1_ space group with four homodimers in an asymmetric unit. The complex was elucidated by MR using PDB entry 1V8B chain A as search model, and subsequently refined to a resolution up to 2.5 Å^48^. The crystal structure of the *Sac*SAHH•NAD•adenosine complex (PDB ID: 7R39) belongs to *P*1 space group with four homodimers in an asymmetric unit. This complex was elucidated by MR using PDB entry 3H9U chain A as search model, and subsequently refined to a resolution up to 2.6 Å^49^. The crystal structure of *Mm*SAHH•NAD•inosine complex (PDB ID: 8COD) belongs to *I*222 space group with one homodimer in an asymmetric unit. The complex was elucidated by MR using PDB entry 5AXA chain A as search model, and subsequently refined to a resolution up to 2.48 Å^26^. The molecules in the asymmetric unit in all the cases were almost identical, being superimposable with a root-mean-square deviation (r.m.s.d.) of 0.17-0.63 Å over 356–393 C_α_ atoms. Data collection and refinement statistics are summarised in Supplementary Table S3. Furthermore, multimeric state analyses of SAHH structures from this study with PDBePISA^50^ indicate that they form stable homotetramers in their biological assembly.

The archaeal SAHH monomer contains the three domains as previously studied SAHHs across other domains of life: substrate-binding domain, cofactor-binding domain, and C-terminal domain^27,32,34,42,51^. The binding modes of the NAD^+^ at the cofactor-binding domain and inosine or adenosine in substrate-binding domain of archaeal SAHHs are similar to those observed in other SAHHs of distinct origin (Fig. 4)^26,28,31,32,34,48,52^. In this study, the nucleoside moieties of the three ligands, adenosine, inosine, and SIH, participate in a similar pattern of hydrogen-bonding and non-bonding interactions. The adenine moiety of adenosine in the *Sac*SAHH•NAD•adenosine complex is interfaced by specific hydrogen bonds, including N1 to Oδ1 of Thr53, and N6 to the main-chain O of His344 and Oδ1 of Glu55. The binding of adenine is additionally supported by non-bonded contacts with Leu338, Met349, and Thr56. In the ribose moiety, only the O3’ is hydrogen bonded to the amino group of Lys177. These interactions are accompanied by non-bonded contacts to Asp181, Glu147, and Thr56. The O5’ forms a hydrogen bond with His51, Nε2 (Supplementary Fig. S19D online). In the *Mma*SAHH•NAD•inosine complex, the hypoxanthine moiety of inosine is recognised by multiple hydrogen bonds, including N7 to main-chain O of His364, O6 to Oδ1 of Lys74, and N1 to Oδ1 of Glu72, and additional non-bonded contacts from Thr75, Leu358, Gly363, Met369, and Phe373. The O2’ and O3’ of the ribose moiety hydrogen bond with Lys198, and there are non-bonded contacts with Asn143, Glu168, Leu355, and NAD502. The O5’ of inosine hydrogen bonds with Nδ1 of His70 and Nε2 of His313 (Supplementary Fig. S19C online). In the *Pfu*SAHH•NAD•inosine complex, the hypoxanthine moiety of inosine is linked by specific hydrogen bonds, including N7 to main-chain O of His350, O6 to Oδ1 of Lys59, and N1 to Oδ1 of Glu57, and additional non-bonded contacts from Thr60, Leu344, Gly349, Met355, and Phe359. The O3’ of the ribose moiety hydrogen bonds with Lys183 and Thr154. Additional non-bonded contacts include Glu153, Leu341, and NAD601. The O5’ of inosine hydrogen bonds with Nδ1 of His70 Nδ1 and His313 Nε2 (Supplementary Fig. S19A online). The hypoxanthine moiety of SIH in *Pfu*SAHH•NAD•SIH complex makes similar contacts as in the *Pfu*SAHH•NAD•inosine complex, with additional non-bonded interactions from Asp187 in the *Pfu*SAHH•NAD•SIH complex. Expectedly, considerable differences are found at the site of binding of the amino acid moiety of SIH. The hydrogen bonds include Sδ with Nε2 of His55, O with Nδ2 of Asn80, OXT with main-chain O of Phe299, and N with main-chain O of Asp128 and main-chain O and Oδ of Ser79 (Supplementary Fig. S19B online). In the *Mm*SAHH•NAD•inosine complex, the hypoxanthine moiety of inosine is recognised by multiple hydrogen bonds, including N1 to Oγ1 of Thr57, N7 to main-chain N of His353, and weak bonds between O6 to Oε1 of Glu39. Additional non-bonded contacts from Leu54, Glu59, Thr60, Leu347, Gly352, and Met358. The O3’ of the ribose moiety hydrogen bonds with Nζ of Lys186, Oγ1 of Thr157 and there are non-bonded contacts with Asp131, Glu156, Asp190, and NAD601. The O5’ of inosine hydrogen bonds with Nε2 of His55 and Nδ1 of His301 (Supplementary Fig. S19E online).

**Figure 4:**
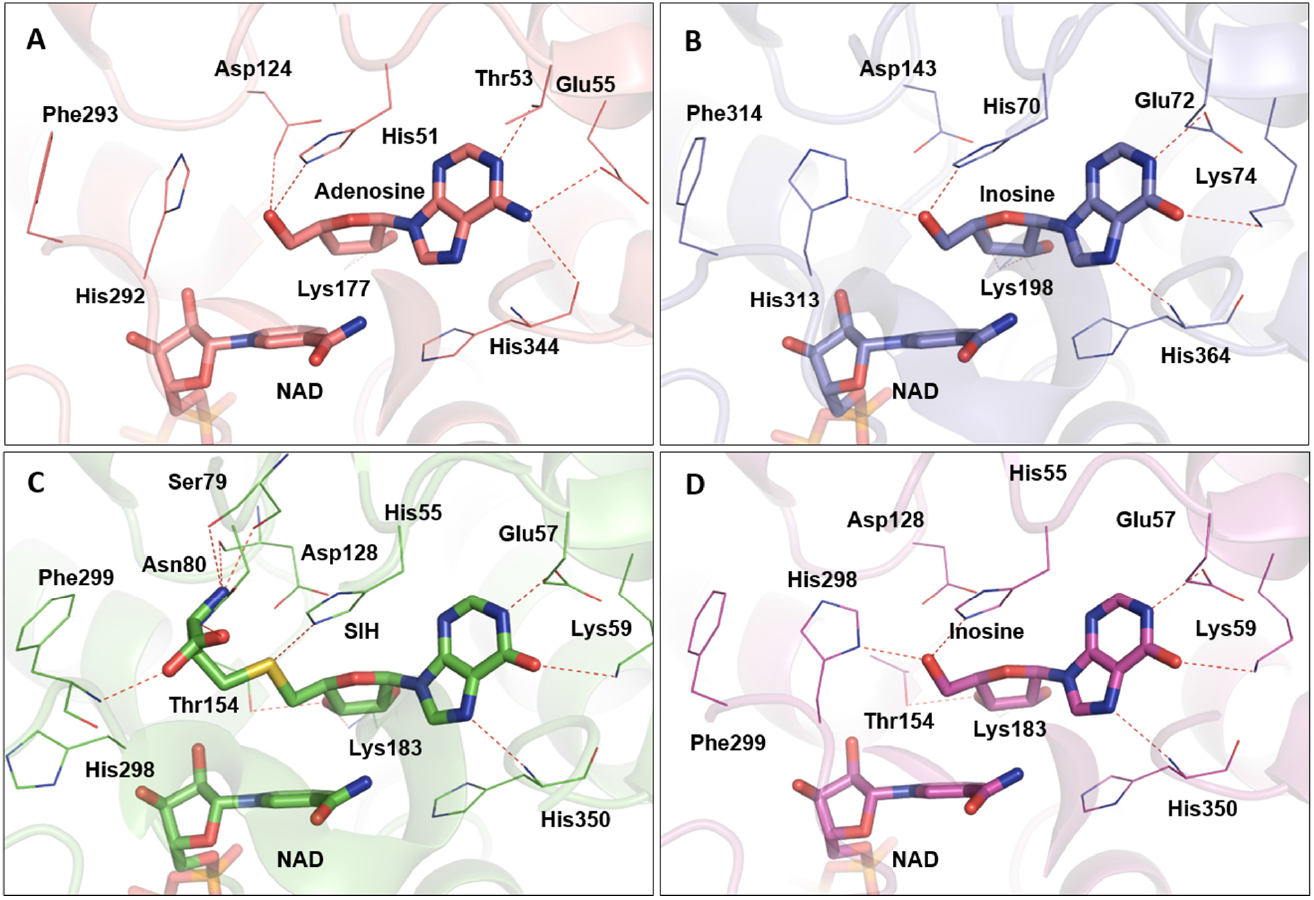
Modes of substrates/product and cofactor binding in archaeal SAHHs (as sticks). A: Mode of adenosine and NAD^+^ cofactor binding in the active site of *Sac*SAHH (PDB ID: 7R39). B: Mode of inosine and NAD^+^ cofactor binding in the active site of *Mma*SAHH (PDB ID: 7R3A). C: Mode of SIH and NAD^+^ cofactor binding in the active site of *Pfu*SAHH (PDB ID: 7R38). Mode of inosine and NAD^+^ cofactor binding in the active site of *Pfu*SAHH (PDB ID: 7R37).

### Different motifs for binding the nucleobase of the substrate/product in the subgroups of archaea

Based on our analysis, the fingerprint motif suggested to distinguish extremophile SAHHs from mesophilic enzymes^19^ can be further specified: Crenarchaeota and a large part of Euryarcheoata feature HxTxE as a signature, matching the sequence signature of bacterial and eukaryotic representatives (HxTxE(Q)); a subgroup of the euryarchaeal and archaeal-type bacterial SAHHs, including the ones analysed in this study have an HxExK motif (Fig. 5A; full sequence alignment in Supplementary Fig. S18 online). The subgroup showing the HxExK motif encompasses mainly the *Methanococci* and *Thermococci;* while euryarchaeal enzymes from various other classes (e.g. *Methanopyri)* also feature the HxTxE motif as found in Crenarchaeota; in other classes (e.g. *Methanomicrobia*), there is no clear trend visible. Our experiments show that the homologues with a HxExK motif prefer SIH as substrate while enzymes with a HxTxE(Q) motif prefer SAH. Evaluating the euryarchaeal structures elucidated in this study, it supports the assumption that the Glu residue (for *Pfu*SAHH, Fig. 5B) forms a hydrogen bond with the heterocyclic nitrogen atom N1 of the nucleoside, analogous to the Thr residue in crenarchaeal, bacterial and eukaryotic homologues (e.g. *Sac*SAHH, Supplementary Fig. S19D online). The Lys residue is found in enzymes that prefer hypoxanthine-containing substrates; here, the positively charged side chain can form a hydrogen bond with the oxygen at C6 in inosine, as seen for *Pfu*SAHH (Figs. 5B and S19A online) and *Mma*SAHH (Fig. S19C online), contributing to the stabilisation of the substrates in the active site. The motif HxTxE found in *Sac*SAHH (Crenarcheota) shows the same interactions with the substrate adenosine as in bacterial and eukaryotic enzymes^19^. As described before^19^, we observed that the *exo*-amino group of adenosine bound in *Sac*SAHH forms additional hydrogen bonds with the carbonyl oxygen atoms in the main chains of Glu342 and His344 (numbering according to *Sac*SAHH) indicating that the adenine ring is in its preferred tautomeric amino form. Regarding the structures of *Pfu*SAHH and *Mma*SAHH with their preferred substrates inosine or SIH, the oxygen attached to C6 seems to form additional hydrogen bonds with the carbonyl oxygen atoms of Asp348 (*Pfu*SAHH) or Asp362 (*Mma*SAHH) in the main chains. This suggests that the hypoxanthine ring is in the imino-hydroxy form (Fig. 5C), although inosine has been shown to be most stable in the amino-oxo form in water^53^. Further studies regarding the differences in the signature sequences for the stabilisation of the nucleobase moiety will be needed to determine their influences on the substrate preference of different SAHH/SIHH homologues.

**Figure 5:**
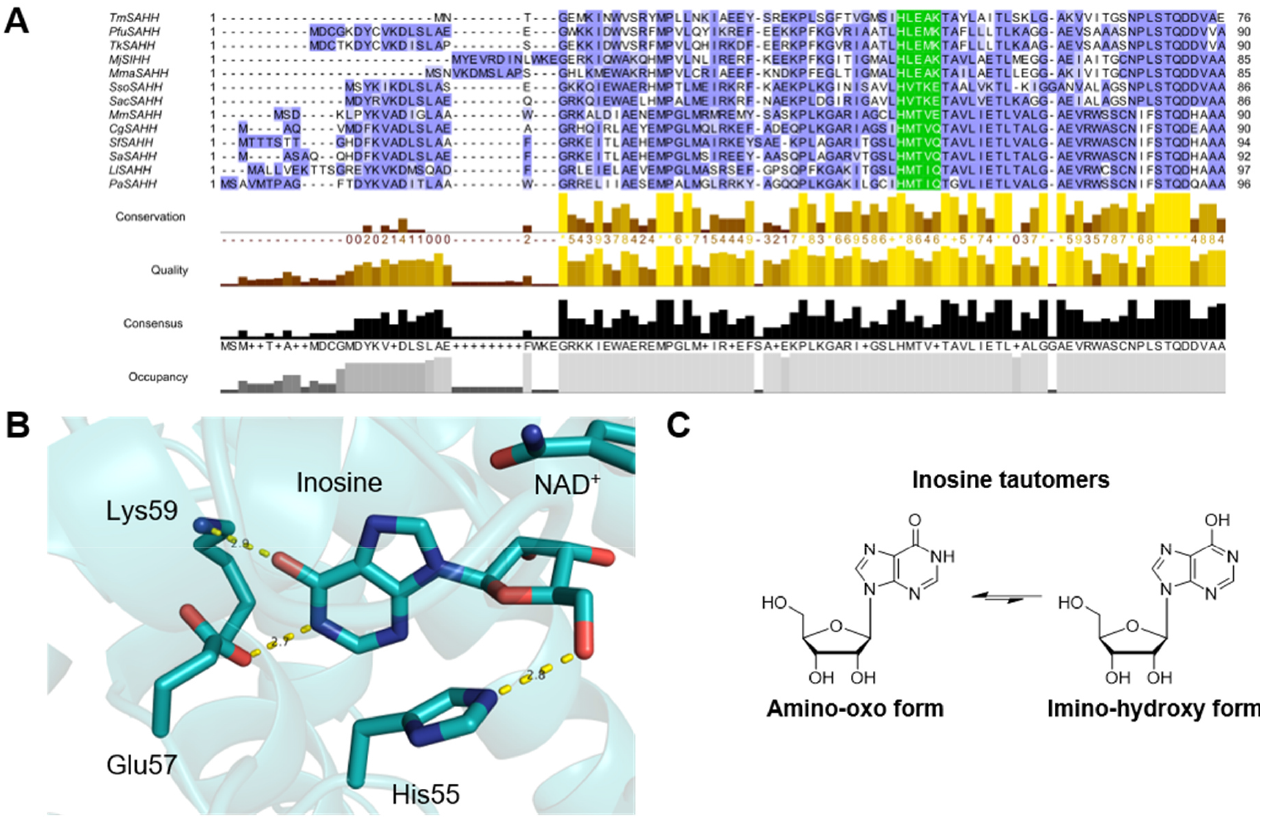
Structural analysis of substrate-binding domain of *Pfu*SAHH. A: Partial alignment of the amino acid sequences of SAHHs tested in this study. The binding motifs are coloured in green while the other residues are coloured by conservation according to the BLOSUM62 algorithm. B: Inosine is bound by the motif HxExK in the substrate-binding domain (cartoon representation; PDB ID: 7R37). C: Tautomers of inosine.

### Overall conformational states and the molecular gate of archaeal SAHHs

Different conformational states of SAHHs have been described, where the open conformation was observed with no bound substrate/product in the substrate-binding domain, while the protein with a bound substrate or analogue shows the overall closed conformation. An example is an unpublished data set (PDB ID: 5UTU) of a protistan/eukaryotic SAHH showing SAH, the substrate for the hydrolysis reaction, bound in a closed conformation^54^. All archaeal SAHH complexes in this study also display a closed conformational state including the SIH bound *Pfu*SAHH complex.

In addition to the closed overall conformation of the *Pfu*SAHH•NAD•SIH complex, the molecular gate loop displays a flexibility. The transitions between the overall conformation and the conformation of the gatekeeper residues are independent from each other. Yet, His298 and Phe299 forming the molecular gate in *Pfu*SAHH can only act as gatekeepers in the closed conformation when substrate-binding domain and cofactor-binding domain form the channel, as described before ^28,31^.

Interestingly, the His298 residue alone displays a noticeable conformational difference in this study. Both, His-IN and His-OUT conformations correlate with the access of a channel for the substrate or product to enter or leave the domain, respectively. In the inosine bound complex, His298 is situated within the active site staging a His-IN conformation (gate shut) where its imidazole ring is oriented towards the γ-carboxylate group of Asp128 with a distance of 4.2 Å. The Nδ1 group of His298 is oriented to the O5’ group of inosine with a distance of 2.7 Å. A similar pattern is observed for *Mma*SAHH and *Sac*SAHH complexes. However, the rotamer conformation of His in the His-IN state of *Sac*SAHH deviates from the rest of the His-IN conformations among the archaeal SAHHs (Fig. 6) in this study. This indicates that there are not only different His orientations possible in the His-OUT state^55^, but also in the His-IN state. Comparison of the *Mm*SAHH structures with bound adenosine and inosine (Supplementary Fig. S19G online) and the structure of *Sac*SAHH suggests that the rotamer conformation is not dependent on the bound substrate. The His298 residue of *PfuS*AHH•NAD•SIH complex swings away from Asp128 by 10.4 Å and forms a hydrogen bond with the carboxylate group of Glu302 with a distance of 2.7 Å signifying a His-OUT conformation leaving an open molecular gate. In a previously described structure of the *Lupinus luteus* SAHH^32^, the channel is open (His-OUT) even though adenosine is bound, while in the SAHH structure from *Mycobacterium tuberculosis* the same trend is seen as for *Pfu*SAHH with the channel closed (His-IN) with adenosine bound and channel opened with SAH bound^31^. Similarly, our complexes of *Pfu*SAHH and *Mma*SAHH with inosine and complex of *Sac*SAHH with adenosine show closed conformation state of the molecular gate in their active site (Fig. 6). Comparison of our structures with already published SAHH structures did not lead to an apparent correlation between the conformation of the gate residues and the bound substrates. Instead, the state of the reaction catalysed by SAHHs may determine the conformation of the gatekeeper residues. Yang *et al*. proposed that the active site is sealed from the bulk solvent after the oxidation reaction at the 3’-position^56^.

**Figure 6:**
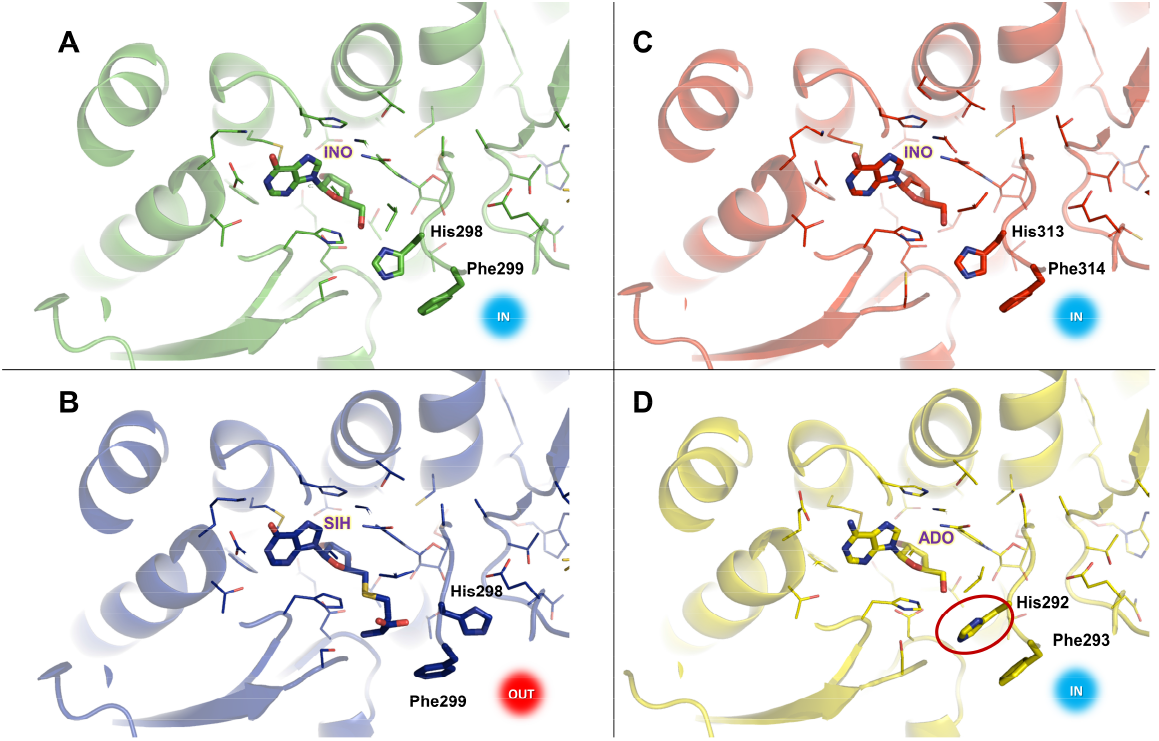
The molecular gate residues in the crystal structures of archaeal SAHHs from this study. The His residue states are represented as His-IN (IN) and His-OUT (OUT). A: In the *Pfu*SAHH•NAD•inosine complex (PDB ID: 7R37), the His gatekeeper residue is in the IN orientation thereby closing the channel entrance. B: In the *Pfu*SAHH•NAD•SIH complex (PDB ID: 7R38), the gatekeeper residue forms an OUT orientation leaving the channel entrance open. C: For the *Mma*SAHH•NAD•inosine complex (PDB ID: 7R3A) the gatekeeper residue is in the IN orientation. D: In the *Sac*SAHH•NAD•adenosine complex (PDB ID: 7R39), the gatekeeper residue is in the IN orientation that makes the channel entrance shut and inaccessible. The side chain of the gatekeeper residue of *Sac*SAHH (circled red) shows another rotamer conformation than in *Pfu*SAHH and *Mma*SAHH.

### Differences in archaeal compared to bacterial and eukaryotic SAHHs

While the protein sequences of archaeal SAHHs are similar throughout the distinct phyla (Supplementary Table S6 online), the substrate range differs. The first archaeal structures of SAHHs were determined to get insights into the structural differences: two stemming from Euryarchaeota (*Mma*SAHH and *Pfu*SAHH) and one from Crenarchaeota (*Sac*SAHH). The structure of the eukaryotic *Mm*SAHH was solved in complex with inosine to get insights into the binding of its disfavoured substrate, also in comparison to homologues, which favour binding of hypoxanthine-containing substrates. Looking at the overall structure of archaeal SAHHs, the major difference is the shortened C-terminus, which interacts with the cofactor of a second subunit. Due to the shortened C-terminus, the archaeal enzyme structures lack hydrogen bonds, which are formed between amino acids (Lys426 and Tyr430) of one subunit with the 2’- and 3’-hydroxy groups of the adenosine moiety and the pyrophosphate of NAD^+^ bound in the adjacent subunit, present in *Mm*SAHH and other mesophilic homologues. In addition, the interfaces between the monomers of the archaeal SAHHs show a cleft, when looking at the surface representation (Fig. 7A). This is not visible in eukaryotic SAHHs, such as the one from mouse (Fig. 7B); also because of the longer C-terminus that covers the cofactor. This means the cofactor is more exposed to the environment in archaea, and indicates a less stable tetrameric form for archaea, as fewer interactions are found between the monomers. *Tm*SAHH was found to require a much higher concentration of NAD+, which was linked back to the shortened C-terminus, impacting the binding affinity^27^. *Tm*SAHH has the shortest C-terminus, which is three amino acids shorter than the archaeal ones. All SAHHs/SIHHs in this work (including *Tm*SAHH) were active without the addition of NAD^+^ at 37 °C and thus no dependency between concentration of NAD^+^ and length of C-terminus was observed in our hands. To verify the arrangement of individual units in the multimeric assembly of the archaeal SAHH crystal structures, tetramers were constructed and analysed for each SAHH individually in their respective space groups using PDBePISA^50^. The analysis of *Pfu*SAHH tetramers (both inosine and SIH complexes) showed an average value for ΔG^int^ (ΔG for the interaction) of -129.0 kcal mol^-1^. Similarly, *Mma*SAHH tetramers exhibited an average ΔG^int^ of -133.5 kcal mol^-1^, while *Sac*SAHH tetramers had an average ΔG^int^ of -127.5 kcal mol^-1^. In comparison, selected long C-terminus tailed eukaryotic SAHHs such as *Mm*SAHH tetramers (both inosine and adenosine complexes) showed an average ΔG^int^ of -186.7 kcal mol^-1^, while SAHH from *Cryptosporidium parvum* (*Cpa*SAHH) exhibited an average ΔG^int^ of -300.5 kcal mol^-1^. These prediction analyses suggest that the eukaryotic SAHH tetramer assembly displays more favorable solvation free energy than the archaeal SAHH tetramer assembly. Regarding the stability of archaeal SAHH tetramers, the shortened C-terminal domain was speculated to either be useful for intramolecular motion at high temperature and prevent denaturation based on a protein model for *Pfu*SAHH^36^, or to positively affect the thermostability of the enzyme^27^. Nevertheless, our structures of different archaeal SAHHs clearly show that they form stable tetramers. Instead of interactions between C-terminal domains of neighbouring monomers, the major forces for stabilisation are provided by aromatic and hydrophobic amino acid residues at the interfaces of the tetramers^36^.

**Figure 7:**
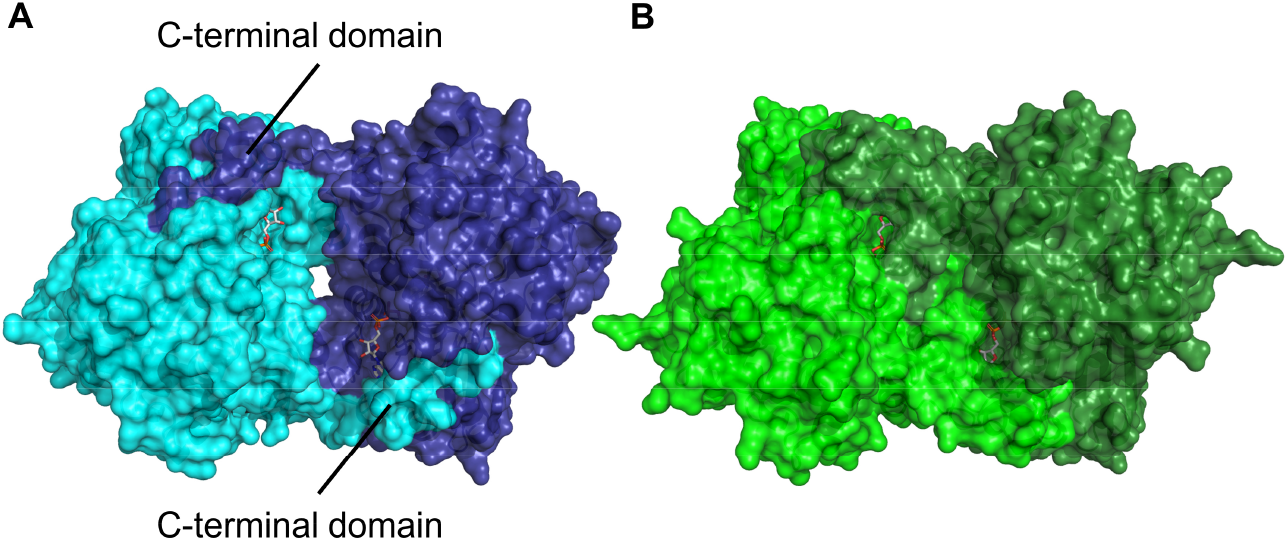
Overall structural analysis of *Mm*SAHH and *Pfu*SAHH, an euryarchaeotic SAHH, regarding the cofactor-binding domain. A: In *Pfu*SAHH, the cofactors are loosely covered by the adjacent monomer while some room is visible between the monomers, indicating a less stable enzyme (PDB ID: 7R38). B: In *Mm*SAHH, the cofactors are tightly covered by the C-terminus of the adjacent monomer (PDB ID: 5AXA).

### Conclusion

In this work, we present the biochemical characterisation of various archaeal, bacterial, and eukaryotic SAHH/SIHH homologues revealing different substrate preferences. While enzymes deriving from the Crenarchaeota phylum (*Sac*SAHH and *Sso*SAHH) or from *C. glutamicum* show catalytic activity only with SAH, the other tested SAHHs/SIHHs accept SAH as well as SIH as substrates. The homologues from *T. maritima* and the euryarchaeal classes of *Methanococci* and *Thermococci* (*Mj*SIHH, *Pfu*SAHH, *Tk*SAHH, and *Mma*SAHH) show higher catalytic activity with SIH. In accordance with the substrate preference of tested SAHHs/SIHHs is the presence of the SAH deaminating enzyme DadD in the organisms employing enzymes with a preference for SIH. Bioinformatical alignment of the amino acid sequences revealed that the SAHH/SIHH homologues with a preference for SIH contain a HxExK motif (as described before for selected extremophile enzymes) for binding of the nucleobase moiety in their active site while the homologues preferring SAH contain a HxTxE(Q) motif instead. In addition to the biochemical investigations, we provide the first structural insights into SAHHs/SIHHs originating from archaeal organisms. The archaeal enzymes show the same binding modes for the cofactor NAD^+^ as well as for the substrates inosine/adenosine and SIH/SAH as homologues from other domains of life. The shortened C-terminal domain of archaeal SAHHs/SIHHs has an influence on their tertiary and quaternary arrangement showing differences compared to bacterial and eukaryotic homologues. The observation of less interactions between archaeal SAHH/SIHH monomers leading to the cofactor NAD^+^ being more exposed to the solvent suggests a less stable homotetrameric assembly for archaeal SAHHs/SIHHs. However, the archaeal enzymes tested in this study form stable homotetramers in which the lower number of interactions could promote thermostability and intramolecular motion at higher temperatures. In summary, our functional and structural knowledge of SAHHs/SIHHs strongly promotes the existence of an alternate route for methyl metabolism and purine salvage, which might run in parallel with the canonical pathway, as SAH is also accepted as a substrate.

## Methods

### Materials

Substrates and reference standards were purchased in the highest purity available from Sigma-Aldrich (adenosine, Hcy, hypoxanthine, inosine, SAH, *S*-methyl-l-methionine), and AppliChem (adenine). Ingredients for buffers and cultivation media were purchased from Carl Roth.

### Cloning, expression, and protein purification

All SAHH/SIHH genes and *Mj*DadD, except *Tk*SAHH and *Sf*SAHH, were purchased as synthetic DNA strings from Invitrogen (Thermo Fisher Scientific, Waltham, MA, USA); the exceptions were cloned directly from genomic DNA. The genes were amplified by PCR and cloned into pET28a(+) using T4 DNA ligase (New England Biolabs GmbH, Frankfurt am Main, Germany) or In-Fusion Cloning (Takara Bio Europe, Saint-Germain-en-Laye, France). All primers are listed in Supplementary Table S1 online. The l-homocysteine *S*-methyltransferase from *Saccharomyces cerevisiae* (*Sc*HSMT) was cloned, expressed and purified as previously described^44^. The enzymes were produced in *E. coli* BL21-Gold(DE3) competent cells (Agilent, Santa Clara, CA, USA). The expressions of *Tk*SAHH and *Sf*SAHH were performed in *E. coli* BL21-CodonPlus (DE3)-RIPL competent cells (Agilent, Santa Clara, CA, USA). LB medium was used for most seed cultures and main cultures, *Pfu*SAHH and *Tk*SAHH were expressed in 2xYT medium. Seed cultures (5 mL) were grown in LB medium with the corresponding antibiotics at 37 °C overnight. The main culture (400 mL plus added seed culture) was grown in medium supplemented with the needed antibiotics at 37 °C. When the OD_600_ reached 0.5, expression was induced by the addition of isopropyl-β-D-thiogalactopyranoside (IPTG; final concentration 0.25 mM, 1 mM for *Pfu*SAHH) and the cultures were shaken at 160 rpm for 18 h at 20 °C. The cells were harvested by centrifugation and lysed by sonication [Branson Sonifier 250, Emerson, St. Louis, MO, USA (duty cycle 50%, intensity 50%, 5 x 30 s with 30 s breaks)] in lysis buffer. The lysis buffer was either 50 mM Tris-HCl, pH 7.4, 500 mM NaCl, 10% (*w*/*v*) glycerol for *Mj*SIHH, *Mma*SAHH, *Pfu*SAHH, and *Sso*SAHH; or 40 mM Tris-HCl, pH 8.0, 100 mM NaCl, 10% (*w*/*v*) glycerol for *Mj*DadD, *Sac*SAHH, *Sf*SAHH, *Tk*SAHH, and *Tm*SAHH. After centrifugation, the proteins were purified by nickel-NTA affinity chromatography [lysis buffer including low concentrations (10 or 10–50 mM) imidazole for washing or high concentrations (200 or 100–300 mM) for eluting the protein], and desalted using a PD-10 column (Cytiva Europe GmbH, Freiburg im Breisgau, Germany). The storage buffer for the enzymes was the same as the lysis buffer. Protein concentration was determined using a NanoDrop 2000, at 280 nm with the molecular weight (including the His_6_-tag, Supplementary Table S1 online), and the protein extinction coefficient (calculated with the ExPASy ProtParam tool^57^).

### Crystallisation and data collection

*Pfu*SAHH with inosine (2 mM) co-crystals appeared after 2-4 days by using the sitting drop vapor diffusion methods at room temperature by combining 0.5 μL of protein at 10 mg/mL with 0.5 μL of a precipitant solution comprising 26% (*w*/*v*) PEG 1500 with 100 mM MTT, pH 8.0. *Pfu*SAHH with SIH co-crystals were obtained with a slightly modified crystallisation conditions comprising 28% (*w*/*v*) PEG 2000 with 100 mM MTT, pH 8.0. *Mma*SAHH (10 mg/mL) with inosine (2 mM) co-crystals were obtained within a week at 22 °C in sitting drops by mixing 0.5 μL of the protein solution with an equal volume of reservoir solution containing 28% (*w*/*v*) PEG 3350, 100 mM malic acid/MES/Tris-HCl buffer (MMT), pH 8.0. *Sac*SAHH (10 mg/mL) with adenosine (2 mM) co-crystals were obtained within a week at 22 °C in sitting drops by mixing 0.5 μL of the protein solution with an equal volume of reservoir solution containing 20% (*w*/*v*) PEG 3350 with 200 mM ammonium iodide. *Mm*SAHH with inosine (4 mM) co-crystals appeared after 4 days by using the sitting drop vapor diffusion methods at room temperature by combining 2 μL of protein at 4 mg/mL with 1 μL of a precipitant solution comprising 22% (*w*/*v*) PEG 3350 with 180 mM sodium formate, pH 6.9. Prior to data collection, the crystals were transferred to a cryosolution containing the respective mother liquor reservoir solutions and flash frozen in liquid nitrogen. Datasets were collected at 100 K at the Swiss Light Source (SLS) on macromolecular crystallography beamline PXI-X06SA. All *Pfu*SAHH crystals were maintained at a constant temperature (100 K) and a total of 900 images (Δφ = 0.2°/image) for inosine complex, and 1800 images (Δφ = 0.2°/image) for the SIH complex were recorded separately for each on an EIGER 16M (Dectris) detector. The datasets were extending up to 2.3 Å resolution for inosine complex, and up to 2.0 Å for the SIH complex. All datasets were processed by XDS^58^ in *P*4_2_2_1_2 space group (a = b = 111.68, c = 121.59; α = β = γ = 90°). *Mma*SAHH crystals were flash-cooled and maintained at a constant temperature at 100 K in a cold nitrogen-gas stream. A dataset with a total of 900 images (Δφ = 0.2°/image) were recorded on an EIGER 16M (Dectris) detector. The datasets were extending up to 2.5 Å resolution in *P*2_1_ space group (a = 65.76, b = 328.9, c = 82.05; α = γ = 90°, β = 107.2°). *Sac*SAHH crystals were maintained at a constant temperature (100 K) and a total of 3600 images (Δφ = 0.1°/image) were recorded on a EIGER 16M (Dectris) detector with data extending up to 2.6 Å resolution. The datasets were processed in *P*1 space group (a = 84.36, b = 88.36, c = 138.54; α = 78.86°, β = 74.85°, γ = 64.85°). *Mm*SAHH crystals were maintained at a constant temperature (100 K) and a total of 900 images (Δφ = 0.1°/ image^−1^) were recorded on an EIGER 16M (Dectris) detector with data extending up to 2.48 Å resolution. The datasets were processed by XDS^58^ in *I*222 space group (a = 97.78, b = 101.96, c = 172.58; α = β = γ = 90°). All the data were integrated by using XDS^58^ then merged and scaled using SCALA from the CCP4 suite of programs^59,60^. The data collection statistics are summarised in Supplementary Table S3 online.

### Structure determination and refinement

Initial phases were determined by molecular replacement using PHASER. Best solutions were obtained using 5AXA based homology model as the starting model for *Pfu*SAHH, 1V8B for *Mma*SAHH and 3H9U for *Sac*SAHH. The asymmetric unit (ASU) of *Pfu*SAHH crystals contained a dimer, whereas *Mma*SAHH comprised four dimers and SacSAHH comprised two dimers. The models were built with COOT^60^, and refinements were carried out with REFMAC using NCS constraints with or without TLS parameters^61^. The NAD^+^ ligands in all the complex crystals were clearly observed in the initial 2Fo-Fc and Fo-Fc maps. To improve the model of the bound substrate and product, we calculated both 2Fo-Fc maps and POLDER omit maps^62^ and fitted the ligands into the respective electron densities, which allowed unambiguous identification of the ligand positioning (Supplementary Fig. S19F online). Incorporation of non-crystallographic symmetry (NCS) restraints greatly expedited model improvement. For *Pfu*SAHH•NAD•inosine complex, a Ramachandran plot calculation indicated that 97% and 3% of the residues occupy the most favored and additionally allowed regions, respectively. For *Pfu*SAHH•NAD•SIH complex, a Ramachandran plot calculation indicated that 96% and 4% of the residues occupy the most favored and additionally allowed regions, respectively. For *Mma*SAHH•NAD•inosine complex, a Ramachandran plot calculation indicated that 96% and 3% of the residues occupy the most favored and additionally allowed regions, respectively. For *Sac*SAHH•NAD•adenosine complex, a Ramachandran plot calculation indicated that 94% and 5% of the residues occupy the most favored and additionally allowed regions, respectively. For *Mm*SAHH•NAD•inosine complex, a Ramachandran plot calculation indicated that 97% and 3% of the residues occupy the most favored and additionally allowed regions, respectively. Analysis of the SAHH structures and comparison with other SAHH structures were carried out using PyMOL (http://pymol.sourceforge.net/, the PyMOL Molecular Graphics System, Version 2.5.0a0 Schrödinger, LLC)^58^.

### NMR spectroscopy

Nuclear magnetic resonance (NMR) spectra were recorded on an Avance DRX 400 spectrometer (Bruker, Billerica, MA, USA), operating at 400.1 MHz (for ^1^H NMR) and 100.6 MHz (for ^13^C NMR). All measurements were performed at 25 °C. Spectra were analysed with TopSpin 3.6.2.

### HPLC analysis

All available substrates and products of the enzymatic reactions are used as authentic reference standards and the retention times are listed in Supplementary Table S2 online. All assays were analysed with an Agilent 1100 Series HPLC using an ISAspher SCX 100-5 column (250 mm x 4.6 mm, 5 μm; ISERA GmbH, Düren, Germany). The HPLC method has been described previously, using 40 mM sodium acetate, pH 4.2, as buffer A, and acetonitrile as buffer B^63^. The injection volume was set to 5 μL.

### SIH enzymatic synthesis and structure verification

SIH was enzymatically synthesised with a 5’-deoxyadenosine deaminase (*Mj*DadD) starting from SAH following a published protocol with modifications^13,14^. 1 mM SAH was incubated with 1 μM *Mj*DadD in 50 mM Tris-HCl buffer, pH 8.0, for 20 h at 37 °C. The enzyme was removed with a spin filter and full conversion checked with HPLC-DAD. The stock solution of SIH (1 mM) was stored at −20 °C. The structure was verified by NMR analysis by running six parallel assays with 4 mL reaction tube and combining them in one 100 mL flask. The solution was freeze-dried, and the powder was resuspended in 700 μL D_2_O and measured with NMR spectroscopy. SIH has been isolated from *Streptomyces flocculus* (*Streptomyces albus* ATCC 13257) previously and the structure was confirmed by UV spectrum and ^1^H NMR analysis^16^. An HPLC method was used to track the conversion from SAH to SIH catalysed by *Mj*DadD (Supplementary Fig. S1 online), here the UV spectra matched the described absorbance maximum shift from 260 nm (SAH) to 248 nm (SIH; Supplementary Fig. S1 online). ^1^H NMR data (Supplementary Fig. S2B online) obtained matched the published data and was extended by measuring ^13^C spectrum (Supplementary Fig. S2C online), as well as 2D spectra (Supplementary Figs. S2D-F online) to further confirm the SIH structure.

### Enzyme activity assays

All assays concerning SAHHs/SIHHs were performed in 50 mM Tris-HCl, pH 7.5, for 20 h at 37 °C in 200 μL reaction volume, if not otherwise stated. For the synthesis reaction, 0.5 mM Hcy and 0.5 mM nucleoside were added, while the hydrolysis reaction was started with 0.2 mM of either SAH or SIH. The SAHH/SIHH was added at 5 μM. A second enzyme, *Sc*HSMT (10 μM), and its methyl donor *S*-Methyl-l-methionine (1 mM) were added to the SAH or SIH hydrolysis to drive the reaction forward. After incubation, 150 μL of the assay sample was added to 50 μL of perchloric acid [final concentration 2.5% (*w*/*v*)] to stop the reaction and spun down for 30 min prior transferring 80 μL to an HPLC vial. All investigated SAHHs and SIHHs were tested in the SAH and SIH hydrolysis, as well as synthesis direction. An overview can be found in Table 1, while the HPLC chromatograms are given in Supplementary Figs. S4–S16 and the SDS gels in Supplementary Fig. S3 online.

### Bioinformatical analysis of amino acid sequences

Alignments of amino acid sequences were performed either with Clustal Omega or T-Coffee online services^64,65^. For the visualisation of the phylogenetic tree, the MEGA11 software was used^66^. To visualise and annotate the sequence alignment, Jalview Version2 was used^67^.

## Acknowledgments

This work was supported by the Deutsche Forschungsgemeinschaft (DFG, German Research Foundation) – 235777276/RTG1976 and the European Research Council (ERC project 716966). We thank Jun.-Prof. Dr Silja Mordhorst (now University of Tübingen) for construction of some plasmids used in this work. We thank Emina Čokljat and Katharina Strack for assistance with protein production and purification. We thank Dr Tomizaki Takashi and the PXI (X06SA) beamline staff of the Swiss Light Source, Paul Scherrer Institute (Villigen, Switzerland) for support in crystallographic data collection. We thank Sascha Ferlaino and Dr Phillippe Bisel (both University of Freiburg) for NMR measurements and interpretation of the results, respectively. Prof. Dr Sonja-Verena Albers (University of Freiburg) is acknowledged for sharing her expertise regarding archaea and for critically reading the manuscript; as well as Prof. Dr Michael Müller (University of Freiburg) for helpful discussions on the binding modes and mechanisms.

## Author Contributions

D.P.: conceptualisation, methodology, investigation, writing - original draft preparation, writing - review and editing, and visualisation; R.S.-B.: conceptualisation, methodology, investigation, writing - original draft preparation, writing - review and editing, and visualisation; L.-H.K.: methodology, investigation; writing - review and editing, and visualisation; P.G.: data interpretation, and writing - review and editing; J.N.A.: conceptualisation, writing - review and editing, supervision, project administration, resources, and funding acquisition.

## Additional Information

**Supplementary information** is available online

