## Supplemental Material for "Archaeal *S*-adenosyl-l-homocysteine hydrolases: structure, function and substrate preferences"

Electronic Supplementary Information

### Experimental

**Table S1.** The primers used for cloning of SAHHs/SIHHs and their molecular weight including $His_6$-tag.

| Enzyme | Forward primer (5'–3') | Reverse primer (5'–3') | Molecular weight [kDa] | Reference |
|---|---|---|---|---|
| *Cg*SAHH | TATATATACATATGGCACAG GTTATGGACTTC | TATATATACTCGAGTTAGTAG CGGTAGTGCTC | 54.59 | This work |
| *Ll*SAHH | TATATATACATATGGCCCTGC TGGTTG | TATATATAAAGCTTTTAGTAG CGATAATGAAACG | 55.49 | Guranowski and Pawelkiewicz, 1977[1]; Brzezinski *et al.,* 2008[2] and 2012[3] |
| *Mj*DadD | TATATATACATATGATCCTGA TCAAAAACG | TATATATACTCGAGTTAGCTG CGCAGAATT | 49.63 | Miller *et al.*, 2014[4] |
| *Mj*SIHH | TATATATACATATGTACGAGG TGCGC | TATATATACTCGAGTTAGGTG CCTTC | 48.77 | Miller *et al.*, 2015[5] |
| *Mma*SAHH | CGCGCGGCAGCCATATGAGC AACGTGAAAGATATGAGCC | GTGCGGCCGCAAGCTTAGGT GCCTTCTTTCCAATCGC | 48.00 | This work |
| *Mm*SAHH | CGCGCGGCAGCCATATGAGC GACAAACTGCCGTATAAAG | GTGCGGCCGCAAGCTAAGCT TAATAGCGATAATGATCCGG | 49.85 | Ishihara *et al.*, 2010[6]; Kusakabe *et al.*, 2015[7] |
| *Pa*SAHH | TATATATACATATGAGCGCA GTTATGACAC | TATATATAAAGCTTTTAATAG CGATAGGTATCC | 53.56 | Czyrko *et al.,* 2018[8] |
| *Pfu*SAHH | CGCGCGGCAGCCATATGGAT TGCGGCAAAGATTATTG | GTGCGGCCGCAAGCTTAGGT GCCATGTTCCCAAC | 49.55 | Porcelli *et al.*, 2005[9] |
| *Sac*SAHH | TATACATATGGATTATCGCGT TAAAGATCTG | TATATAAGCTTAGGTGCCGTA TTTCCACTG | 48.34 | This work |
| *Sa*SAHH | TATATATACATATGGCAAGC GCCCAGCA | TATATATAAGCTTCAGTACCG GTAGTGGTC | 54.72 | This work |
| *Sf*SAHH | TATACATATGACGACGACCTC CACGAC | TATAAGCTTTCAGTAGCGGTA GTGG | 54.87 | This work |
| *Sso*SAHH | CGCGCGGCAGCCATATGAGC TACAAAATCAAAGATCTGAG | GTGCGGCCGCAAGCTTAGGT GCCGCTTTTCCACTG | 48.13 | Porcelli *et al.*, 1993[10] and 2000[11] |
| *Tk*SAHH | CGCGCGGCAGCCATATGGAC TGCACGAAGGATT | GTGCGGCCGCAAGCTAAGCT TCAGGTGCCGTGCTCCCAGC | 49.04 | This work |
| *Tm*SAHH | TATATATACATATGAACACCG GTGAGA | TATATATACTCGAGTTACTGC CAGCTA | 47.01 | Hermann *et al.*, 2007[12]; Lozada-Ramírez *et al.*, 2013[13] |

### HPLC analysis

**Table S2.** HPLC retention times.

| Substance | Retention time [min] |
|---|---:|
| Adenine | 9.2 |
| Adenosine | 7.9 |
| Hypoxanthine | 3.4 |
| Inosine | 2.7 |
| SAH | 8.0 |
| SIH | 3.0 |

### Crystallography

**Table S3.** Data collection and refinement statistics (Molecular Replacement).

| | ***Mma*SAHH• NAD•inosine** | ***Pfu*SAHH• NAD•inosine** | ***Pfu*SAHH• NAD•SIH** | ***Sac*SAHH• NAD•adenosine** | ***Mm*SAHH• NAD•inosine** |
|---|---|---|---|---|---|
| PDB ID | 7R3A | 7R37 | 7R38 | 7R39 | 8COD |
| **Data collection** | | | | | |
| Space group | $P2_1$ | $P4_22_12$ | $P4_22_12$ | $P1$ | $I222$ |
| Cell dimensions | | | | | |
| a, b, c [Å] | 65.8, 328.9, 85.1 | 112.5, 112.5, 122.8 | 111.7, 111.7, 121.6 | 84.4, 88.4, 138.6 | 98.2, 102.5, 173.4 |
| α, β, γ [°] | 90, 107.2, 90 | 90, 90, 90 | 90, 90, 90 | 78.9, 74.9, 64.9 | 90, 90, 90 |
| Resolution [Å] | 45.44–2.65 (2.74–2.65) | 48.62–2.28 (2.37–2.28) | 53.4–2.05 (2.12–2.05) | 46.16–2.50 (2.59–2.50) | 54.9–2.48 (2.56–2.48) |
| $R_{merge}$ | 0.1748 (1.075) | 0.1495 (1.191) | 0.1048 (0.9204) | 0.1348 (0.582) | 0.177 (0.97) |
| $I/\sigma I$ | 6.73 (2.13) | 13.51 (1.66) | 16.82 (2.70) | 6.67 (2.35) | 7.8 (2.1) |
| Completeness [%] | 97.3 (98.2) | 99.8 (98.8) | 99.7 (99.9) | 87.5 (85.9) | 98.8 (99) |
| Redundancy | 3.6 (3.5) | 12.7 (8.3) | 12.8 (13.4) | 3.6 (3.6) | 5.7 (5.9) |
| **Refinement** | | | | | |
| Resolution [Å] | 45.44–2.65 (2.74–2.65) | 48.62–2.28 (2.37–2.28) | 53.4–2.05 (2.12–2.05) | 46.16–2.50 (2.59–2.50) | 54.9–2.48 (2.56–2.48) |
| No. reflections | 141280 (13525) | 36333 (3519) | 48676 (4780) | 105739 (10341) | 31048 (3056) |
| $R_{work}$ /$R_{free}$ [%] | 18.8/24.3 | 16.1/21.6 | 17.6/21.9 | 22.9/28.2 | 17.0/21.8 |
| No. atoms | 26322 | 6891 | 7147 | 27556 | 7067 |
| Protein | 25440 | 6604 | 6706 | 25760 | 6650 |
| Ligand/ion | 608 | 126 | 88 | 504 | 128 |
| Water | 274 | 161 | 353 | 1292 | 289 |
| *B*-factors | 67.14 | 46.77 | 38.41 | 40.96 | 40.98 |

| | | | | | |
|---|---|---|---|---|---|
| Protein | 67.59 | 46.98 | 38.16 | 40.23 | 41.16 |
| Ligand/ion | 55.84 | 38.49 | 34.81 | 31.83 | 35.81 |
| Water | 50.24 | 44.84 | 44.11 | 59.02 | 39.23 |
| R.m.s. deviations | | | | | |
| Bond lengths [Å] | 0.030 | 0.014 | 0.014 | 0.008 | 0.014 |
| Bond angles [°] | 2.33 | 1.81 | 1.84 | 1.69 | 1.86 |

Each structure was solved from a single crystal. Values in parentheses are for highest-resolution shell.

### SIH enzymatic synthesis and structure verification

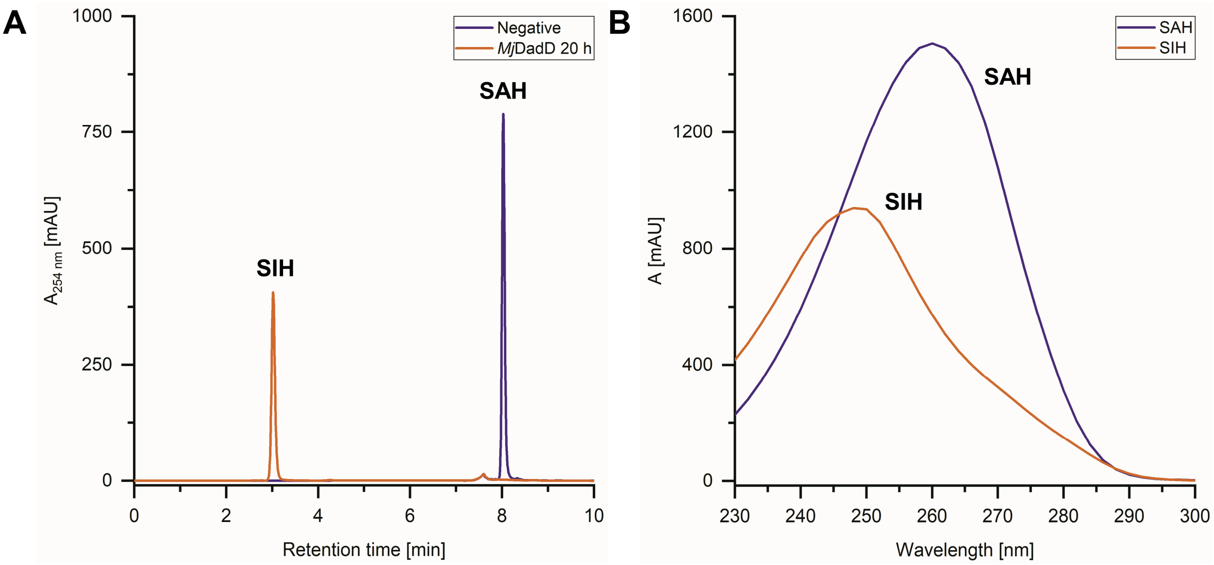

**Figure S1. A.** HPLC chromatogram showing the conversion of SAH (purple; 8.0 min) to SIH (orange; 3.0 min) catalysed by *Mj*DadD. **B.** UV spectrum of SAH and SIH showing the absorbance maximum shift between SAH (purple; 260 nm) and SIH (orange; 248 nm).

DMSO-d6 was initially used to solve the compound and the solvent signals are still visible in the NMR spectra [$^{13}$C NMR: DMSO-d6 38.5 ppm (septet, $CD_3$)]; ultimately $D_2O$ was used for dissolving the powder after lyophilisation. Signals for the enzyme storage buffer are also visible in the NMR spectra [$^{1}$H NMR of glycerol and Tris signals: δ = 3.24 (dd for 2 H, glycerol), 3.34 (dd for 2 H, glycerol), 3.44-3.51 (m, 1 H, C-H in glycerol, also overlapping with Tris signal) $^{13}$C NMR: δ = 72.0 (C-H in glycerol), 62.4 (C-$H_2$ in glycerol), 60.6 (C-$H_2$ in Tris); smaller signal for quaternary C in Tris not visible at this concentration].

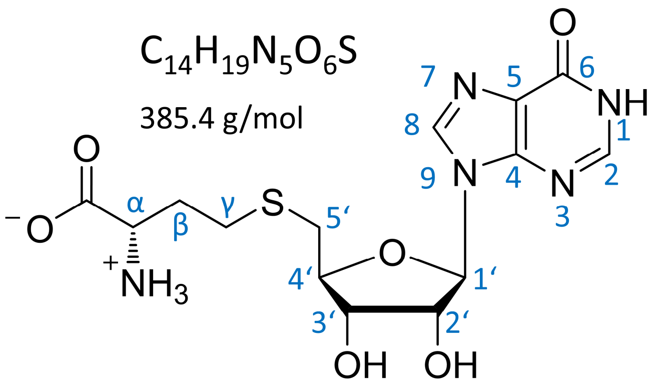

**SIH**

**Figure S2A.** Structure, molecular formula and molecular weight of SIH. Carbon atoms are numbered according to the assignment of the NMR signals.

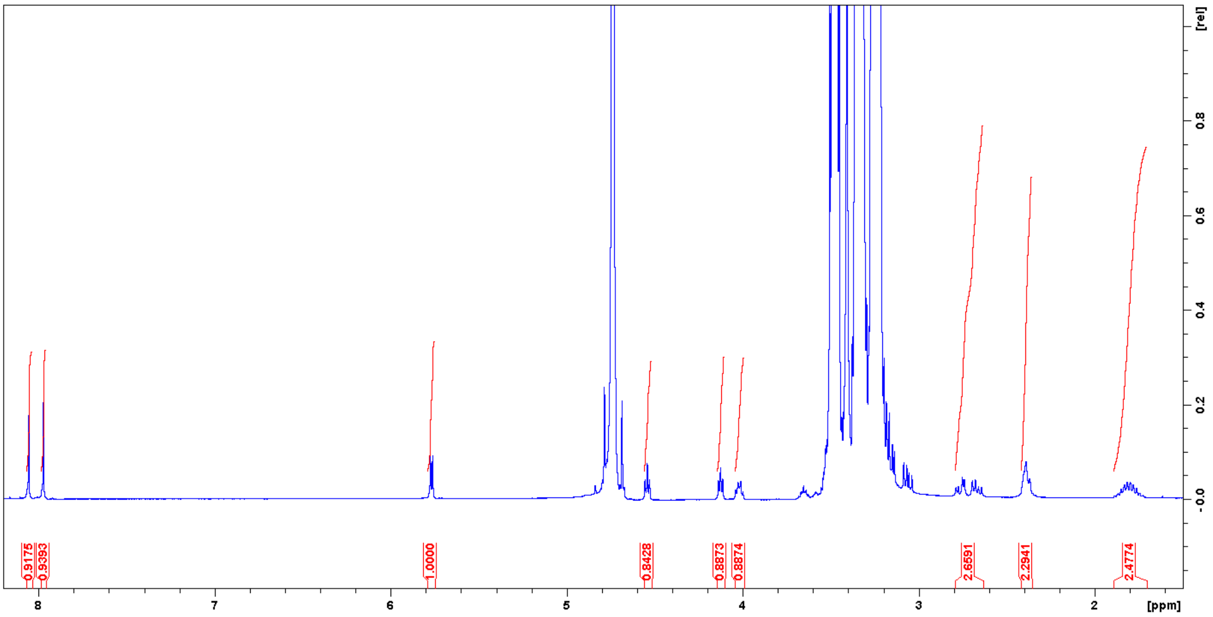

**Figure S2B.** $^{1}$H NMR spectrum of enzymatically synthesised SIH.

$^{1}$H NMR ($D_2O$): δ = 8.06 (s, 1 H, H-8), 7.98 (s, 1 H, H-2), 5.77 (d, *J* = 5.06 Hz, 1 H, H-1'), 4.55 (t, J = 5.06 Hz, 1 H, H-2'), 4.13 (t, *J* = 5.06 Hz, 1 H, H-3'), 4.02 (dt, J = 4.87, 6.38 Hz, 1 H, H-4'), 3.52-3.55 (m, 1 H, $H_\alpha$), 2.66 (dd, J = 6,98, 14.20 Hz, 1 H, H-5'$_A$), 2.76 (dd, J = 5.06, 14.20 Hz, 1 H, H-5'$_B$), 2.34-2.44 (m, 2 H, $H_\gamma$), 1.70-1.90 (m, 2 H, $H_\beta$).

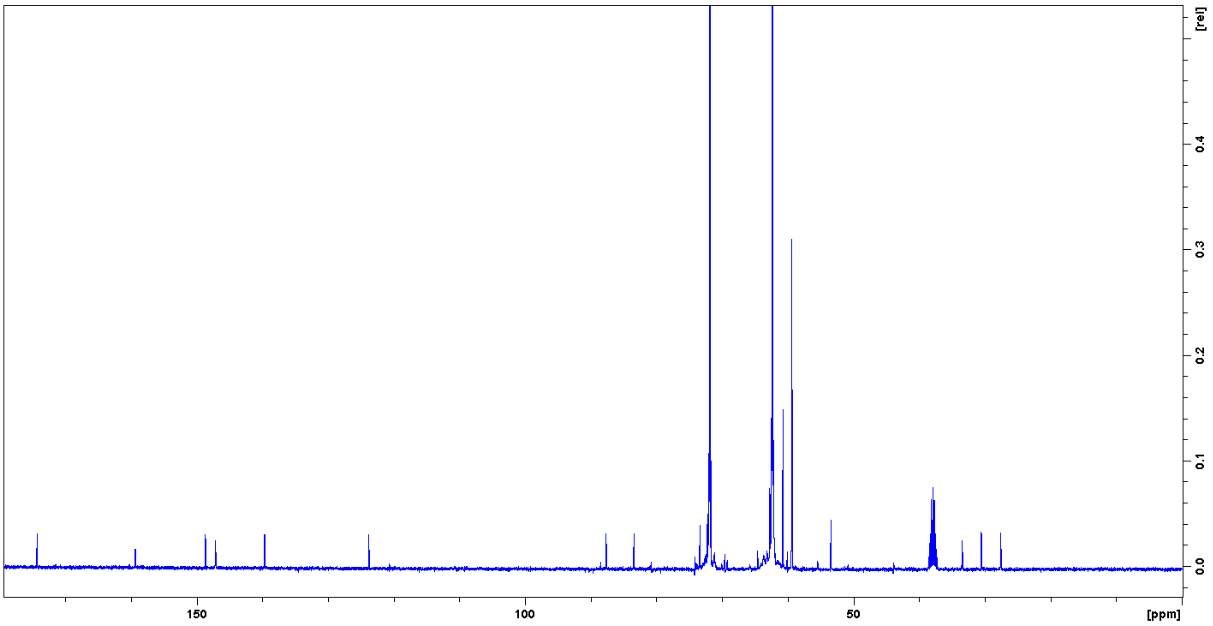

**Figure S2C.** $^{13}C$ NMR spectrum of enzymatically synthesised SIH.

$^{13}C$ NMR ($D_2O$): δ = 174.4 (COOH), 159.2 (C-6), 148.7 (C-4), 147.1 (C-2), 139.7 (C-8), 123.8 (C-5), 87.6 (C-1), 83.4 (C-4’), 73.3 (C-2’), 72.3 (C-3’), 53.4 ($C_\alpha$), 33.4 (C-5’), 30.4 ($C_\beta$), 27.6 ($C_\gamma$).

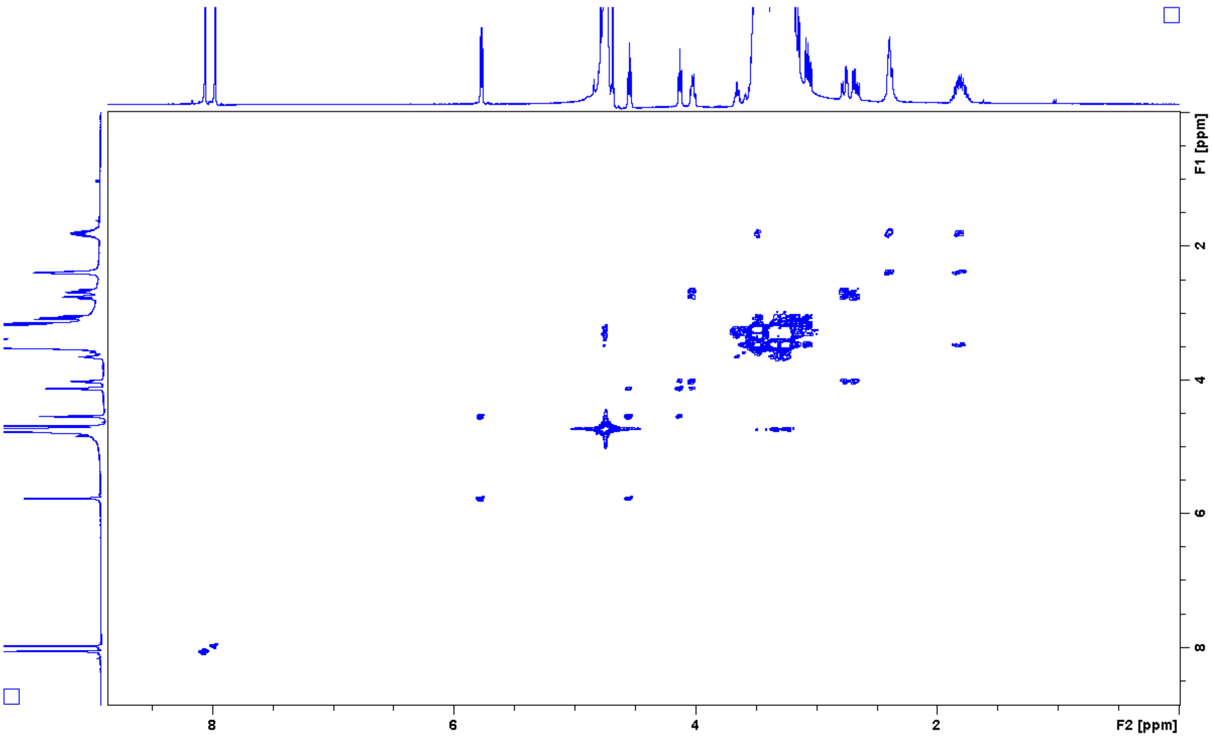

**Figure S2D.** H,H-COSY spectrum of enzymatically synthesised SIH.

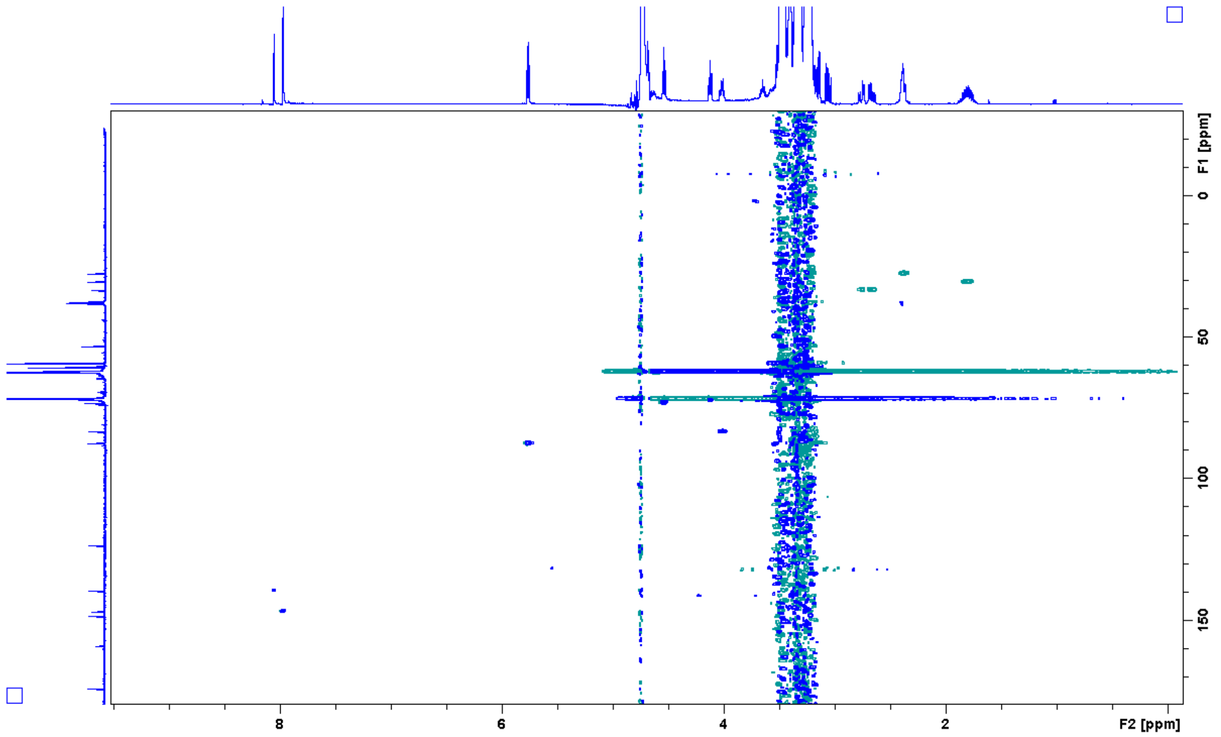

**Figure S2E.** HSQC spectrum of enzymatically synthesised SIH.

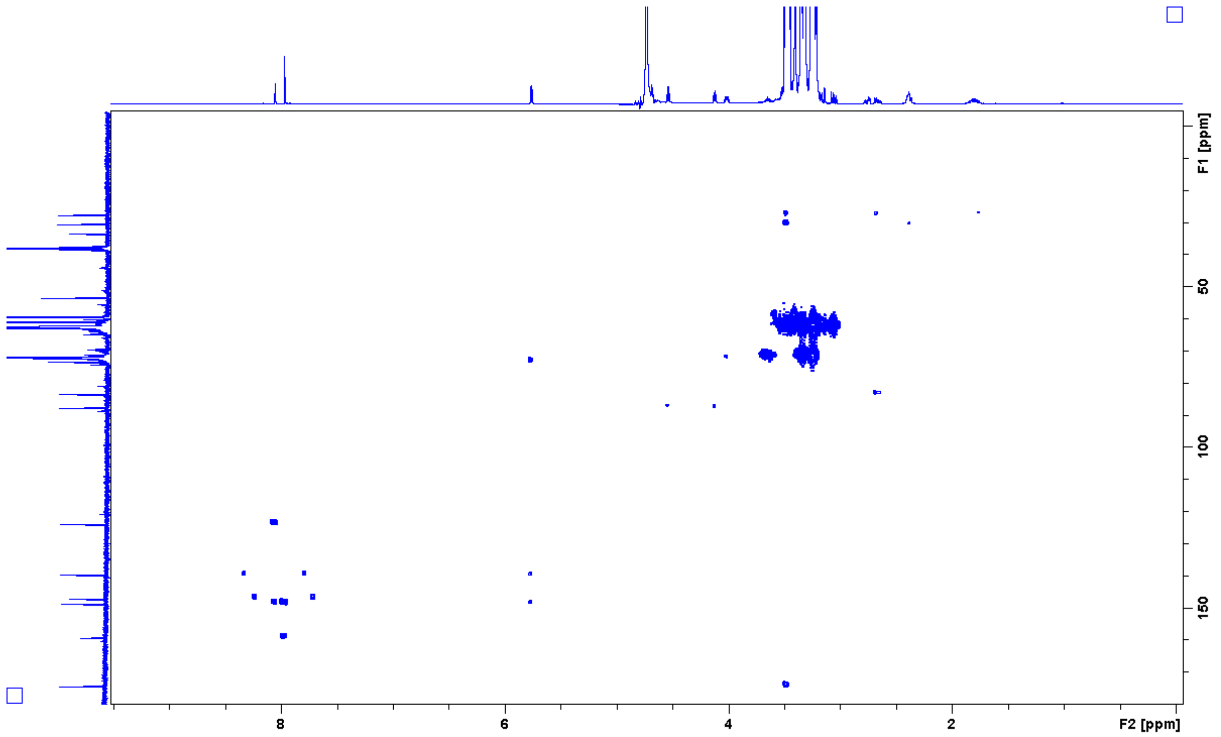

**Figure S2F.** HMBC spectrum of enzymatically synthesised SIH.

### SAHH/SIHH characterisation

**Table S4.** Enzymes used in this study with their UniProt accession numbers [1–3,5–13].

| Enzyme | Organism | Domain | Phylum/Kingdom | UniProt accession number |
|---|---|---|---|---|
| *Sac*SAHH | *Sulfolobus acidocaldarius* | Archaea | Crenarchaeota | Q4JAZ7 (SAHH_SULAC) |
| *Sso*SAHH | *Saccharolobus solfataricus* | Archaea | Crenarchaeota | P50252 (SAHH_SACS2) |
| *Mj*SIHH | *Methanocaldococcus jannaschii* | Archaea | Euryarchaeota | Q58783 (SIHH_METJA) |
| *Mma*SAHH | *Methanococcus maripaludis* | Archaea | Euryarchaeota | Q6LYR8 (SIHH_METMP) |
| *Pfu*SAHH | *Pyrococcus furiosus* | Archaea | Euryarchaeota | P50251 (SAHH_PYRFU) |
| *Tk*SAHH | *Thermococcus kodakarensis* | Archaea | Euryarchaeota | Q5JED2 (SAHH_THEKO) |
| *Cg*SAHH | *Corynebacterium glutamicum* | Bacteria | Actinomycetota | Q8NSC4 (SAHH_CORGL) |
| *Pa*SAHH | *Pseudomonas aeruginosa* | Bacteria | Pseudomonadota | Q9I685 (SAHH_PSEAE) |
| *Sa*SAHH | *Streptomyces albus* | Bacteria | Actinomycetota | Not assigned |
| *Sf*SAHH | *Streptomyces flocculus* | Bacteria | Actinomycetota | Not assigned |
| *Tm*SAHH | *Thermotoga maritima* | Bacteria | Thermotogota | O51933 (SAHH_THEMA) |
| *Ll*SAHH | *Lupinus luteus* | Eukaryota | Plantae | Q9SP37 (SAHH_LUPLU) |
| *Mm*SAHH | *Mus musculus* | Eukaryota | Animalia | P50247 (SAHH_MOUSE) |

The protein standard featuring pink bands is the Precision Plus Protein Dual Color Standard by Bio-Rad Laboratories GmbH (Feldkirchen, Germany; taken from https://www.bio-rad.com/en-de/sku/1610374-precision-plus-protein-dual-color-standards-500-ul?ID=1610374), while the one with green and orange bands is the Color Prestained Protein Standard, Broad Range (11–245 kDa; taken from https://international.neb.com/products/p7712-color-prestained-protein-standard-broad-range-11-245-kda) manufactured by New England Biolabs GmbH (Frankfurt am Main, Germany). The standard with only blue bands is the Blue Prestained Protein Standard, Broad Range (11–190 kDa; New England Biolabs GmbH, Frankfurt am Main, Germany taken from https://international.neb.com/products/p7706-blue-prestained-protein-standard-broad-range-11-190-kda).

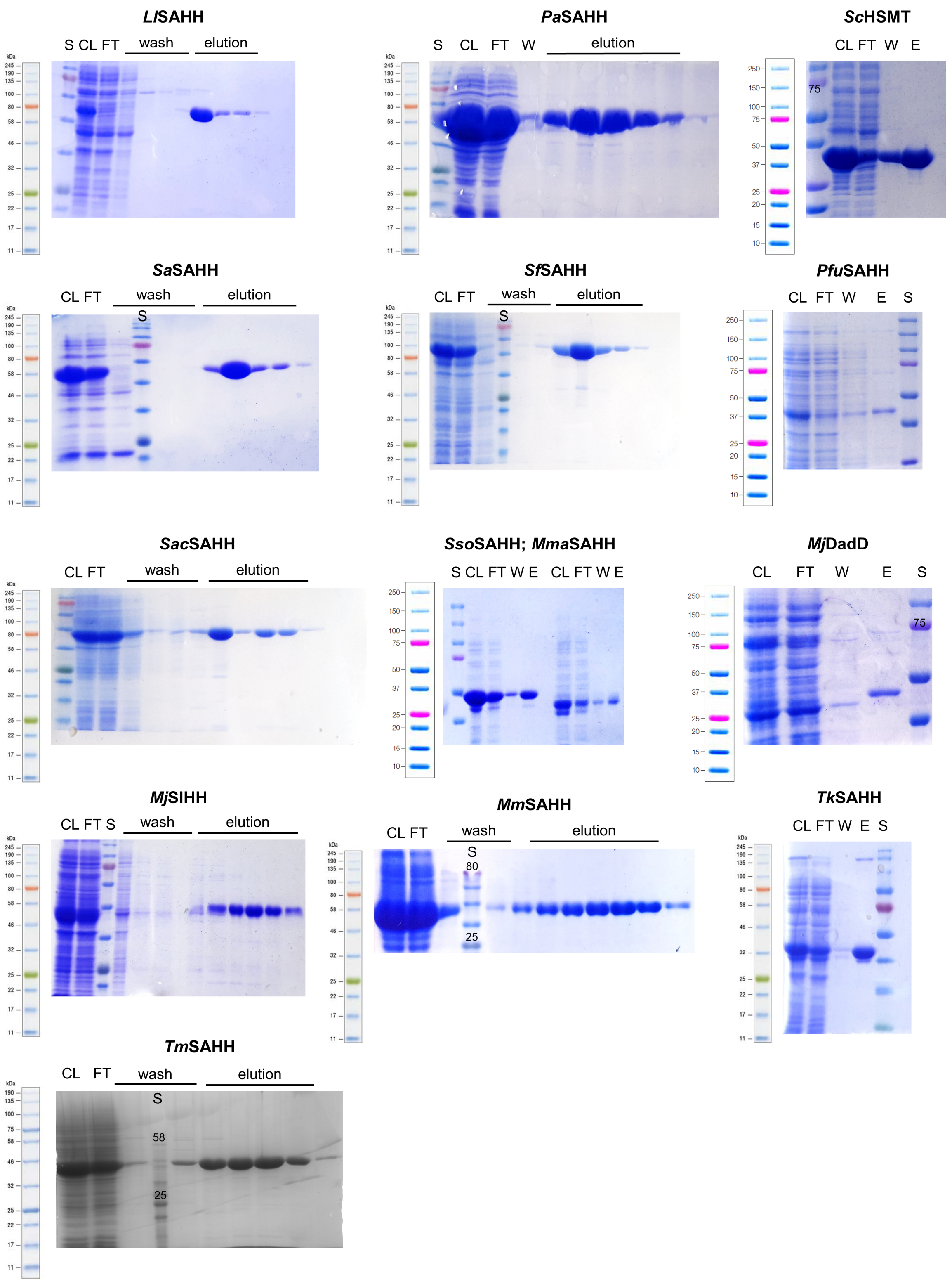

**Figure S3. SDS gels of all purifications of enzymes used in this work.**[14] CL, crude lysate; E, elution; FT, flow through; S, standard protein ladder; W, wash. Gels were cropped if other samples were not relevant.

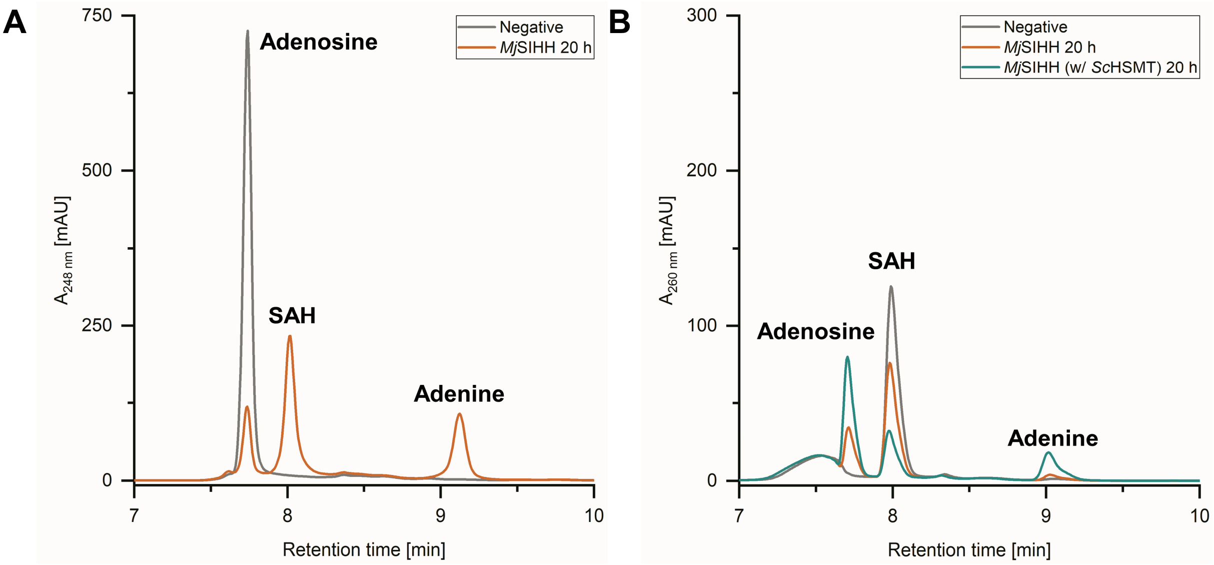

**Figure S4A.** HPLC chromatograms showing **A.** the SAH synthesis reaction and **B.** the SAH hydrolysis reaction (with and without the addition of *Sc*HSMT) catalysed by *Mj*SIHH.

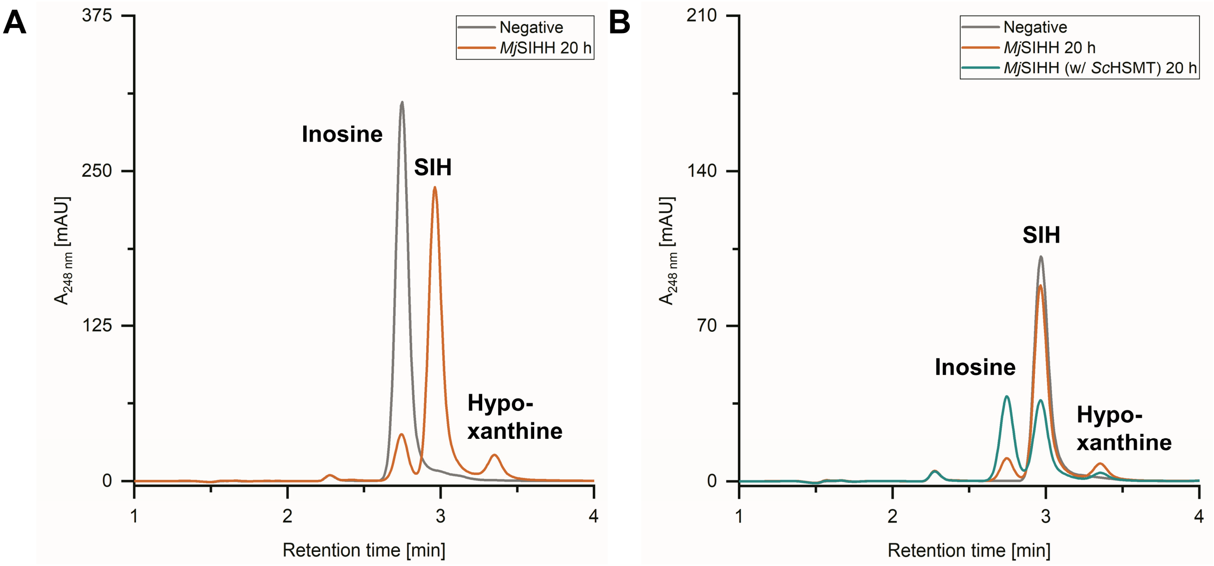

**Figure S4B.** HPLC chromatograms showing **A.** the SIH synthesis reaction and **B.** the SIH hydrolysis reaction (with and without the addition of *Sc*HSMT) catalysed by *Mj*SIHH.

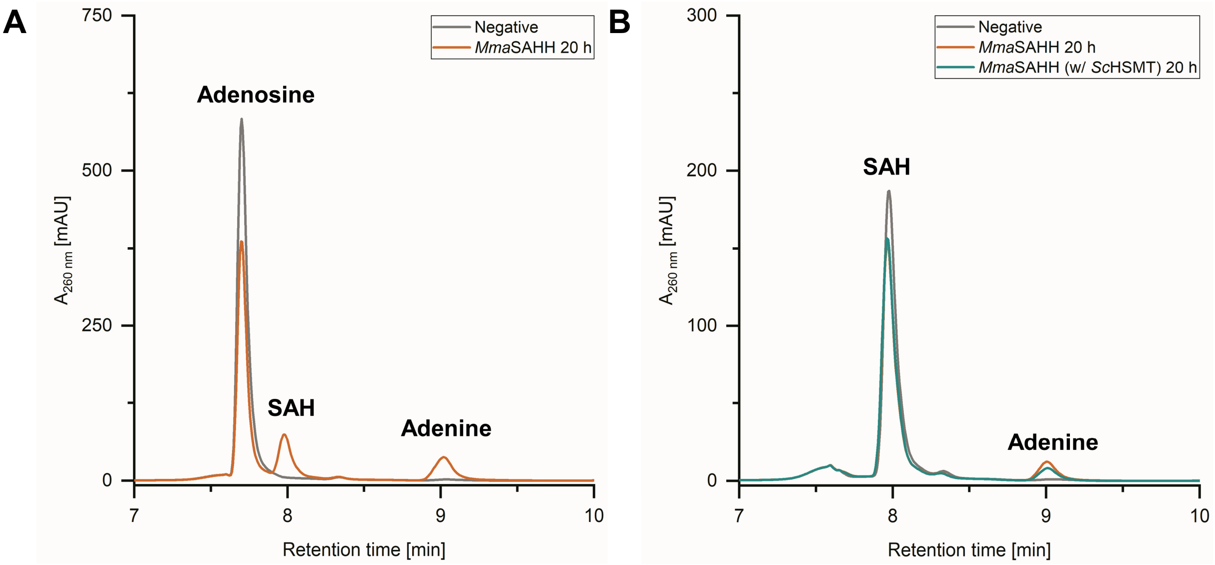

**Figure S5A.** HPLC chromatograms showing **A.** the SAH synthesis reaction and **B.** the SAH hydrolysis reaction (with and without the addition of *Sc*HSMT) catalysed by *Mma*SAHH.

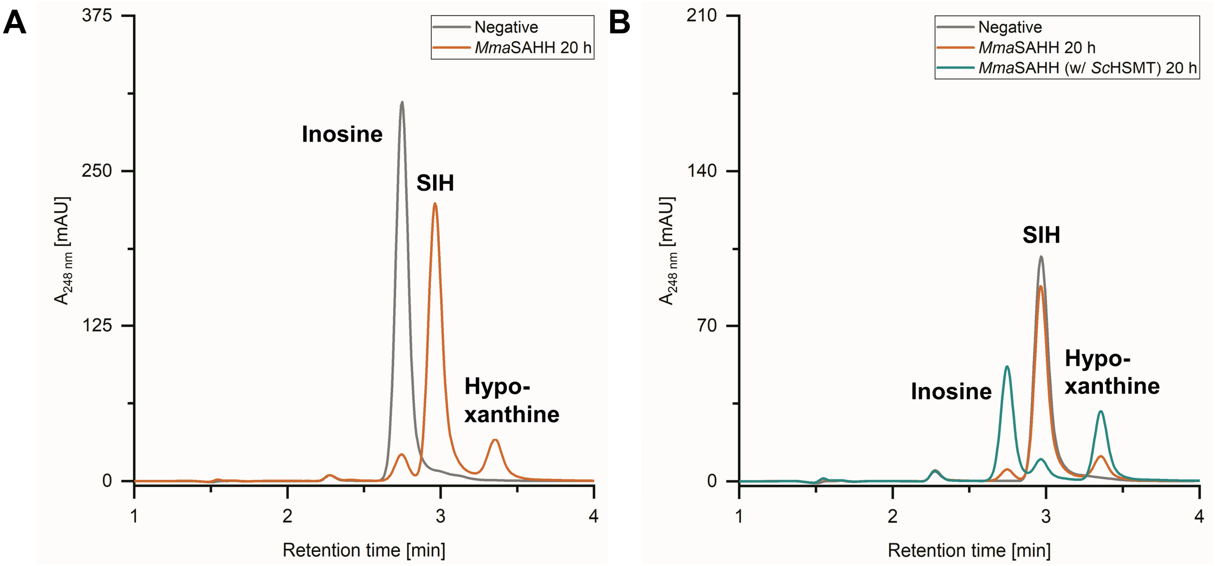

**Figure S5B.** HPLC chromatograms showing **A.** the SIH synthesis reaction and **B.** the SIH hydrolysis reaction (with and without the addition of *Sc*HSMT) catalysed by *Mma*SAHH.

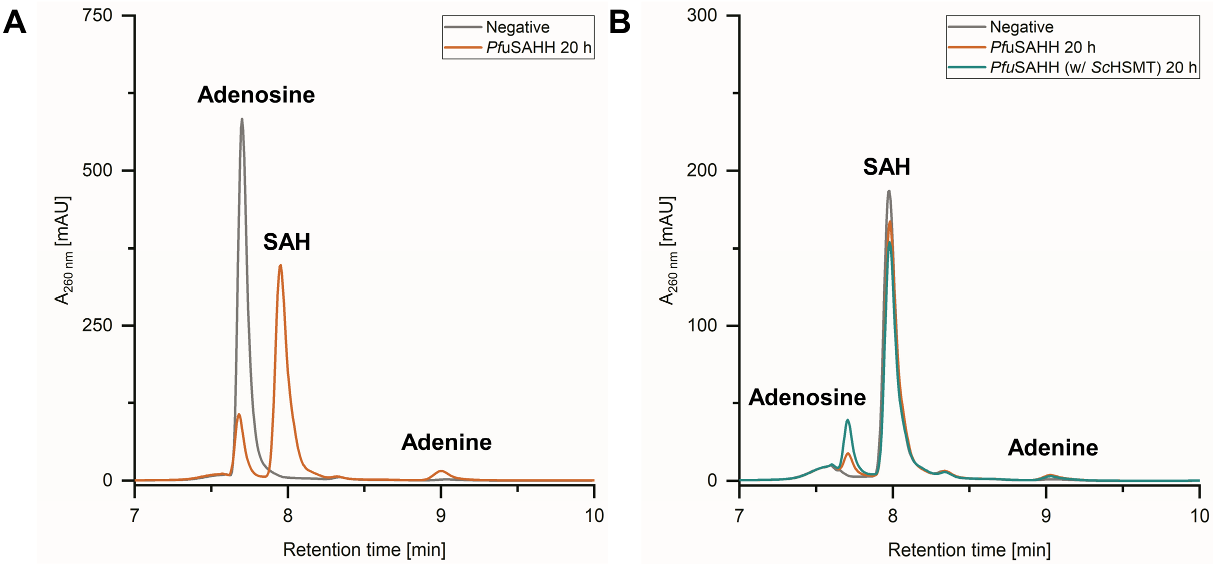

**Figure S6A.** HPLC chromatograms showing **A.** the SAH synthesis reaction and **B.** the SAH hydrolysis reaction (with and without the addition of *Sc*HSMT) catalysed by *Pfu*SAHH.

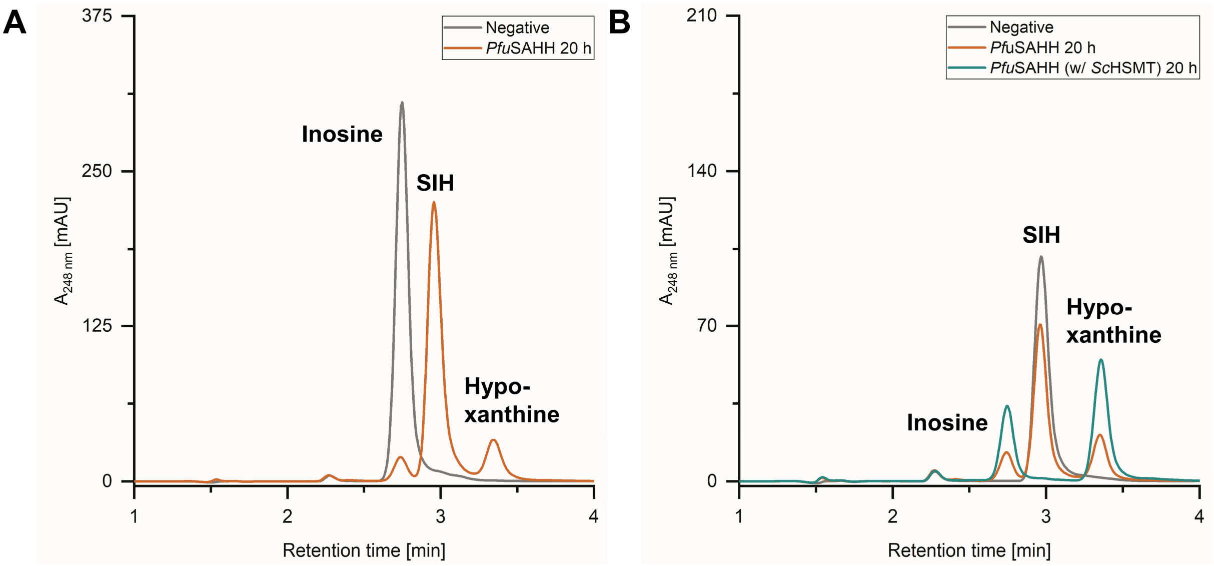

**Figure S6B.** HPLC chromatograms showing **A.** the SIH synthesis reaction and **B.** the SIH hydrolysis reaction (with and without the addition of *Sc*HSMT) catalysed by *Pfu*SAHH.

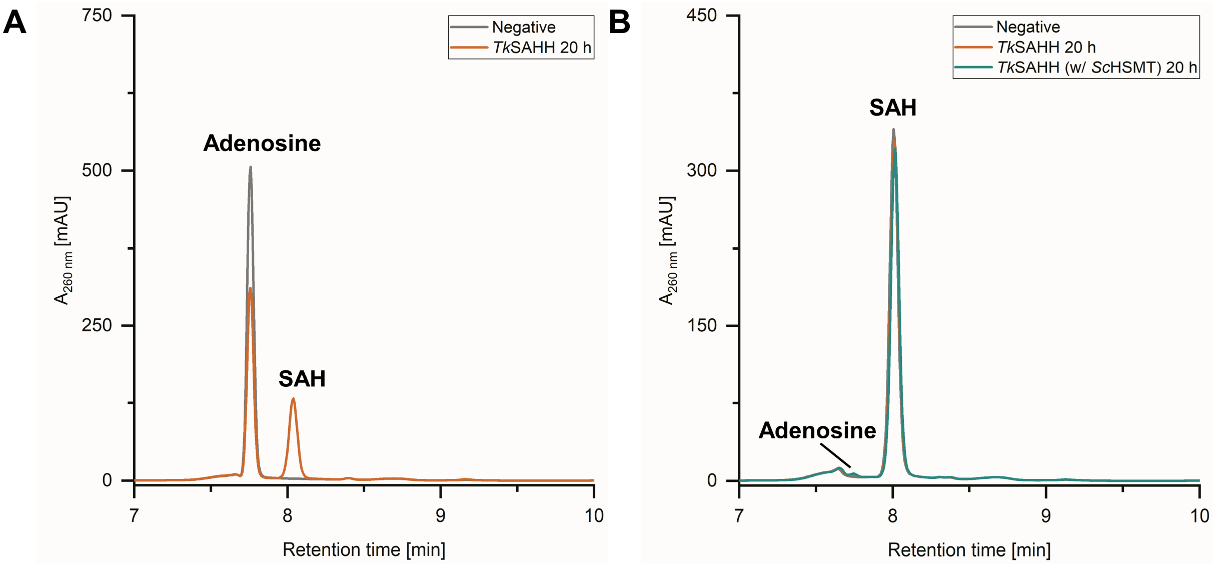

**Figure S7A.** HPLC chromatograms showing **A.** the SAH synthesis reaction and **B.** the SAH hydrolysis reaction (with and without the addition of *Sc*HSMT) catalysed by *Tk*SAHH.

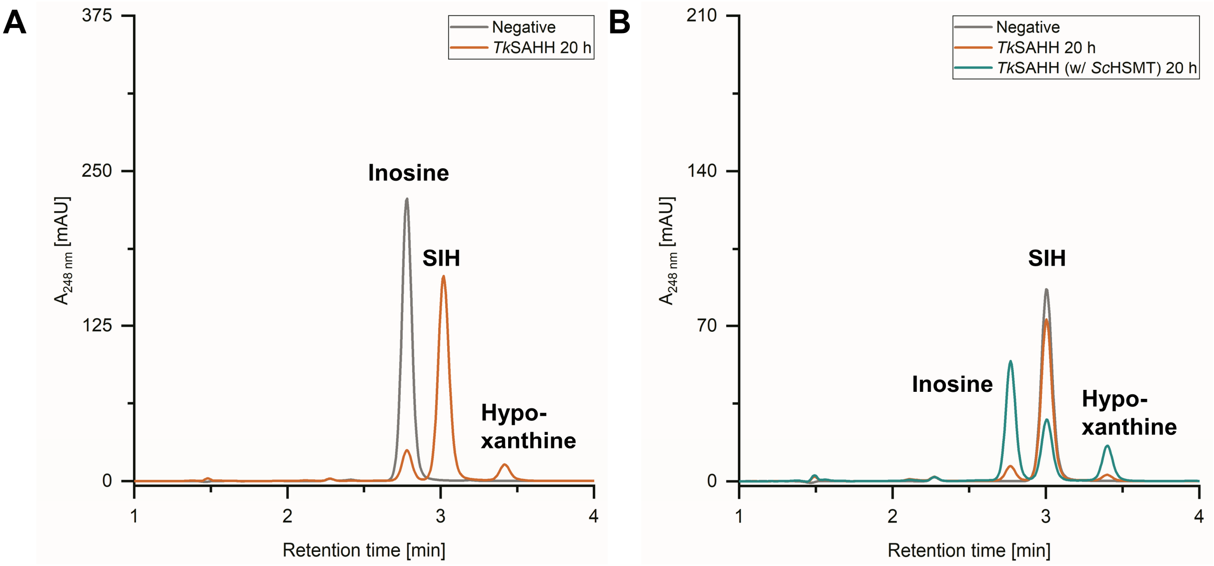

**Figure S7B.** HPLC chromatograms showing **A.** the SIH synthesis reaction and **B.** the SIH hydrolysis reaction (with and without the addition of *Sc*HSMT) catalysed by *Tk*SAHH.

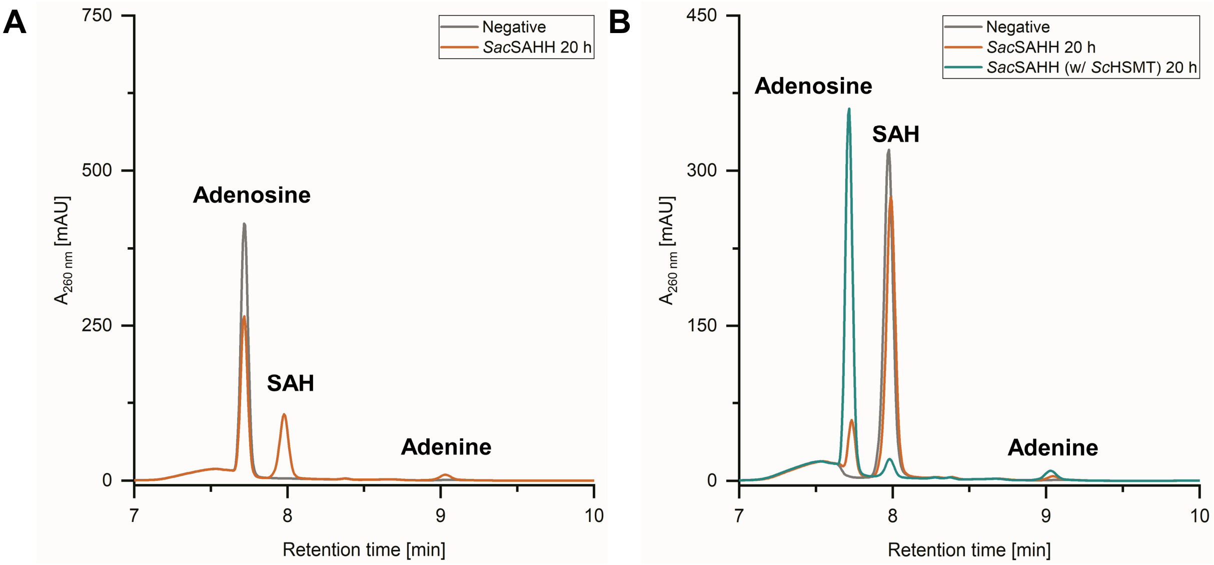

**Figure S8A.** HPLC chromatograms showing **A.** the SAH synthesis reaction and **B.** the SAH hydrolysis reaction (with and without the addition of *Sc*HSMT) catalysed by *Sac*SAHH.

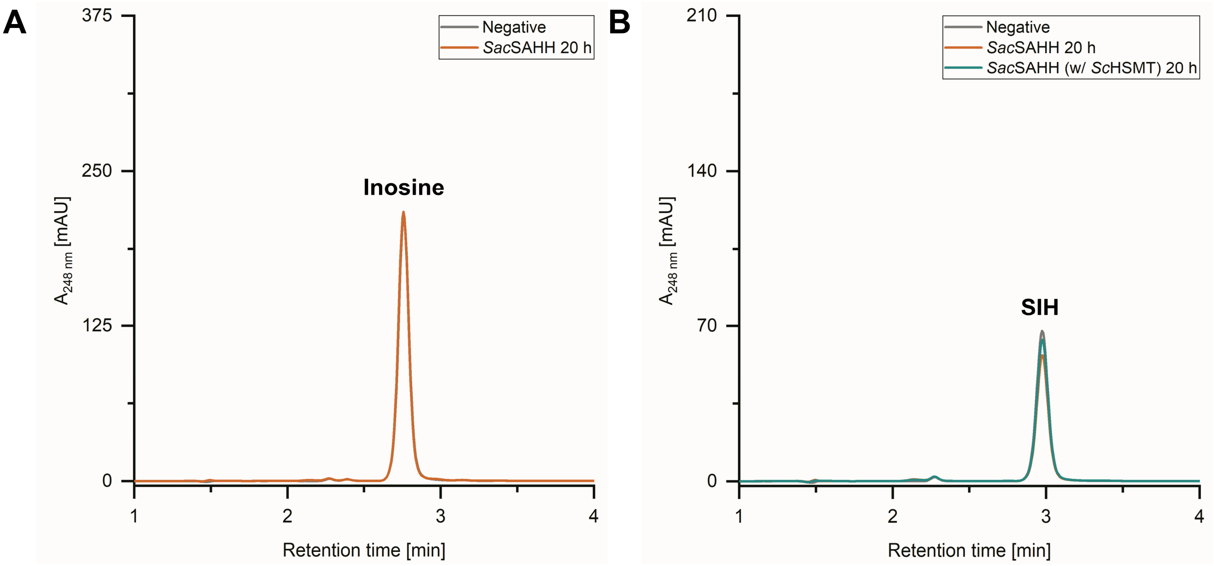

**Figure S8B.** HPLC chromatograms showing **A.** the SIH synthesis reaction and **B.** the SIH hydrolysis reaction (with and without the addition of *Sc*HSMT) catalysed by *Sac*SAHH.

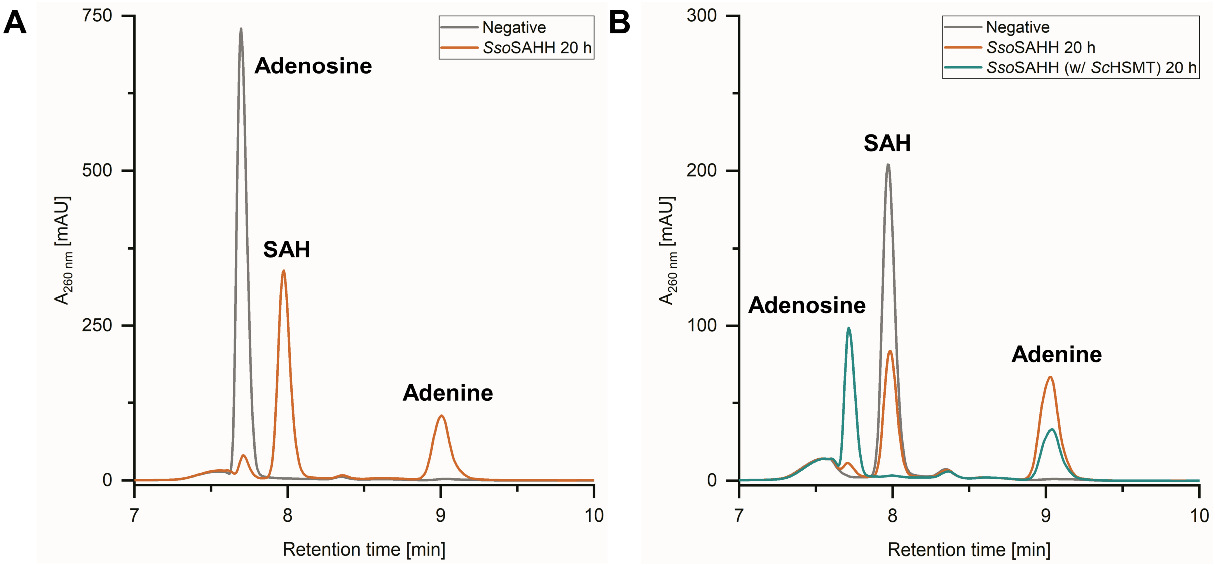

**Figure S9A.** HPLC chromatograms showing **A.** the SAH synthesis reaction and **B.** the SAH hydrolysis reaction (with and without the addition of *Sc*HSMT) catalysed by *Sso*SAHH.

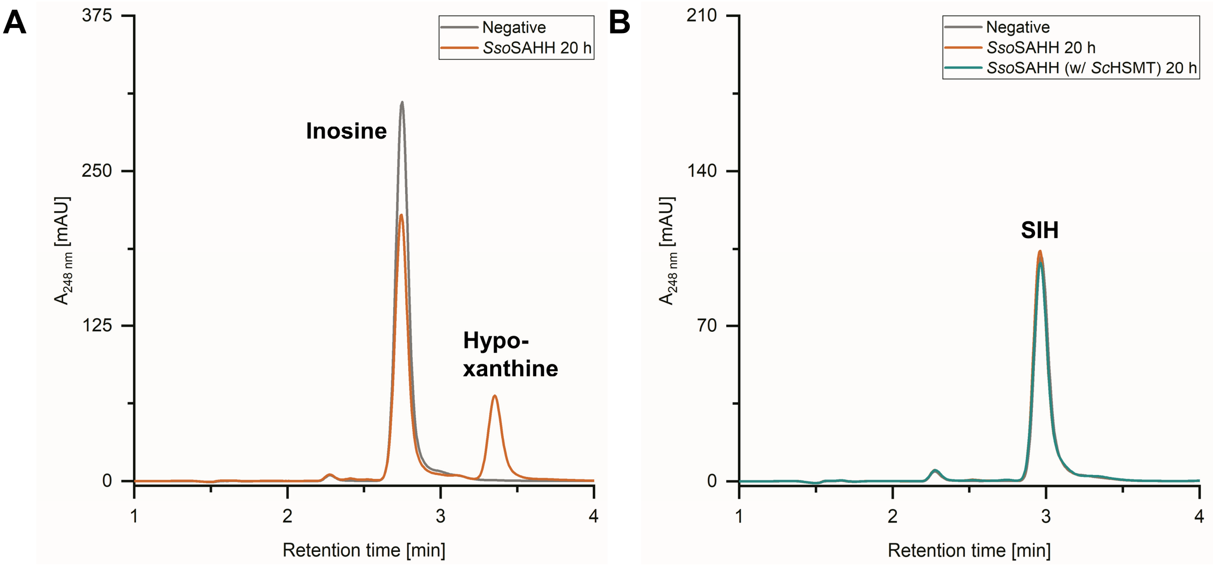

**Figure S9B.** HPLC chromatograms showing **A.** the SIH synthesis reaction and **B.** the SIH hydrolysis reaction (with and without the addition of *Sc*HSMT) catalysed by *Sso*SAHH.

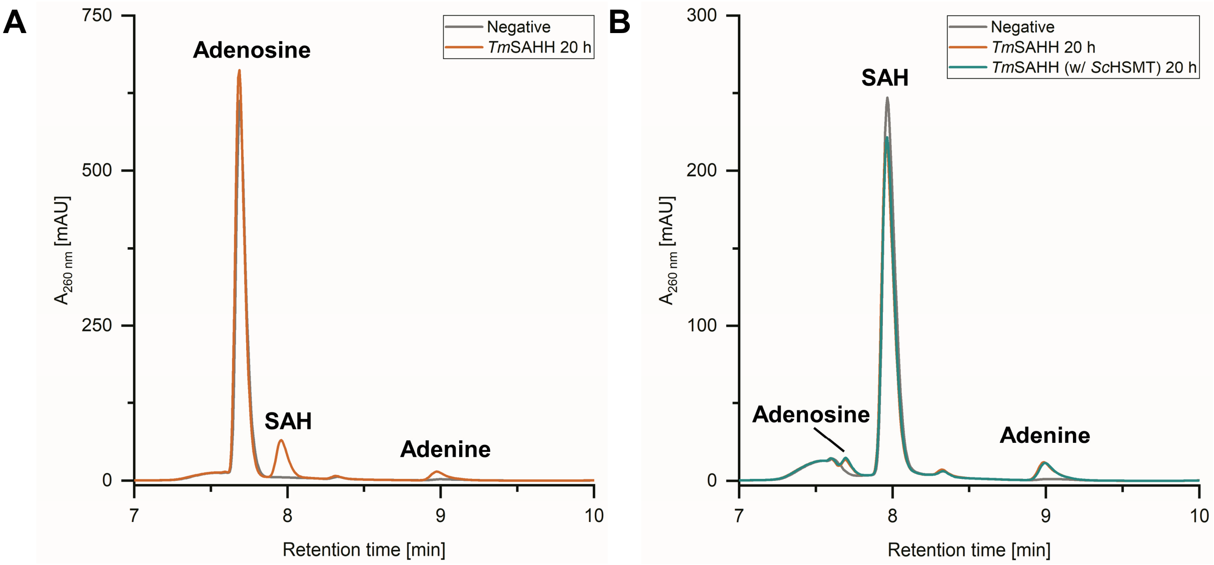

**Figure S10A.** HPLC chromatograms showing **A.** the SAH synthesis reaction and **B.** the SAH hydrolysis reaction (with and without the addition of *Sc*HSMT) catalysed by *Tm*SAHH.

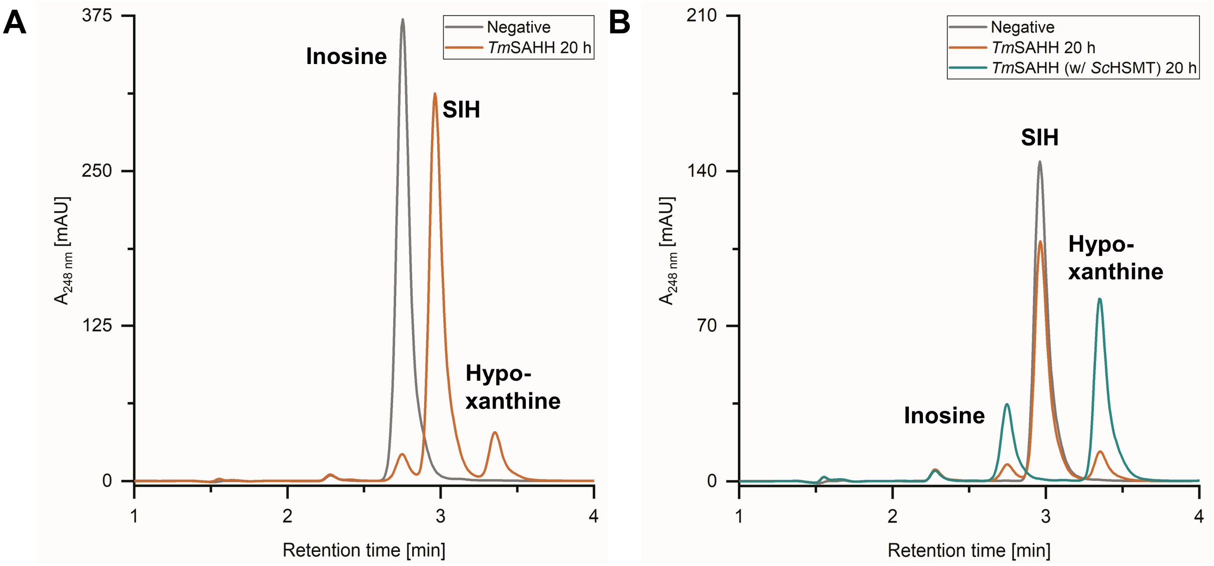

**Figure S10B.** HPLC chromatograms showing **A.** the SIH synthesis reaction and **B.** the SIH hydrolysis reaction (with and without the addition of *Sc*HSMT) catalysed by *Tm*SAHH.

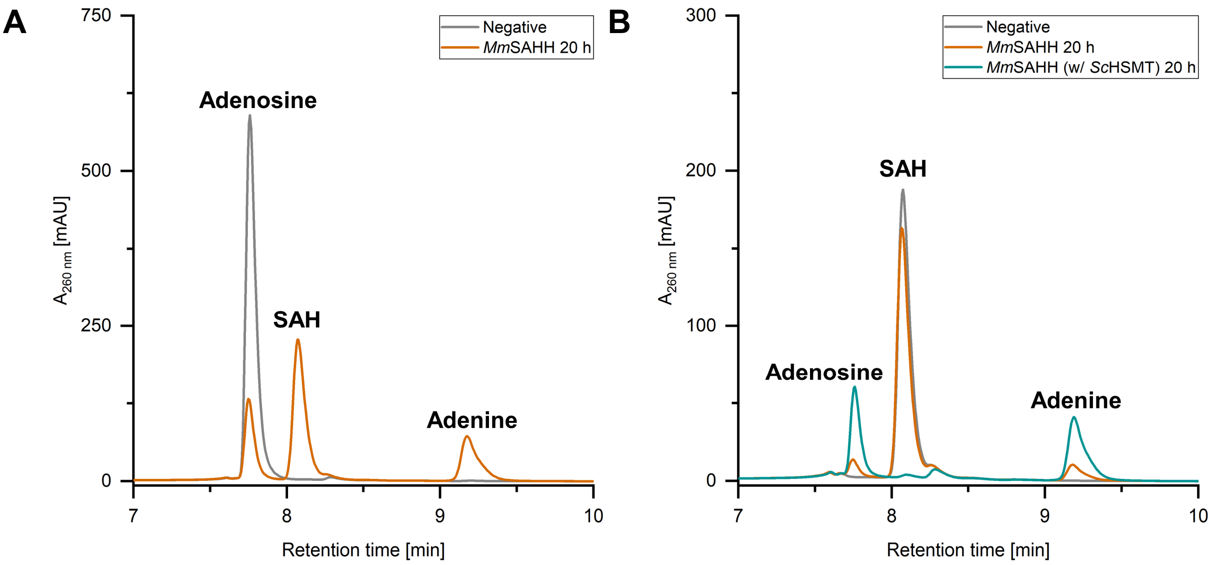

**Figure S11A.** HPLC chromatograms showing **A.** the SAH synthesis reaction and **B.** the SAH hydrolysis reaction (with and without the addition of *Sc*HSMT) catalysed by *Mm*SAHH.

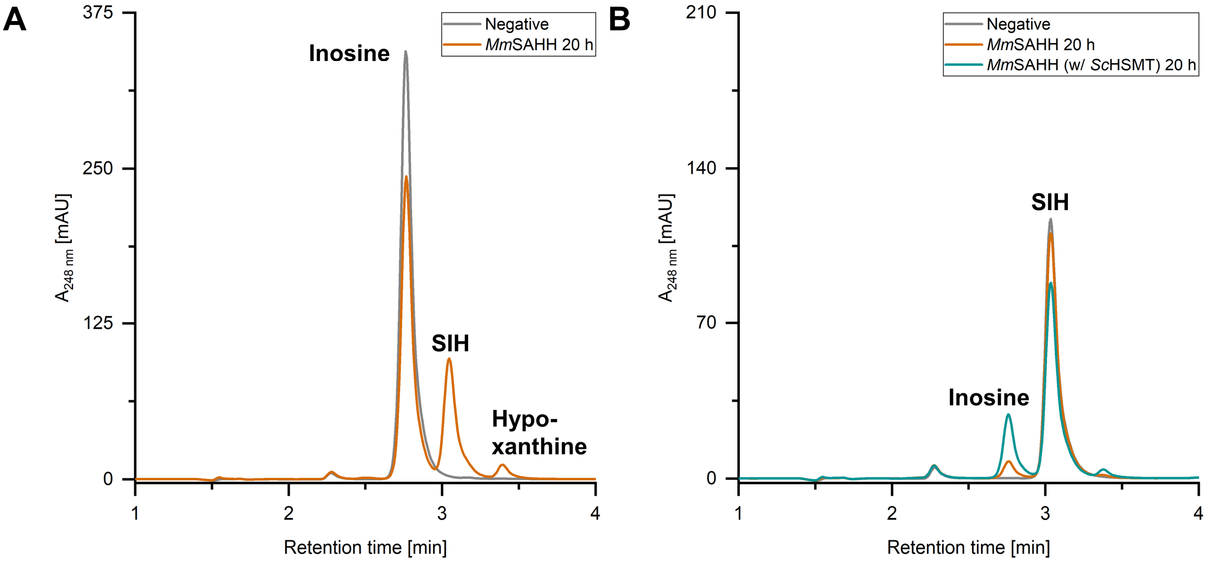

**Figure S11B.** HPLC chromatograms showing **A.** the SIH synthesis reaction and **B.** the SIH hydrolysis reaction (with and without the addition of *Sc*HSMT) catalysed by *Mm*SAHH.

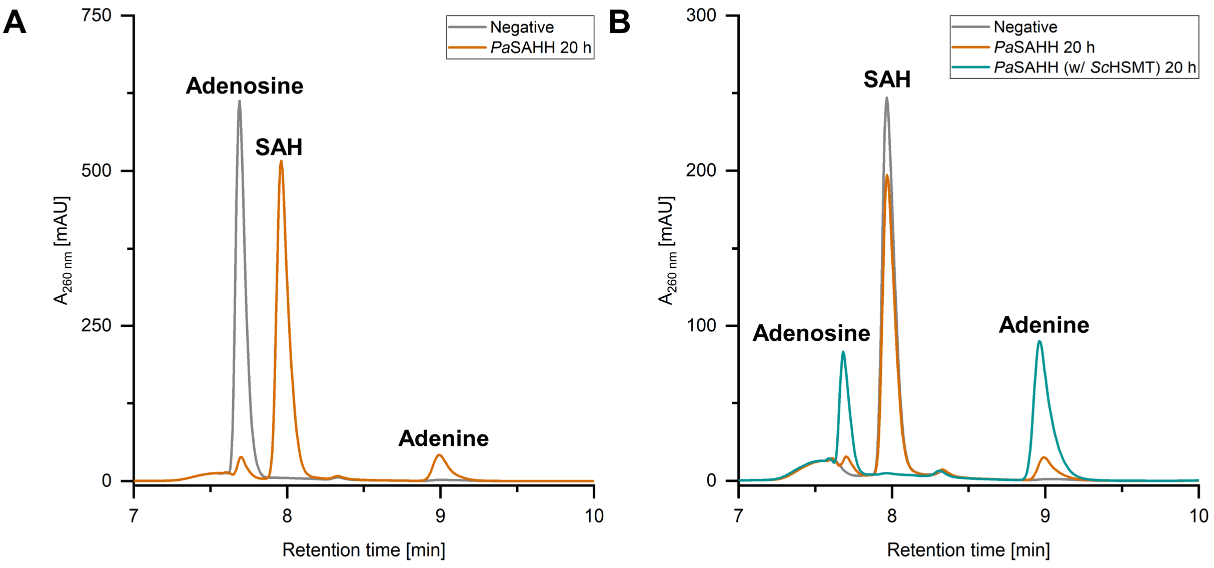

**Figure S12A.** HPLC chromatograms showing **A.** the SAH synthesis reaction and **B.** the SAH hydrolysis reaction (with and without the addition of *Sc*HSMT) catalysed by *Pa*SAHH.

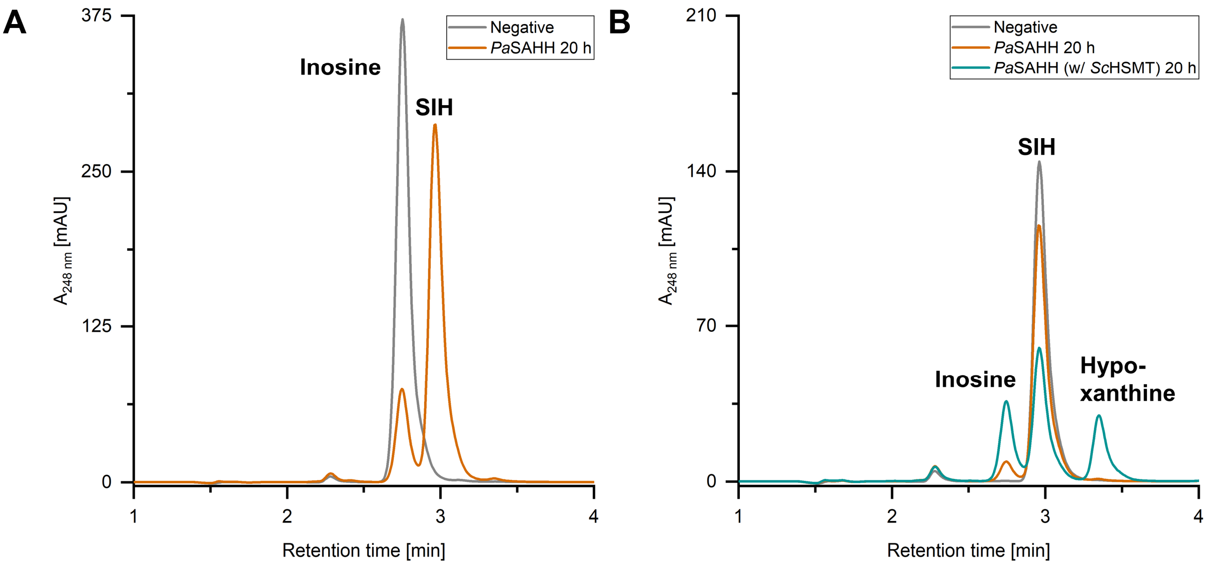

**Figure S12B.** HPLC chromatograms showing **A.** the SIH synthesis reaction and **B.** the SIH hydrolysis reaction (with and without the addition of *Sc*HSMT) catalysed by *Pa*SAHH.

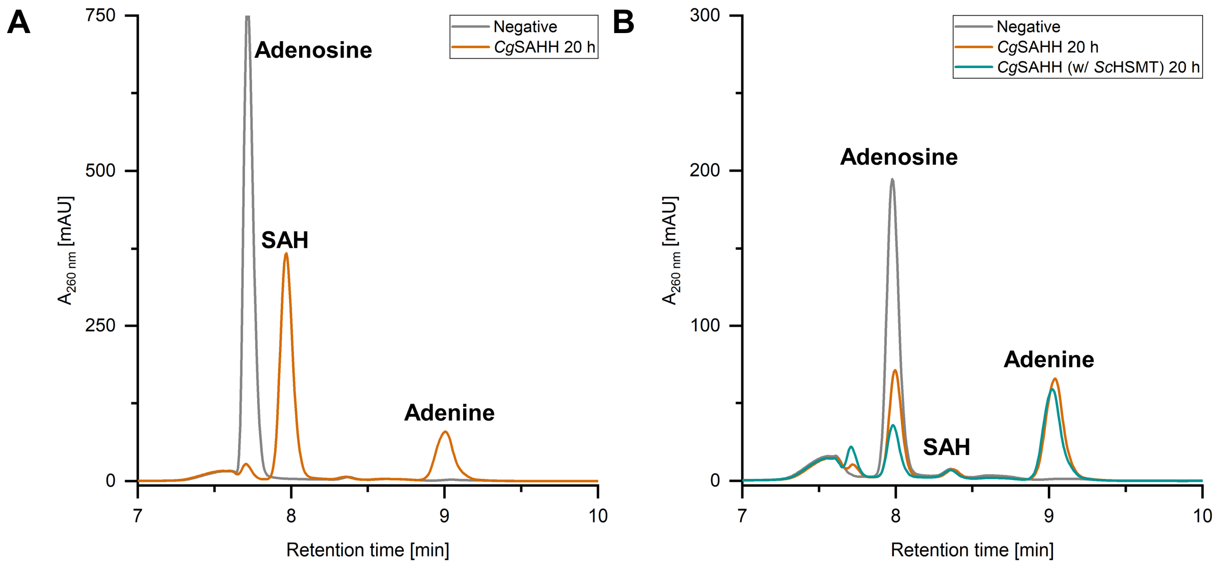

**Figure S13A.** HPLC chromatograms showing **A.** the SAH synthesis reaction and **B.** the SAH hydrolysis reaction (with and without the addition of *Sc*HSMT) catalysed by *Cg*SAHH.

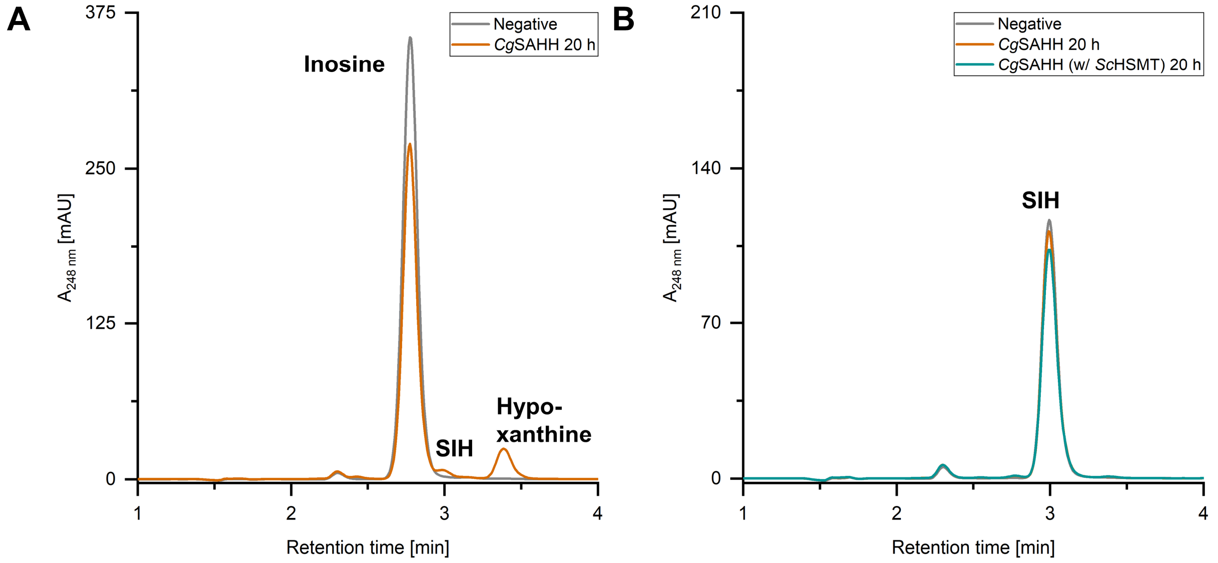

**Figure S13B.** HPLC chromatograms showing **A.** the SIH synthesis reaction and **B.** the SIH hydrolysis reaction (with and without the addition of *Sc*HSMT) catalysed by *Cg*SAHH.

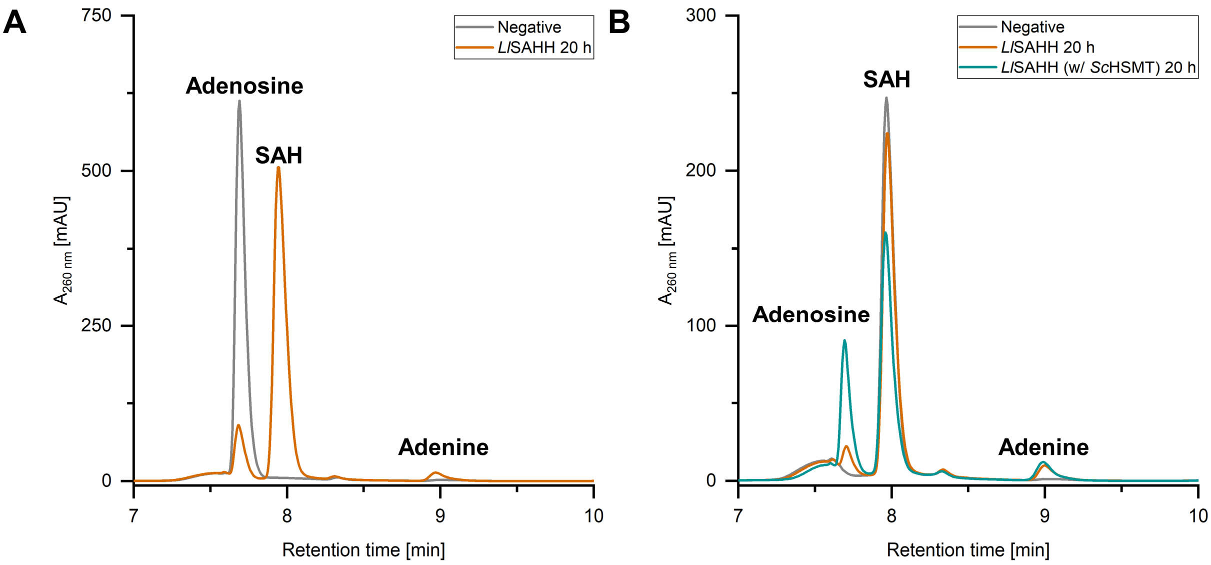

**Figure S14A.** HPLC chromatograms showing **A.** the SAH synthesis reaction and **B.** the SAH hydrolysis reaction (with and without the addition of *Sc*HSMT) catalysed by *Ll*SAHH.

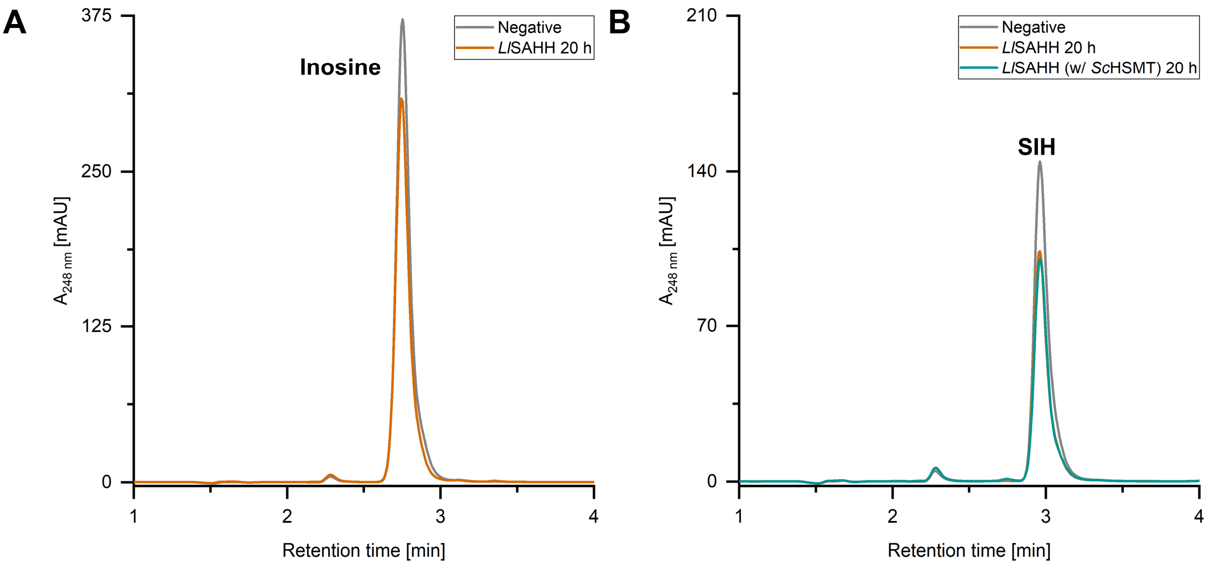

**Figure S14B.** HPLC chromatograms showing **A.** the SIH synthesis reaction and **B.** the SIH hydrolysis reaction (with and without the addition of *Sc*HSMT) catalysed by *Ll*SAHH.

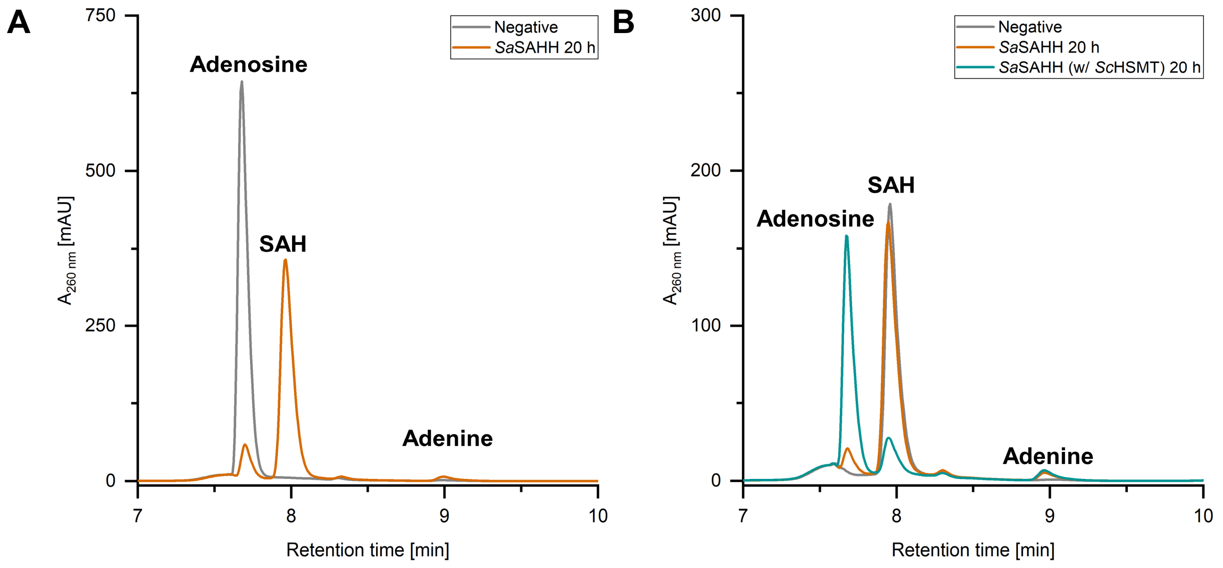

**Figure S15A.** HPLC chromatograms showing **A.** the SAH synthesis reaction and **B.** the SAH hydrolysis reaction (with and without the addition of *Sc*HSMT) catalysed by *Sa*SAHH.

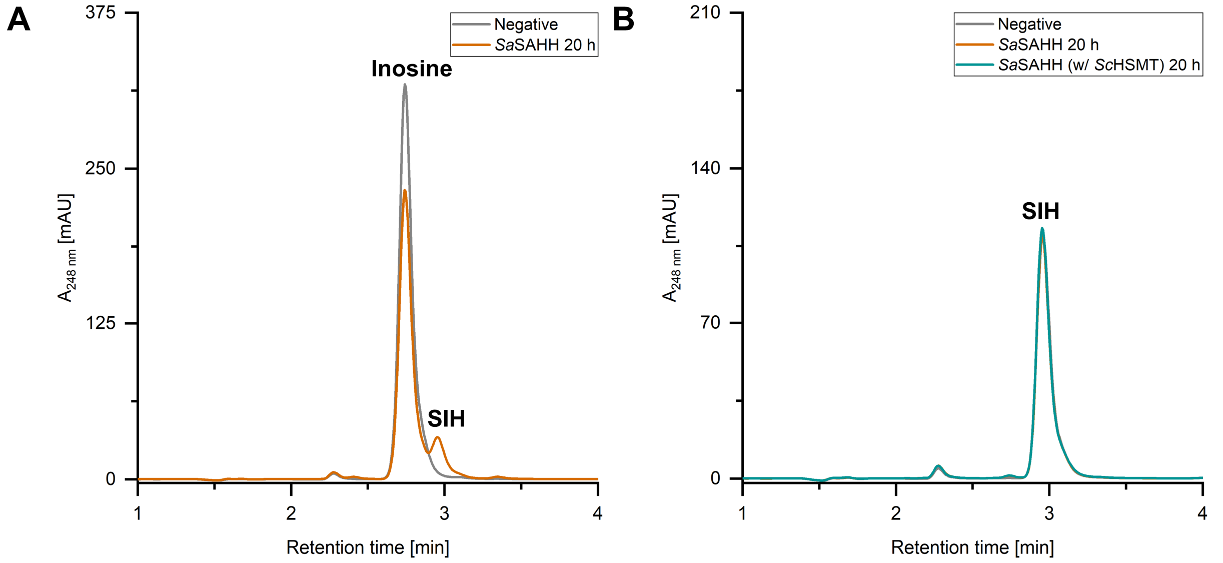

**Figure S15B.** HPLC chromatograms showing **A.** the SIH synthesis reaction and **B.** the SIH hydrolysis reaction (with and without the addition of *Sc*HSMT) catalysed by *Sa*SAHH.

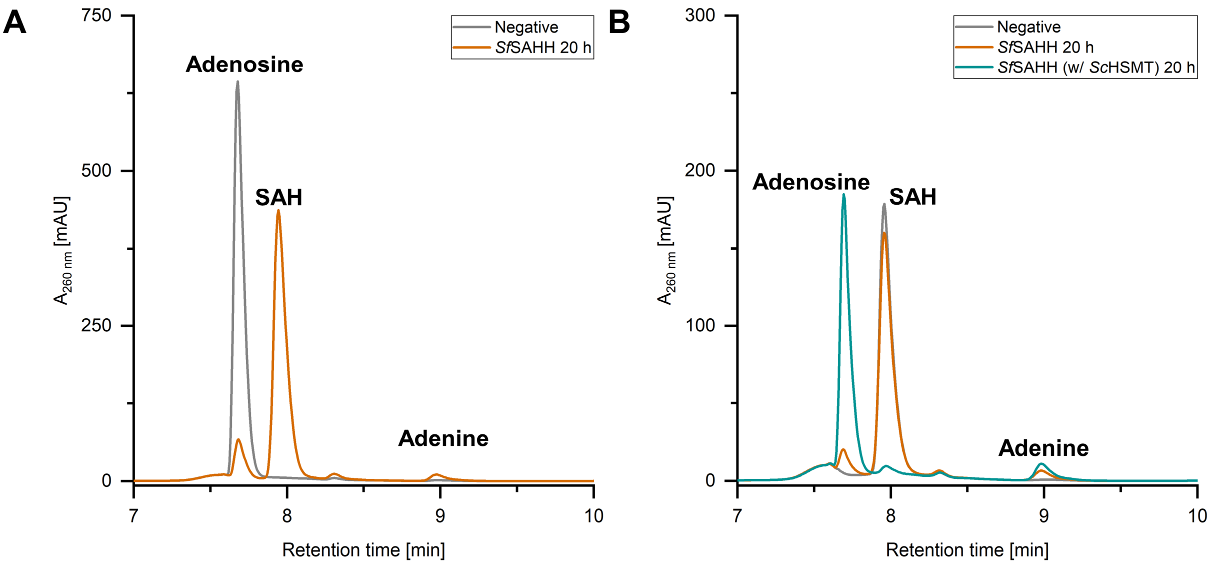

**Figure S16A.** HPLC chromatograms showing **A.** the SAH synthesis reaction and **B.** the SAH hydrolysis reaction (with and without the addition of *Sc*HSMT) catalysed by *Sf*SAHH.

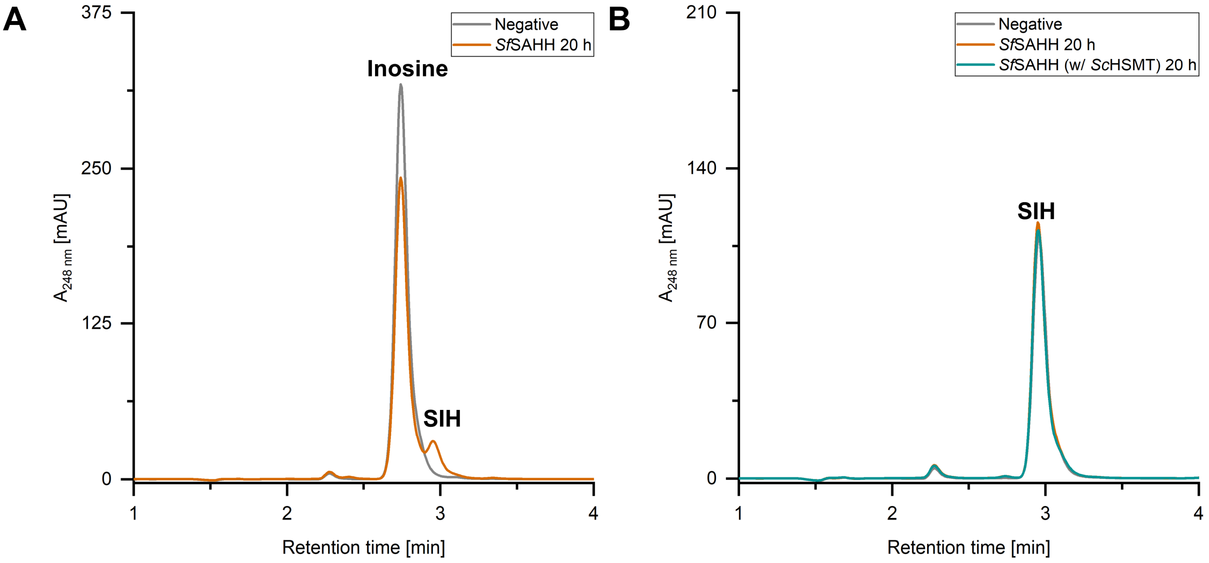

**Figure S16B.** HPLC chromatograms showing **A.** the SIH synthesis reaction and **B.** the SIH hydrolysis reaction (with and without the addition of *Sc*HSMT) catalysed by *Sf*SAHH.

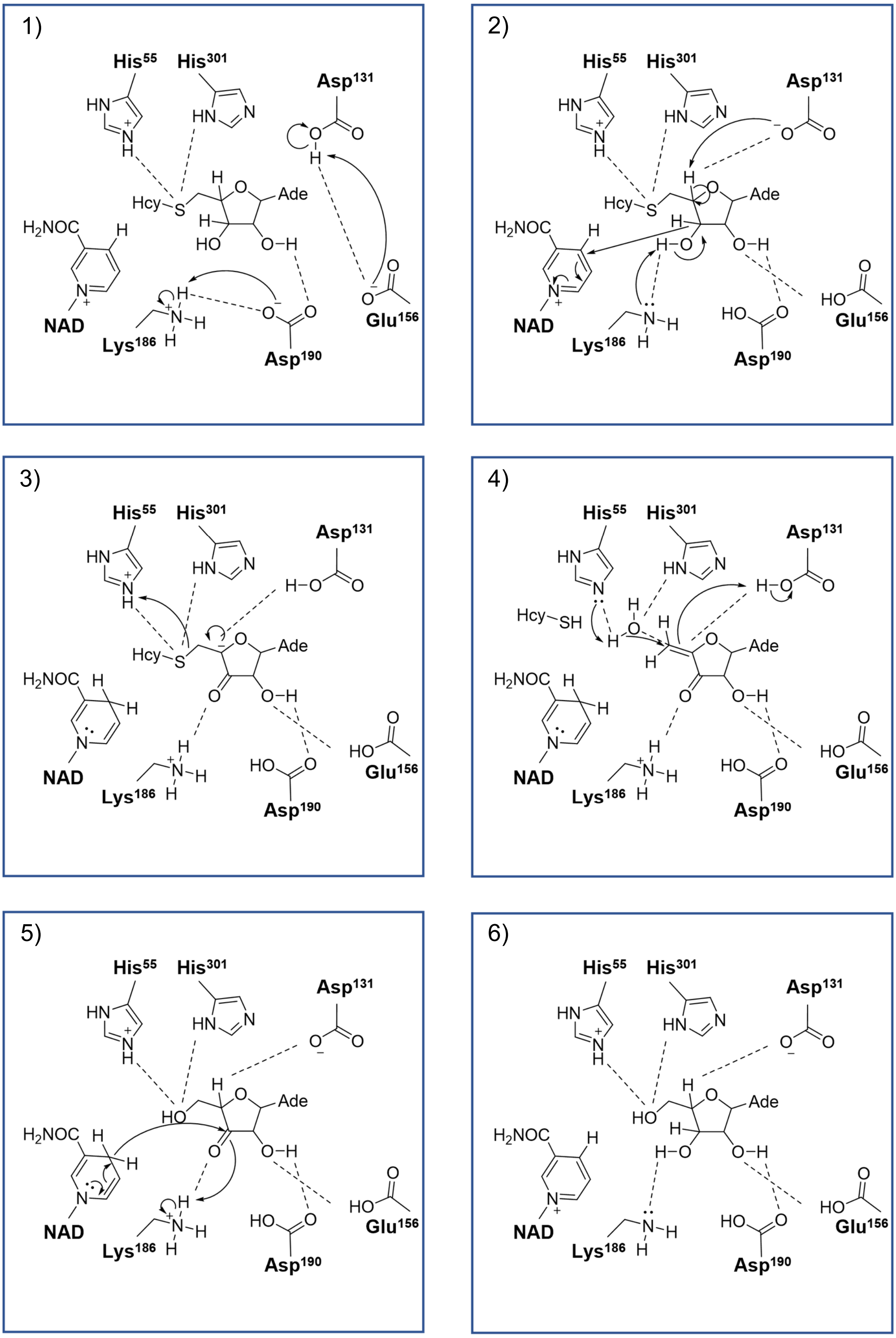

**Figure S17.** Molecular mechanism of SAH cleavage. 1) Interactions between amino acid residues before the catalytic reaction. 2) Formation of the 3´-keto intermediate and abstraction of the 4´-proton leading to the carbanion intermediate. 3) Formation of the 3´-keto-4´,5´dehydro intermediate after elimination of L-homocysteine. 4) Nucleophilic attack of an activated water molecule resulting in 3´-keto-adenosine. 5) Reduction of the 3´-keto-adenosine intermediate.6) Final product, adenosine still bound in the active site.

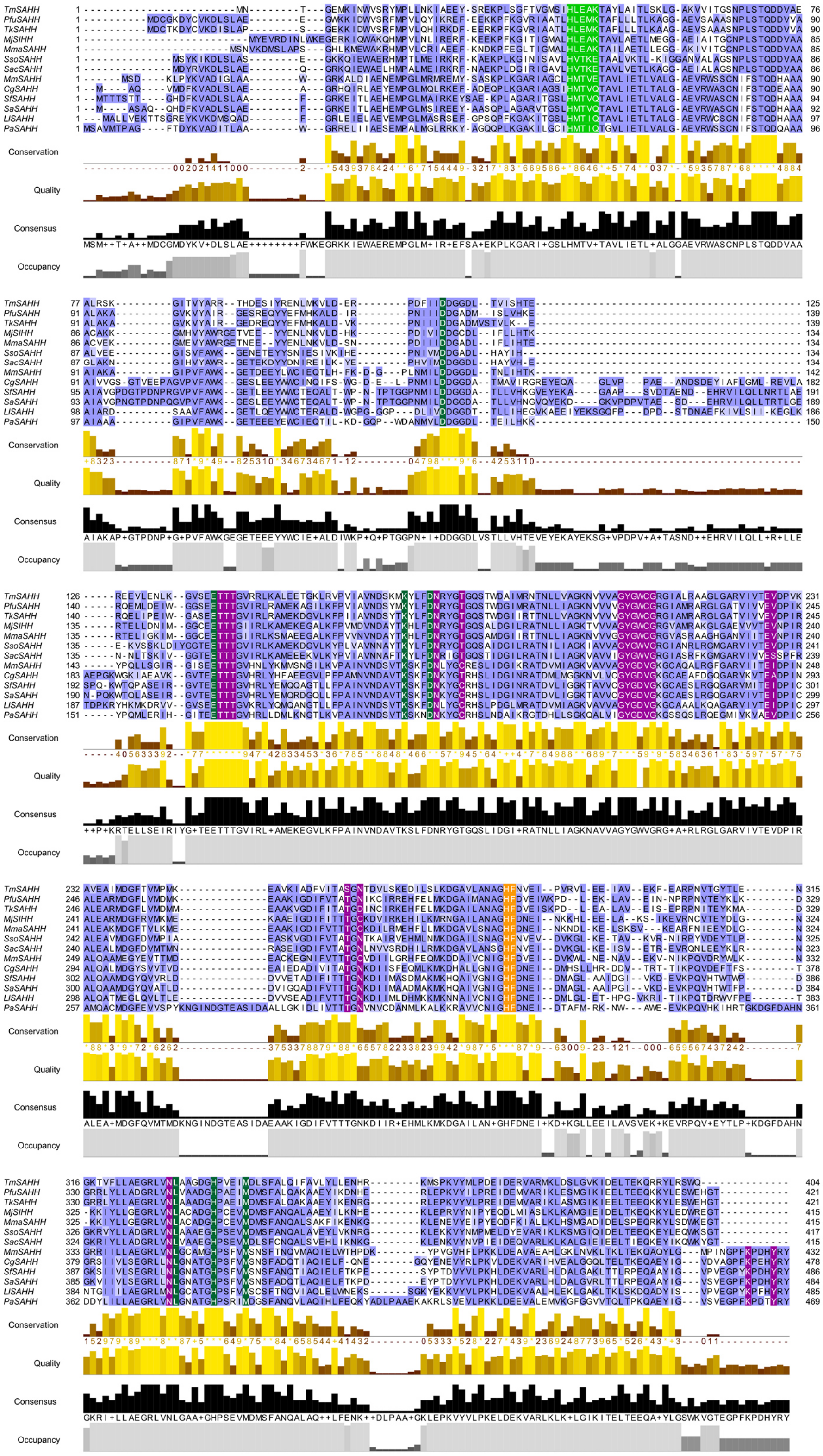

**Figure S18.** Multiple sequence alignment of selected SAHHs/SIHHs. Residues forming the sequence signature for binding of the nucleobase are coloured in light green. The other residues interacting with the substrate are coloured in dark green while the residues interacting with the cofactor $NAD^+$ are in purple. His and Phe of the molecular gate are coloured orange. The alignment was calculated in Clustal Omega[15] and the figure was prepared using Jalview Version2[16].

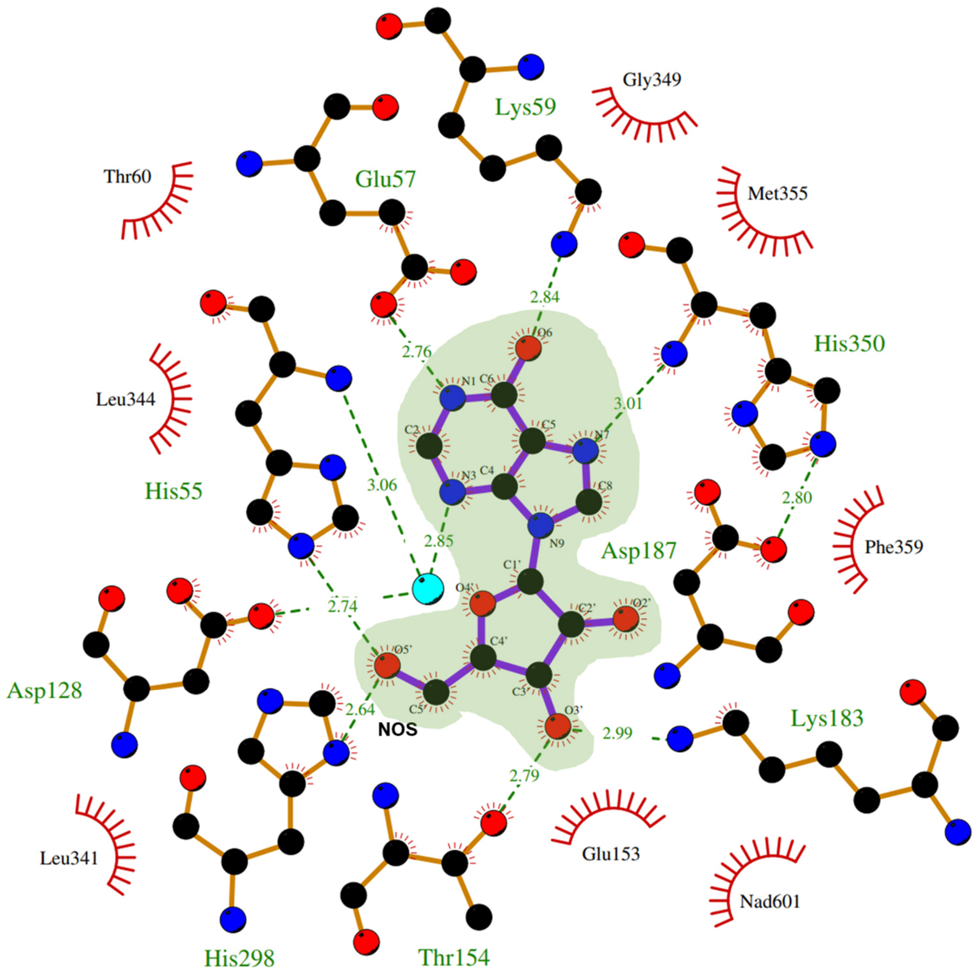

**Figure S19A.** LigPlot showing the interactions of *Pfu*SAHH-bound inosine (PDB ID: 7R37 chain A) with the amino acid backbones and side-chains in the active site.

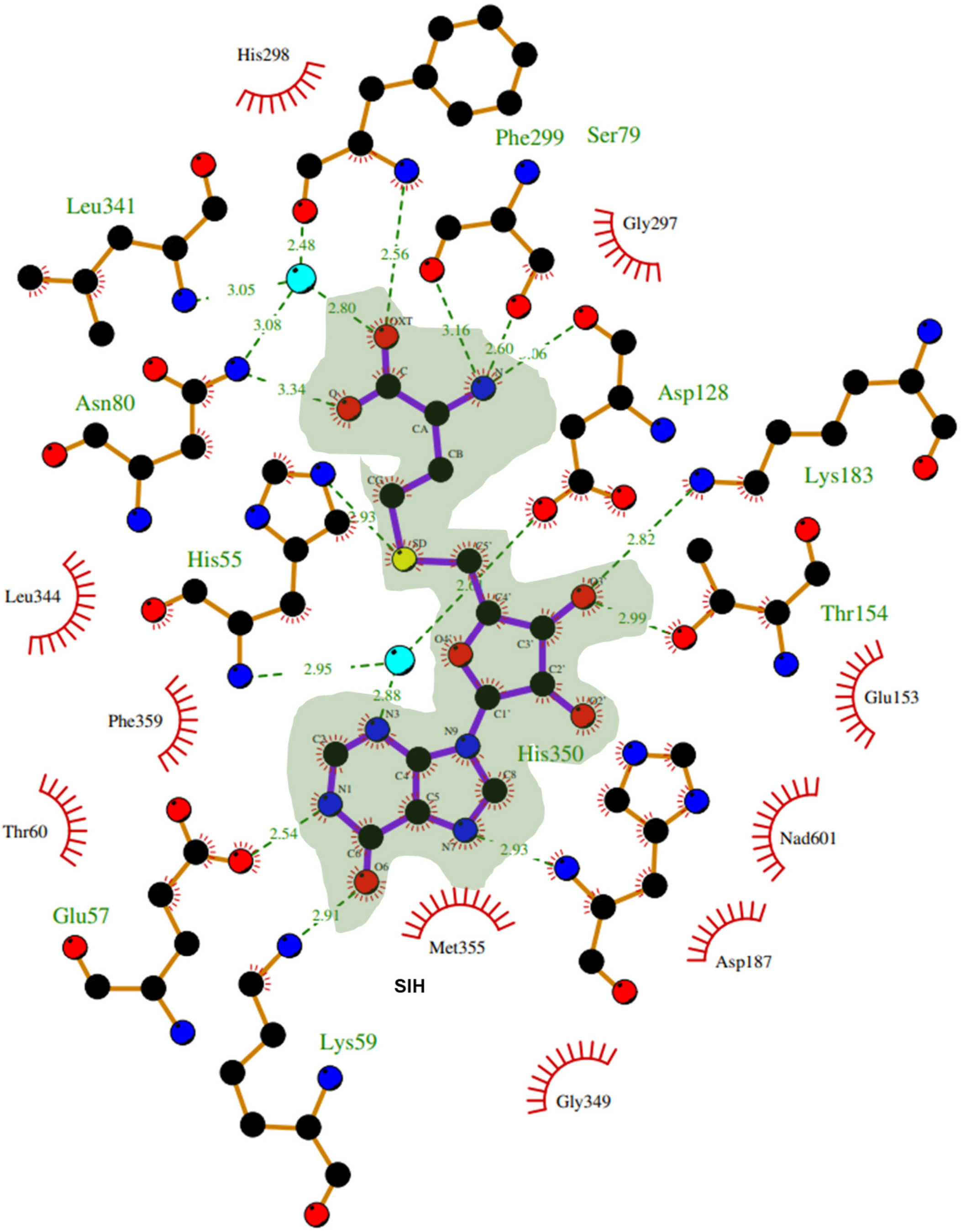

**Figure S19B**. LigPlot showing the interactions of *Pfu*SAHH-bound SIH PDB ID: 7R38 chain A) with the amino acid backbones and side-chains in the active site.

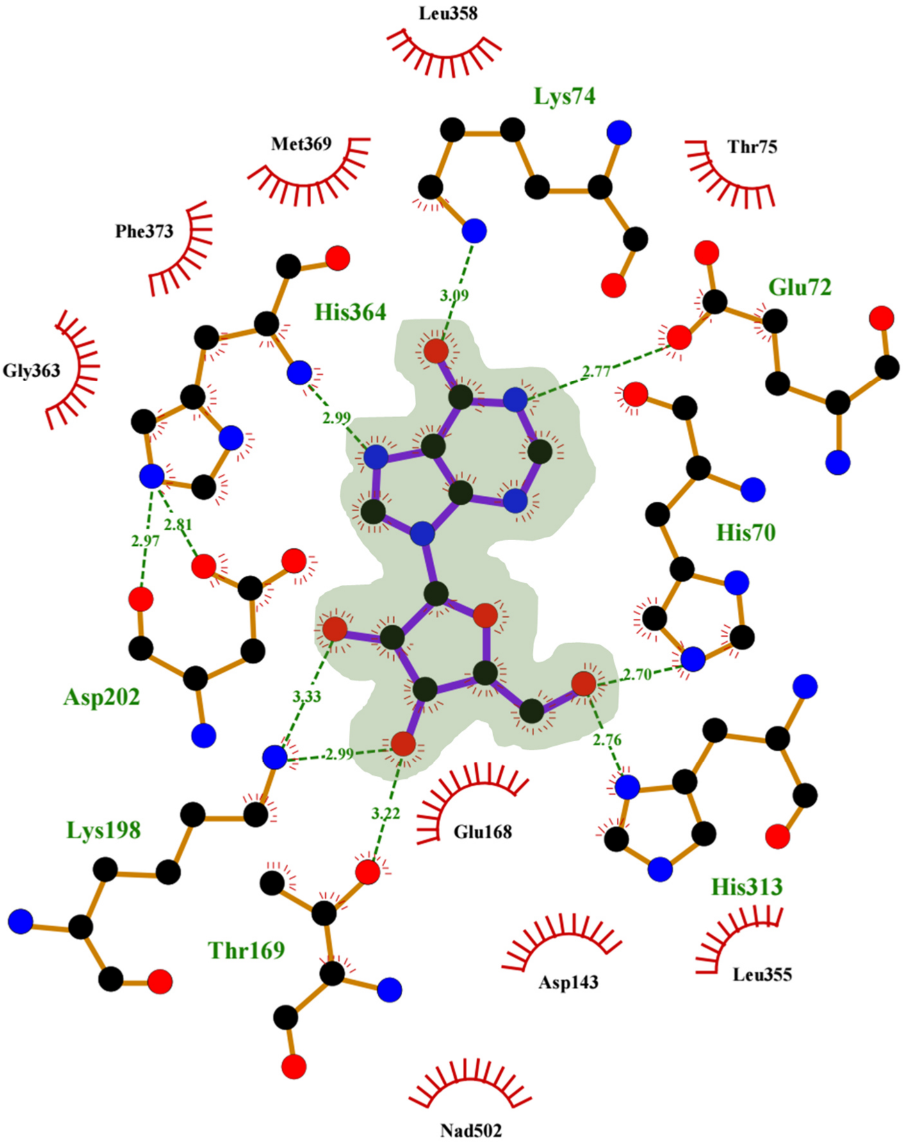

**Figure S19C**. LigPlot showing the interactions of *Mma*SAHH-bound inosine (PDB ID: 7R3A chain A) with the amino acid backbones and side-chains in the active site.

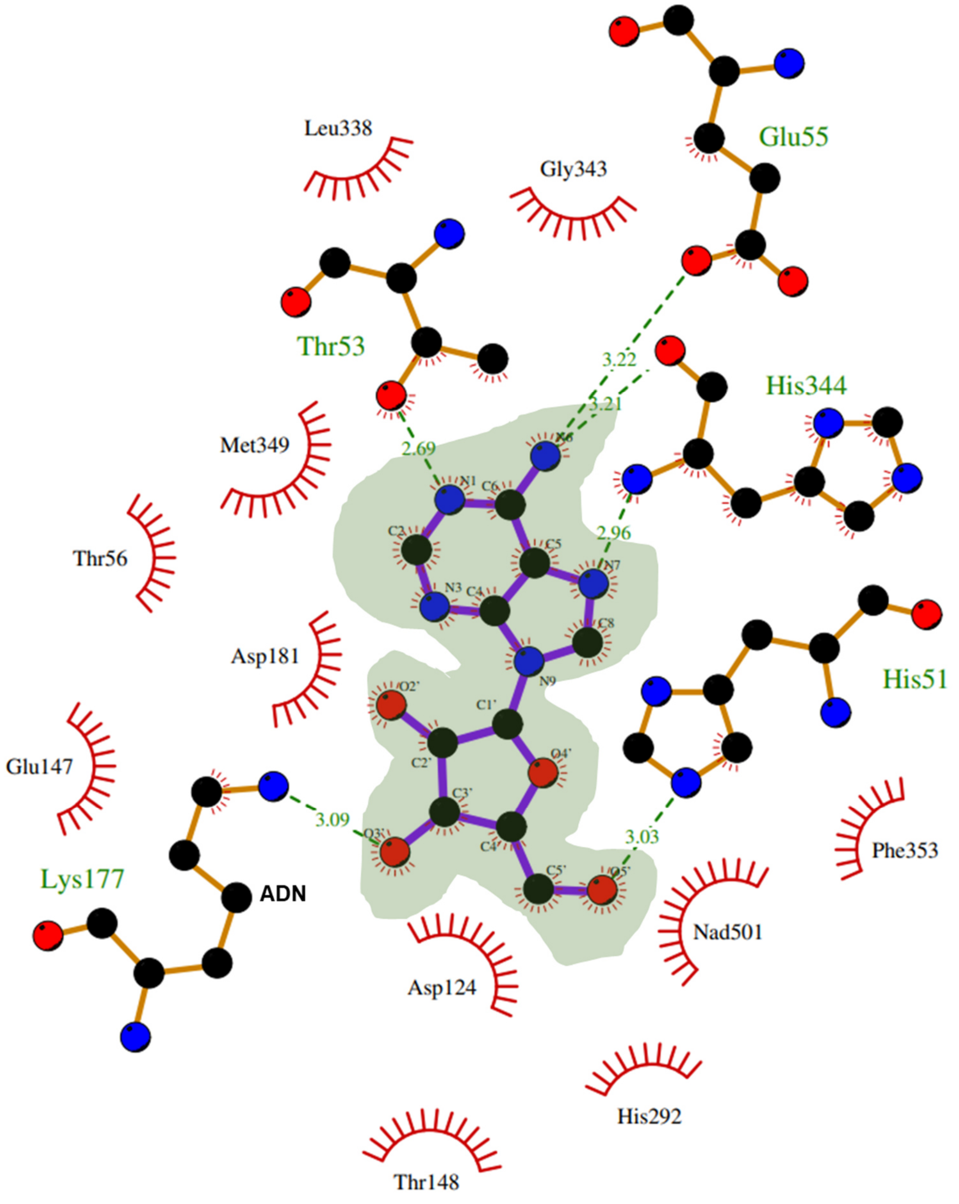

**Figure S19D**. LigPlot showing the interactions of *Sac*SAHH-bound adenosine (PDB ID: 7R39 chain A) with the amino acid backbones and side-chains in the active site.

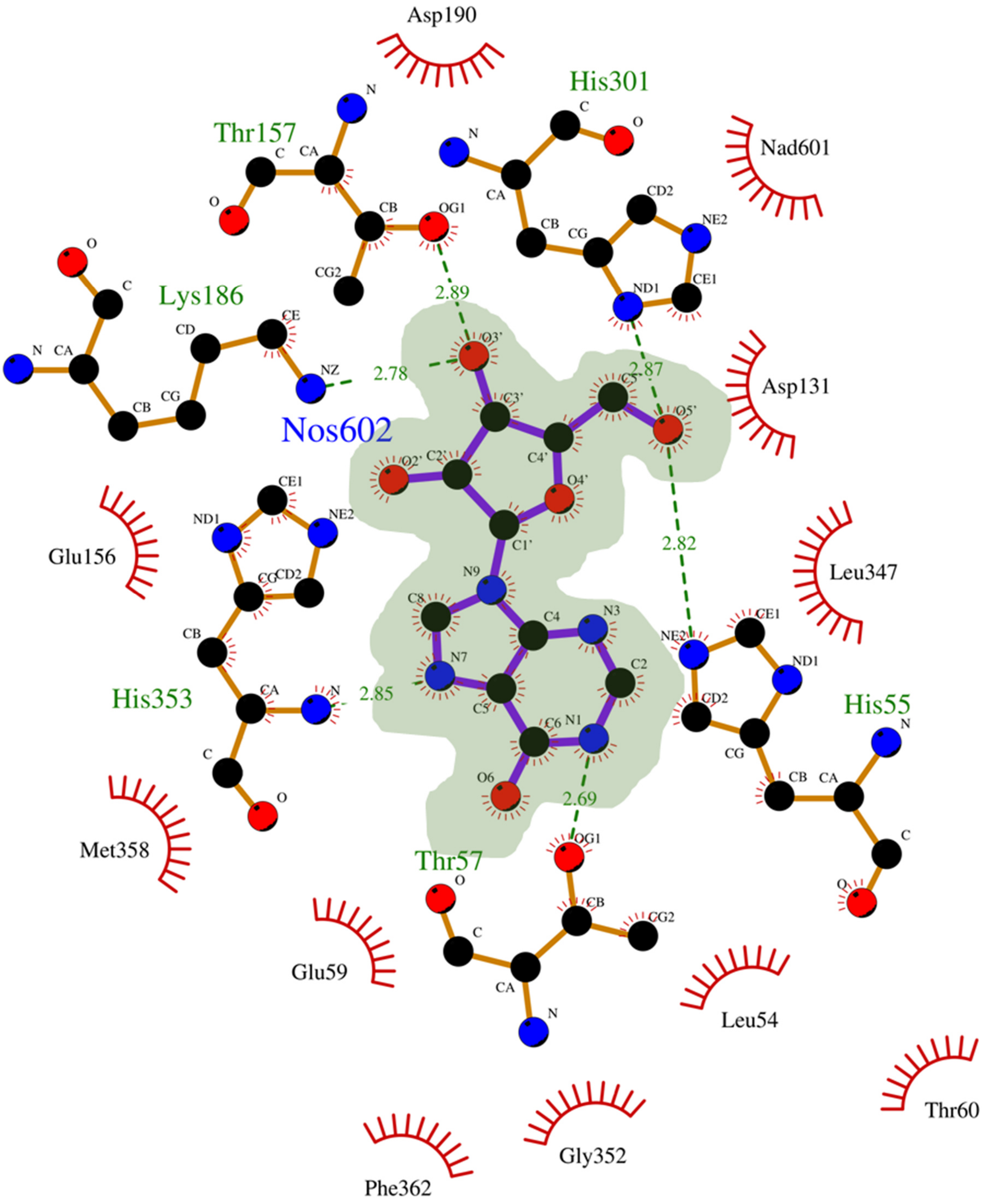

**Figure S19E**. LigPlot showing the interactions of *Mm*SAHH-bound inosine (PDB ID: 8COD chain A) with the amino acid backbones and side-chains in the active site.

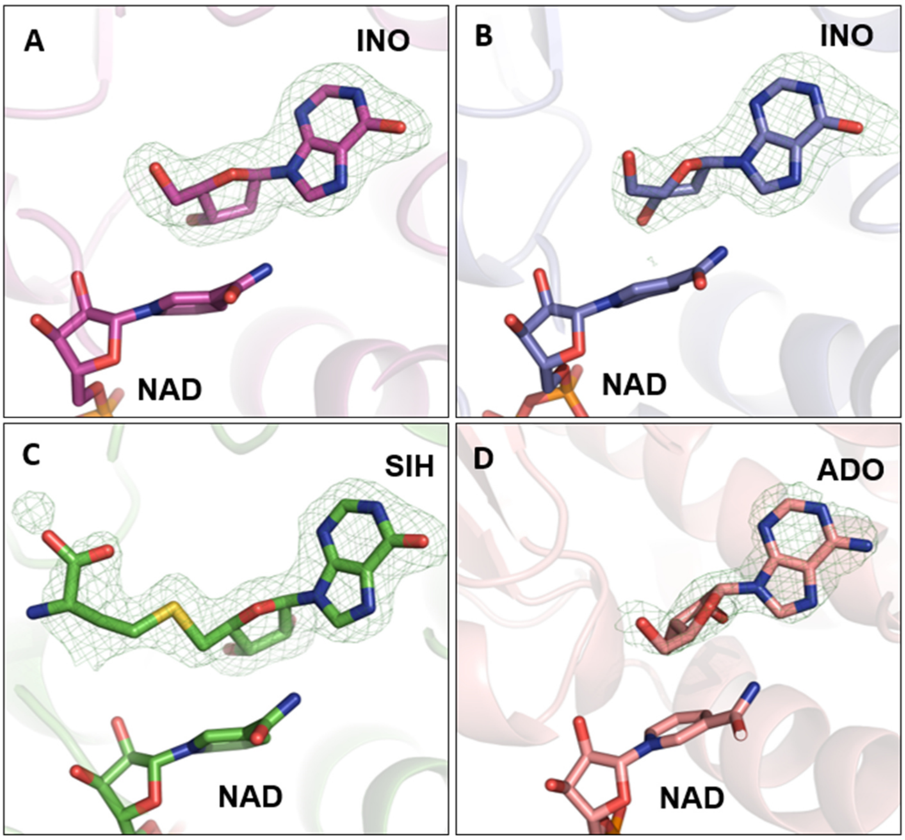

**Figure S19F**. The mFo-DFc polder OMIT difference electron density maps (green mesh) of the bound nucleosides and are contoured at 3.0σ above the mean. Mode of inosine (INO) in the *Pfu*SAHH active site (A) (PDB ID: 7R37), *Mma*SAHH active site (B) (PDB ID: 7R3A), SIH at *Pfu*SAHH active site (C) (PDB ID: 7R38) and adenosine (ADO) in *Sac*SAHH active site (D) (PDB ID: 7R39).

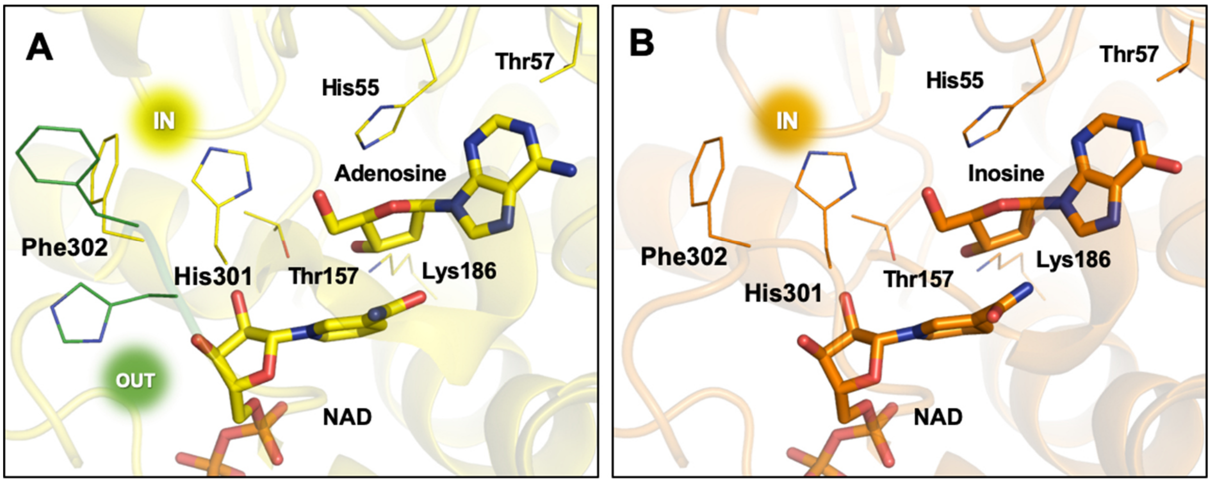

**Figure S19G**. The molecular gate residues in the crystal structures of *Mus musculus* SAHHs. The His residue states are represented as His-IN (IN) and His-OUT (OUT). A: In the *Mm*SAHH•NAD•Adenosine complex (PDB ID: 5AXA), the gatekeeper residue forms both IN and OUT conformation leaving the channel gate shut and open, respectively. B: In the *Mm*SAHH•NAD•inosine complex (PDB ID: 8COD), the His gatekeeper residue is in the IN orientation thereby closing the channel entrance.

**Table S5.** Homologues of *Mj*DadD found in the genomes whose SAHH was characterised in this work, using BLASTP[17].

| Organism | UniProt accession number | Sequence identity/similarity [%] to *Mj*DadD |
|---|---|---|
| *Corynebacterium glutamicum* | Not available | Not available |
| *Lupinus luteus* | Not available | Not available |
| *Methanocaldococcus jannaschii* | Q58936 (DADD_METJA) | 100.0/100.0 |
| *Methanococcus maripaludis* | Q6LX61 (DADD_METMP) | 69.2/83.4 |
| *Mus musculus* | Not available | Not available |
| *Pseudomonas aeruginosa* | Not available | Not available |
| *Pyrococcus furiosus* | Q8U0P7 (MTAD_PYRFU) | 46.0/65.0 |
| *Sulfolobus acidocaldarius* | Not available | Not available |
| *Saccharolobus solfataricus* | Not available | Not available |
| *Streptomyces albus* | Not available | Not available |
| *Streptomyces flocculus* | Not available | Not available |
| *Thermococcus kodakarensis* | Q5JER0 (MTAD_THEKO) | 50.6/67.4 |
| *Thermotoga maritima* | Q9X034 (MTAD_THEMA) | 39.1/57.9 |

**Table S6.** Sequence identities given in percent [%] based on the amino acid sequence for all investigated SAHHs in this work.

| | ***Sac*** | ***Sso*** | ***Mj*** | ***Mma*** | ***Pfu*** | ***Tk*** | ***Cg*** | ***Pa*** | ***Sa*** | ***Sf*** | ***Tm*** | ***Ll*** | ***Mm*** |
|---|---|---|---|---|---|---|---|---|---|---|---|---|---|
| ***Sac*** | | | | | | | | | | | | | |
| ***Sso*** | 72.05 | - | | | | | | | | | | | |
| ***Mj*** | 59.32 | 60.14 | - | | | | | | | | | | |
| ***Mma*** | 56.42 | 57.25 | 75.66 | - | | | | | | | | | |
| ***Pfu*** | 59.04 | 61.3 | 63.61 | 59.52 | - | | | | | | | | |
| ***Tk*** | 57.59 | 60.34 | 62.89 | 60.48 | 88.12 | - | | | | | | | |
| ***Cg*** | 47.71 | 48.2 | 42.65 | 42.17 | 43.17 | 43.71 | - | | | | | | |
| ***Pa*** | 40.82 | 41.35 | 38.89 | 35.99 | 38.1 | 38.1 | 54.76 | - | | | | | |
| ***Sa*** | 46.51 | 46.28 | 43.37 | 42.17 | 43.23 | 42.52 | 68.2 | 58.39 | - | | | | |
| ***Sf*** | 46.75 | 46.28 | 44.34 | 42.41 | 43.71 | 42.99 | 67.57 | 57.67 | 89.26 | - | | | |
| ***Tm*** | 52.99 | 53.35 | 57.07 | 56.08 | 59.9 | 58.91 | 40.84 | 36.97 | 39.6 | 39.85 | - | | |
| ***Ll*** | 44.1 | 45.08 | 42.89 | 43.13 | 42.04 | 40.38 | 58.33 | 52.27 | 65.12 | 65.68 | 37.87 | - | |
| ***Mm*** | 47.1 | 49.28 | 44.44 | 43.24 | 46.67 | 45.24 | 58.8 | 59.07 | 64.58 | 64.81 | 41.94 | 62.79 | - |

**Table S7.** Overview of SAHHs with solved structures deposited in the PDB databank.

| **Organism** | **Domain, Phylum/Kingdom** | **PDB-ID** | **Bound Molecules** | **Reference** |
|---|---|---|---|---|
| *Acanthamoeba castellanii* | Eukaryota, Amoebozoa | 6UK3 | Adenosine | Unpublished |
| *Bradyrhizobium elkanii* | Bacteria, Proteobacteria | 4LVC | Adenosine | Manszewski *et al.*, 2015[18] |
| | | 5M65<br>5M66<br>5M67<br>5M5K | Adenine<br>Adenosine<br>Adenine, 2´-deoxyadenosine<br>Adenosine, Cordycepin | Manszewski *et al.*, 2017[19] |
| | | 6EXI | $NAD^+$-free, Adenosine | Kailing *et al.*, 2018[20] |
| *Brucella abortus* | Bacteria, Proteobacteria | 3N58 | Adenosine | Unpublished |
| *Burkholderia pseudomallei* | Bacteria, Proteobacteria | 3D64<br>3GLQ | None<br>9-β-D-arabinofuranosyladenine | Unpublished |
| *Cryptosporidium parvum* | Eukaryota, Apicomplexa | 5HM8<br>5T8K<br>5TJ9<br>5TLS<br>5UTU | Adenosine<br>Adenine<br>Aristeromycin<br>DZ2002<br>SAH, Adenosine | Unpublished |
| *Elizabethkingia anophelis* | Bacteria, Bacteroidetes | 6APH | Adenosine | Unpublished |

| | | | | |
|---|---|---|---|---|
| *Homo sapiens* | Eukaryota, Animalia | 1A7A | Adenosine analogue | Turner *et al.*, 1998[21] |
| | | 1LI4 | Neplanocin | Yang *et al.*, 2003[22] |
| | | 3GVP | None | Unpublished |
| | | 3NJ4 | Fluoro-neplanocin A | Lee *et al.*, 2011[23] |
| | | 4YVF | Complex Isoindoline/chloroaniline inhibitor | Nakao *et al.*, 2015[24] |
| | | 5W4B | Benzothiazole inhibitor | Uchiyama *et al* ., 2017[25] |
| | | 5W49 | Oxadiazole inhibitor | |
| | | 4PGF | Adenosine (mono-acetylated protein | Wang *et al.*, 2014[26] |
| | | 4PFJ | Adenosine (bi-acetylated protein) | |
| *Leishmania major* | Eukaryota, Euglenozoa | 3G1U | Adenosine | Unpublished |
| *Lupinus luteus* | Eukaryota, Plantae | 3OND | Adenosine | Brzezinski *et al.*, 2012[3] |
| | | 3ONE | Adenine | |
| | | 3ONF | 3´-deoxyadenosine | |
| *Mus musculus* | Eukaryota, Animalia | 5AXA | Adenosine | Kusakabe *et al.*, 2015[7] |
| | | 5AXB | Noraristeromycin | |
| | | 5AXC | 3´-keto aristeromycin | |
| | | 5AXD | Ribavirin | |
| *Mycobacterium tuberculosis* | Bacteria, Actinobacteria | 2ZIZ | 3-deazaadenosine | Reddy *et al.*, 2008[27] |
| | | 2ZJ0 | 2-fluoroadenosine | |
| | | 2ZJ1 | 3´-keto aristeromycin | |
| | | 3CE6 | Adenosine | |
| | | 3DHY | 5´-ethylthioadenosine | |
| *Naegleria fowleri* | Eukaryota, Percolozoa | 5V96 | Adenosine | Unpublished |
| *Plasmodium falciparum* | Eukaryota, Apicomplexa | 1V8B | Adenosine | Tanaka *et al.*, 2004[28] |
| *Pseudomonas aeruginosa* | Bacteria, Proteobacteria | 6F3M | Adenosine, zinc | Czyrko *et al.*, 2018[8] |
| | | 6F3N | SAH/adenosine, zinc | |
| | | 6F3O | Adenine, zinc | |
| | | 6F3P | 3´-deoxyadenosine | |
| | | 6F3Q | Adenine, rubidium | |
| *Rattus norvegicus* | Eukaryota, Animalia | 1B3R | None | Hu *et al.*, 1999[29] |
| | | 1K0U | Eritadenine | Huang *et al.*, 2002[30] |
| | | 1KY4 | None | Takata *et al.,*2002[31] |
| | | 2H5L | 3-deazaeritadenine | Yamada *et al.*, 2005[32] |
| *Synechocystis sp.* | Bacteria, Cyanobacteria | 7O5L | Adenosine, rubidium | Malecki *et al.,*2022[33] |
| | | 7O5M | Adenosine | |
| *Trypanosoma brucei* | Eukaryota, Euglenozoa | 3H9U | Adenosine | unpublished |
| *Thermotoga maritima* | Bacteria, Thermotogae | 3X2E | None (open) | Zheng *et al.*, 2015[34] |
| | | 3X2F | None (closed) | |
| | | 5TOV | NADH | Brzezinski *et al.*, 2017[35] |
| | | 5TOW | NADH, adenosine | |

**DNA sequences**

>*Sc*HSMT

ATGAAGCGCATTCCAATCAAAGAACTAATAGTTGAGCACCCCGGAAAAGTTCTTATCCTTGATGGTGGACAGGGTACAGA
ATTGGAAAACAGAGGCATTAACATAAATAGTCCGGTATGGTCTGCAGCTCCTTTTACGAGCGAATCCTTTTGGGAGCCATC
TTCTCAAGAGCGAAAGGTGGTAGAAGAAATGTACAGAGACTTTATGATTGCTGGCGCAAACATATTAATGACAATTACTT
ACCAGGCAAACTTTCAAAGCATATCTGAGAATACCTCGATTAAAACTCTGGCTGCTTACAAGCGTTTTCTCGATAAAATCG
TGTCATTTACTCGTGAATTTATTGGTGAGGAAAGGTACTTAATCGGGAGTATTGGCCCATGGGCAGCACATGTATCCTGTG
AATATACTGGTGACTATGGTCCCCATCCTGAGAATATTGATTACTACGGCTTTTTCAAACCCCAGCTGGAGAACTTCAACCA
AAATAGAGATATTGATCTTATTGGTTTTGAAACGATTCCAAATTTTCATGAGTTAAAGGCTATTTTATCCTGGGATGAAGAT
ATTATTTCGAAGCCCTTTTATATTGGGTTGTCGGTGGATGACAATAGTTTGCTACGAGACGGTACCACTTTGGAAGAAATT
TCTGTCCATATAAAAGGCCTCGGAAATAAAATTAACAAGAATCTCTTATTAATGGGAGTTAACTGTGTCAGTTTCAATCAA
TCGGCATTAATTCTTAAAATGTTGCACGAGCATCTACCTGGCATGCCTCTGCTAGTTTACCCAAACAGTGGAGAAATCTAC
AATCCCAAAGAGAAGACATGGCACCGGCCGACTAATAAGTTGGATGACTGGGAGACCACGGTTAAGAAATTCGTTGATA
ATGGTGCGCGCATTATTGGCGGTTGTTGTAGAACGTCTCCTAAAGATATCGCCGAAATTGCATCAGCTGTAGATAAATACT
CCTAA

>*Mj*DadD (codon-optimised for *E. coli*)

ATGATCCTGATCAAAAACGTGTTTGTGAATGGTAAACGCCAGGATATTCTGATCGAAGGCAACAAGATCAAAAAAATCGG
CGAGGTGAAAAAAGAAGAAATCGAAAACGCCGAAATCATCGACGGCAAAAACAAAATTGCCATTCCGGGTCTGATTAAC
ACCCATACACATATTCCGATGACACTGTTTCGTGGTGTTGCAGATGATCTGCCGCTGATGGAATGGCTGAATAACTATATT
TGGCCGATGGAAGCCAAACTGAACGAAGAAATTGTTTATTGGGGCACCCTGCTGGGTTGTATTGAAATGATTCGTAGCGG
CACCACCACCTTTAATGATATGTATTTTTTCCTGGAAGGGATCGCCAAAGCAGTTGATGAAAGCGGTATGCGTGCAGTTCT
GGCCTATGGTATGATTGACCTGTTTGATGAAGAACGTCGTGAACGCGAACTGAAAAATGCAGAGAAATACATCAACTATA
TCAACAGCCTGAACAACAGCCGTATTATGCCTGCACTGGGTCCGCATGCACCGTATACCTGTAGCAAAGAACTGTTAATG
GAAGTGAATAACCTGGCCAAAAAGTATAACGTGCCGATTCATATTCATCTGAACGAAACCCTGGATGAGATCAAGATGGT
TAAAGAAAAAACCGGTATGGAACCGTTCATCTATCTGAATAGCTTTGGCTTTTTTGATGATGTTCGTGCAATTGCCGCACA
TTGTGTTCATCTGACCGATGAAGAAATCAAGATCATGAAACAGAAAAACATTAACGTGTCGCATAACCCGATTAGCAATCT
GAAACTGGCAAGCGGTGTTGCACCGATTCCGAAACTGCTGGCCGAAGGTATTAATGTTACCCTGGGCACCGATGGTTGTG
GTAGCAATAATAACCTGAACCTGTTCGAAGAGATTAAAGTTAGCGCCATTCTGCATAAAGGCGTTAATCTGAATCCGACC
GTTGTTAAAGCAGAAGAAGCATTTAACTTCGCCACCAAAAATGGTGCAAAAGCCCTGAACATTAAAGCCGGTGAAATTCG
TGAAGGTTATCTGGCAGATATTGTGCTGATTAATCTGGATAAACCGTATCTGTACCCGAAAGAAAACATTATGAGCCATCT
GGTGTATGCCTTTAATGGCTTCGTGGATGATGTGATTATTGATGGCAACATTGTTATGCGTGATGGCGAAATTCTGACCGT
TGACGAAGAAAAAGTTTACGAAAAAGCCGAAGAGATGTATGAAATTCTGCGCAGCTAA

>*Cg*SAHH

ATGGCACAGGTTATGGACTTCAAGGTTGCCGATCTTTCACTAGCAGAGGCAGGACGTCACCAGATTCGTCTTGCAGAGTA
TGAGATGCCAGGTCTCATGCAGTTGCGCAAGGAATTCGCAGACGAGCAGCCTTTGAAGGGCGCCCGAATTGCTGGTTCTA
TCCACATGACGGTCCAGACCGCCGTGCTTATTGAGACCCTCACTGCTTTGGGCGCTGAGGTTCGTTGGGCTTCCTGCAACA
TTTTCTCCACCCAGGATGAGGCTGCAGCGGCTATCGTTGTCGGCTCCGGCACCGTCGAAGAGCCAGCTGGTGTTCCAGTA
TTCGCGTGGAAGGGTGAGTCACTGGAGGAGTACTGGTGGTGCATCAACCAGATCTTCAGCTGGGGCGATGAGCTGCCAA
ACATGATCCTCGACGACGGCGGTGACGCCACCATGGCTGTTATTCGCGGTCGCGAATACGAGCAGGCTGGTCTGGTTCCA
CCAGCAGAGGCCAACGATTCCGATGAGTACATCGCATTCTTGGGCATGCTGCGTGAGGTTCTTGCTGCAGAGCCTGGCAA
GTGGGGCAAGATCGCTGAGGCCGTTAAGGGTGTCACCGAGGAAACCACCACCGGTGTGCACCGCCTGTACCACTTCGCT
GAAGAAGGCGTGCTGCCTTTCCCAGCGATGAACGTCAACGACGCTGTCACCAAGTCCAAGTTTGATAACAAGTACGGCAC
CCGCCACTCCCTGATCGACGGCATCAACCGCGCCACTGACATGCTCATGGGCGGCAAGAACGTGCTTGTCTGCGGTTACG
GCGATGTCGGCAAGGGCTGCGCTGAGGCTTTCGACGGCCAGGGCGCTCGCGTCAAGGTCACCGAAGCTGACCCAATCAA

CGCTCTTCAGGCTCTGATGGATGGCTACTCTGTGGTCACCGTTGATGAGGCCATCGAGGACGCCGACATCGTGATCACCG
CGACCGGCAACAAGGACATCATTTCCTTCGAGCAGATGCTCAAGATGAAGGATCACGCTCTGCTGGGCAACATCGGTCAC
TTTGATAATGAGATCGATATGCATTCCCTGTTGCACCGCGACGACGTCACCCGCACCACGATCAAGCCACAGGTCGACGA
GTTCACCTTCTCCACCGGTCGCTCCATCATCGTCCTGTCCGAAGGTCGCCTGTTGAACCTTGGCAACGCCACCGGACACCC
ATCATTTGTCATGTCCAACTCTTTCGCCGATCAGACCATTGCGCAGATCGAACTGTTCCAAAACGAAGGACAGTACGAGAA
CGAGGTCTACCGTCTGCCTAAGGTTCTCGACGAAAAGGTGGCACGCATCCACGTTGAGGCTCTCGGCGGTCAGCTCACCG
AACTGACCAAGGAGCAGGCTGAGTACATCGGCGTTGACGTTGCAGGCCCATTCAAGCCGGAGCACTACCGCTACTAA

>*Ll*SAHH (codon-optimised for *E. coli*)

ATGGCCCTGCTGGTTGAAAAAACCACCAGTGGTCGTGAATATAAAGTGAAAGATATGAGCCAGGCAGATTTTGGTCGTCT
GGAAATTGAACTGGCCGAAGTTGAAATGCCTGGTCTGATGGCAAGCCGTAGCGAATTTGGTCCGAGCCAGCCGTTTAAA
GGTGCAAAAATTACCGGTAGCCTGCACATGACCATTCAGACCGCAGTTCTGATTGAAACCCTGACCGCACTGGGTGCCGA
AGTTCGTTGGTGTAGCTGTAACATTTTTAGCACCCAGGATCATGCAGCAGCAGCAATTGCACGTGATAGCGCAGCAGTTTT
TGCATGGAAAGGCGAAACCCTGCAAGAATATTGGTGGTGTACCGAACGTGCACTGGATTGGGGTCCTGGTGGTGGTCCG
GATCTGATTGTTGATGATGGTGGTGATACCACACTGCTGATTCATGAAGGTGTTAAAGCCGAAGAGATCTATGAAAAAAG
CGGTCAGTTTCCGGATCCTGATAGCACCGATAATGCAGAATTCAAAATTGTGCTGAGCATCATCAAAGAAGGCCTGAAAA
CCGATCCGAAACGCTATCACAAAATGAAAGATCGTGTTGTTGGTGTGAGCGAAGAAACCACCACCGGTGTTAAACGTCTG
TATCAGATGCAGGCAAATGGCACCCTGCTGTTTCCGGCAATTAATGTTAATGATAGCGTGACCAAAAGCAAATTTGATAAC
CTGTATGGTTGTCGTCATAGCCTGCCGGATGGCCTGATGCGTGCAACCGATGTTATGATTGCAGGTAAAGTTGCAGTTGTT
GCAGGTTATGGTGATGTTGGTAAAGGTTGTGCAGCAGCCCTGAAACAGGCAGGCGCACGTGTTATTGTTACCGAAATTG
ATCCGATTTGTGCACTGCAGGCAACCATGGAAGGTCTGCAGGTTCTGACCCTGGAAGATGTTGTTAGCGAAGCAGATATT
TTTGTTACCACCACGGGCAACAAAGATATCATTATGCTGGACCACATGAAAAAAATGAAAAACAACGCCATCGTGTGCAA
CATCGGCCATTTTGATAATGAGATTGATATGCTGGGCTTAGAAACCCATCCGGGTGTGAAACGTATTACCATTAAACCGCA
GACCGATCGTTGGGTTTTTCCTGAAACCAATACCGGCATTATTATCCTGGCGGAAGGTCGTCTGATGAATCTGGGTTGTGC
AACCGGTCATCCGAGCTTTGTTATGAGCTGTAGCTTTACCAATCAGGTTATTGCACAGCTGGAACTGTGGAATGAAAAATC
AAGCGGCAAATACGAGAAAAAGGTTTATGTTCTGCCGAAACACCTGGATGAAAAAGTTGCCGCACTGCATCTGGAAAAA
CTGGGTGCAAAACTGACCAAACTGAGCAAAGATCAGGCCGATTATATCAGCGTTCCGGTTGAAGGTCCGTATAAACCGTT
TCATTATCGCTACTAA

>*Mj*SIHH (codon-optimised for *E. coli*)

ATGTACGAGGTGCGCGATATCAACCTGTGGAAAGAAGGTGAACGTAAAATTCAGTGGGCAAAACAGCACATGCCGGTTC
TGAATCTGATTCGTGAACGTTTCAAAGAAGAGAAACCGTTTAAAGGCATTACCATTGGTATGGCACTGCATCTGGAAGCA
AAAACCGCAGTTCTGGCAGAAACCCTGATGGAAGGTGGTGCAGAAATTGCCATTACCGGTTGTAATCCGCTGAGCACCCA
GGATGATGTTGCAGCAGCATGTGCAAAAAAAGGTATGCATGTTTATGCATGGCGTGGTGAAACCGTGGAAGAATATTAT
GAAAACCTGAACAAAGTGCTGGATCACAAACCGGATATTGTGATTGATGATGGTTGCGATCTGATCTTTCTGCTGCATACC
AAACGTACCGAACTGCTGGATAACATTATGGGTGGTTGTGAAGAAACCACCACCGGTATTATTCGTCTGAAAGCAATGGA
AAAAGAAGGCGCACTGAAATTTCCGGTTATGGATGTTAATGATGCCTACACCAAACACCTGTTTGATAATCGTTATGGCAC
CGGTCAGAGCGCACTGGATGGTATTCTGCGTGCAACCAATCTGCTGATTGCAGGTAAAACCGTTGTTGTTGCAGGTTATG
GTTGGTGTGGTCGTGGTGTTGCAATGCGTGCAAAAGGTCTGGGTGCAGAAGTTGTTGTTACCGAAGTTAATCCGATTCGT
GCACTGGAAGCCCGTATGGATGGTTTTCGTGTTATGAAAATGGAAAAGGCAGCCGAAATTGGCGATATCTTTATTACAAC
CACCGGTTGCAAAGATGTGATCCGCAAAGAACATATTCTGAAAATGCGTAATGGTGCCATTCTGGCAAATGCAGGTCATT
TTGATAACGAGATCAACAAGAAACACCTGGAAGAACTGGCCAAAAGCATTAAAGAAGTTCGTAATTGCGTCACCGAATAT
GATCTGGGCAACAAAAAAATCTATCTGCTTGGTGAAGGTCGTCTGGTTAATCTGGCATGTGCAGATGGTCATCCGTGTGA
AGTTATGGATATGAGCTTTGCAAATCAGGCACTGGCAGCAGAATATATCCTGAAAAATCACGAAAAACTGGAACCGCGTG
TTTATAACATTCCGTATGAACAGGATCTGATGATTGCCAGCCTGAAACTGAAAGCCATGGGTATTGAAATTGATGAGCTG
ACCAAAGAGCAGAAGAAGTATCTGGAAGATTGGCGTGAAGGCACCTAA

>*Mma*SAHH (codon-optimised for *E. coli*)

ATGAGCAACGTGAAAGATATGAGCCTGGCACCGAGCGGTCATCTGAAAATGGAATGGGCAAAACGTCACATGCCGGTTC
TGTGTCGTATTGCCGAAGAATTCAAAAACGATAAACCGTTTGAAGGTCTGACCATTGGTATGGCACTGCATCTGGAAGCA
AAAACCGCAATTCTGGCAGAAACCCTGCTGGAAGGTGGTGCAAAAATTGTTATTACCGGTTGTAATCCGCTGAGCACCCA
GGATGATGTTGCAGCAGCATGTGTTGAAAAAGGTATGGAAGTTTATGCATGGCGTGGTGAAACCAACGAAGAATATTAT
GAGAACCTGAACAAAGTGCTGGATAGCAATCCGGATATCATTATTGATGATGGTGCCGATCTGATCTTTCTGATTCATACC
GAACGTACCGAACTGATTGGCAAAATTATGGGTGGTTGTGAAGAAACCACCACCGGTATTATTCGTCTGAAAAGCATGGC
CGAAGAAGGTGCACTGAAATTTCCGGTTGTTAATGTGAATGATGCCTACACCAAACACCTGTTTGATAATCGTTATGGCAC
CGGTCAGAGCGCAATGGATGGCATTATTCGTACCACCAATCTGCTGATTGCAGGTAAAAATGTTGTGGTTGGTGGTTATG
GTTGGTGTGGTCGTGGTGTTGCAAGCCGTGCAGCAGGTCATGGTGCAAATGTGATTATTACCGAAGTTAATCCGATTCGT
GCACTGGAAGCCAAGATGGATGGTTTTACCGTGTTAAAAATGGAAGAGGCAGCCAAAATTGGCGATATTTTTGTTACCAC
CACAGGCTGCAAAGATATTCTGCGTATGGAACATTTTCTGCTGATGAAAGATGGTGCAGTTCTGAGCAATGCAGGCCATT
TTGATAACGAGATTAACAAAAACGACCTGAAAGAGCTGAGCAAGAGCGTTAAAGAAGCACGCTTTAACATCGAAGAGTA
TGATCTGGGCAACAAAAAGATTTATCTGCTTGGTGAAGGTCGTCTGGTTAATCTGGCATGTGCAGATGGTCATCCGTGTG
AAGTTATGGATATGAGCTTTGCAAATCAGGCACTGAGCGCCAAATTCATCAAAGAAAATAAAGGCAAGCTGGAAAACGA
GGTGTATGAAATTCCGTATGAGCAGGATTTCAAAATCGCACTGCTGAAACTGCATAGCATGGGTGCAGATATTGATGAAC
TGAGTCCGGAACAGCGTAAATATCTGAGCGATTGGAAAGAAGGCACCTAA

>*Mm*SAHH (codon-optimised for *E. coli*)

ATGAGCGACAAACTGCCGTATAAAGTTGCAGATATTGGTCTGGCAGCATGGGGTCGTAAAGCACTGGATATTGCAGAAA
ATGAAATGCCTGGTCTGATGCGTATGCGTGAAATGTATAGCGCAAGCAAACCGCTGAAAGGTGCACGTATTGCAGGTTGT
CTGCACATGACCGTTGAAACCGCAGTTCTGATTGAAACCCTGGTTGCACTGGGTGCAGAAGTTCGTTGGAGCAGCTGTAA
CATTTTTAGCACCCAGGATCATGCAGCAGCAGCAATTGCAAAAGCAGGTATTCCGGTTTTTGCATGGAAAGGTGAAACCG
ATGAAGAATATCTGTGGTGTATTGAACAGACCCTGCACTTTAAAGATGGTCCGCTGAATATGATTCTGGATGATGGTGGT
GATCTGACCAATCTGATTCATACCAAATATCCGCAGCTGCTGAGCGGTATTCGTGGTATTAGCGAAGAAACCACCACCGGT
GTTCATAACCTGTATAAAATGATGAGCAACGGCATTCTGAAAGTTCCGGCAATTAATGTTAATGATAGCGTGACCAAAAG
CAAATTCGATAATCTGTATGGTTGTCGCGAAAGCCTGATTGATGGTATTAAACGTGCAACCGATGTTATGATTGCAGGTAA
AGTTGCCGTTGTTGCAGGTTATGGTGATGTTGGTAAAGGTTGTGCACAGGCACTGCGTGGTTTTGGTGCACGTGTGATTA
TTACCGAAATTGATCCGATTAATGCACTGCAGGCAGCAATGGAAGGCTATGAAGTTACCACCATGGATGAAGCATGTAAA
GAAGGCAACATTTTTGTTACAACCACCGGTTGCGTTGATATCATTCTGGGTCGTCATTTTGAGCAGATGAAAGATGATGCC
ATTGTGTGCAATATCGGCCATTTTGATGTTGAGATTGATGTGAAATGGCTGAATGAAAACGCCGTGGAAAAAGTGAACAT
TAAACCGCAGGTTGATCGCTATTGGCTGAAAAATGGTCGTCGTATTATTCTGCTGGCAGAAGGTCGTCTGGTTAATCTGG
GTTGTGCAATGGGTCATCCGAGCTTTGTTATGAGCAATAGCTTTACCAATCAGGTGATGGCACAGATTGAACTGTGGACC
CATCCGGATAAATATCCGGTTGGTGTTCATTTCCTGCCGAAAAAACTGGATGAGGCAGTTGCAGAAGCACATCTGGGTAA
ACTGAATGTGAAACTGACCAAACTGACAGAAAAACAGGCACAGTATCTGGGTATGCCGATTAACGGTCCGTTTAAACCGG
ATCATTATCGCTATTAA

>*Pa*SAHH (codon-optimised for *E. coli*)

ATGAGCGCAGTTATGACACCGGCAGGTTTTACCGATTATAAAGTTGCAGATATTACCCTGGCAGCATGGGGTCGTCGTGA
ACTGATTATTGCAGAAAGCGAAATGCCTGCACTGATGGGTCTGCGTCGTAAATATGCAGGTCAGCAGCCGCTGAAAGGT
GCAAAAATTCTGGGTTGTATTCACATGACCATTCAGACCGGTGTTCTGATTGAAACCCTGGTTGCACTGGGTGCAGAAGTT
CGTTGGAGCAGCTGTAACATTTTTAGCACCCAGGATCAGGCAGCAGCAGCAATTGCAGCAGCCGGTATTCCGGTTTTTGC
ATGGAAAGGTGAAACCGAAGAGGAATATGAATGGTGTATTGAACAGACCATTCTGAAAGATGGTCAGCCGTGGGATGCA
AATATGGTTCTGGATGATGGTGGTGATCTGACCGAAATTCTGCATAAAAAGTATCCGCAGATGCTGGAACGTATTCATGG
TATTACCGAAGAAACCACCACCGGTGTGCATCGTCTGCTGGATATGCTGAAAAATGGCACCCTGAAAGTTCCGGCAATTA
ATGTTAATGATAGCGTGACCAAAAGCAAAAACGACAACAAATATGGCTGTCGCCATAGCCTGAATGATGCAATTAAACGT
GGCACCGATCATCTGCTGAGCGGTAAACAGGCACTGGTTATTGGTTATGGTGATGTTGGTAAAGGTAGCAGCCAGAGCC
TGCGTCAAGAAGGTATGATTGTTAAAGTTGCCGAAGTTGATCCGATTTGTGCAATGCAGGCATGTATGGATGGTTTTGAA

GTTGTTAGCCCGTATAAAAACGGCATTAATGATGGCACCGAAGCAAGCATTGATGCAGCACTGCTGGGTAAAATTGATCT
GATTGTTACCACCACGGGTAATGTGAATGTTTGTGATGCAAACATGCTGAAAGCCCTGAAAAAACGTGCAGTTGTTTGCA
ACATTGGCCACTTTGATAACGAAATTGATACCGCCTTTATGCGCAAAAATTGGGCATGGGAAGAAGTTAAACCGCAGGTT
CATAAAATTCATCGTACCGGTAAAGATGGCTTTGATGCCCATAATGATGATTATCTGATTCTGCTGGCAGAAGGTCGTCTG
GTTAATCTGGGTAATGCAACCGGTCATCCGAGCCGTATTATGGATGGCAGCTTTGCAAATCAGGTGCTGGCACAGATTCA
CCTGTTTGAACAGAAATATGCCGATCTGCCTGCAGCAGAAAAAGCCAAACGTCTGAGCGTTGAAGTTCTGCCGAAAAAAC
TGGATGAAGAGGTTGCCCTGGAAATGGTTAAAGGTTTTGGTGGTGTTGTTACCCAGCTGACCCCGAAACAGGCAGAATAT
ATCGGTGTTAGCGTGGAAGGTCCGTTTAAACCGGATACCTATCGCTATTAA

>*Pfu*SAHH (codon-optimised for *E. coli*)

ATGGATTGCGGCAAAGATTATTGCGTTAAAGATCTGAGCCTGGCAGAAGAAGGTTGGAAAAAAATCGATTGGGTTAGCC
GTTTTATGCCGGTTCTGCAGTATATCAAACGCGAATTCGAAGAGAAAAAACCGTTTAAAGGTGTTCGTATTGCAGCAACCC
TGCATCTGGAAATGAAAACCGCATTTCTGCTGCTGACCCTGAAAGCCGGTGGTGCAGAAGTTAGCGCAGCAGCAAGCAAT
CCGCTGAGCACCCAGGATGATGTTGTTGCAGCACTGGCAAAAGCGGGTGTTAAAGTTTATGCAATTCGTGGTGAAAGCC
GTGAGCAGTATTATGAGTTCATGCATAAAGCACTGGATATCCGTCCGAACATCATTATTGATGATGGTGCAGATATGATCA
GCCTGGTTCATAAAGAACGTCAAGAAATGCTGGATGAAATTTGGGGTGGTAGCGAAGAAACCACCACCGGTGTTATTCGT
CTGCGTGCAATGGAAAAAGCAGGCATTCTGAAATTTCCGGTTATTGCCGTTAACGACAGCTACATGAAATACCTGTTTGAT
AATCGTTATGGCACCGGTCAGAGCACCTGGGATGGTATTATGCGTGCAACCAATCTGCTGATTGCAGGTAAAAATGTTGT
GGTGGTTGGTTATGGTTGGTGTGGTCGTGGTATTGCAATGCGTGCACGTGGTCTGGGTGCAACCGTTATTGTTGTTGAAG
TTGATCCGATTAAAGCCCTGGAAGCACGTATGGATGGTTTTCTGGTTATGGATATGAAAGAGGCAGCAAAAATCGGCGAT
ATTTTTGTTACCGCAACCGGCAACATTAAATGCATTCGTCGTGAACATTTCGAGCTGATGAAAGATGGTGCAATTATGGCA
AATGCCGGTCATTTTGATGTGGAAATTTGGAAACCGGATCTGGAAAAACTGGCCGTGGAAATCAATAATCCGCGTCCGAA
TGTTACCGAGTATAAACTGAAAGACGGTCGTCGTCTGTATCTGCTGGCAGATGGTCGTCTGGTTAATCTGGTTGCAGCCG
ATGGTCATCCGGCAGAAATCATGGATATGTCATTTGCACTGCAGGCCAAAGCAGCCGAATATATCAAAGATAATCATGAA
CGTCTGGAACCGAAGGTGTATATTCTGCCTCGTGAAATTGATGAAATGGTGGCACGTATTAAACTGGAAAGCATGGGCAT
TAAAATCGAAGAACTGACCGAAGAACAGAAAAAGTATCTGGAAAGTTGGGAACATGGCACCTAA

>*Sac*SAHH (codon-optimised for *E. coli*)

ATGGATTATCGCGTTAAAGATCTGAGCCTGGCAGAACAGGGTCGTAAACAAATTGAATGGGCAGAACTGCACATGCCTG
CACTGATGGAAATTCGTAAACGTTTTAATGCAGAGAAACCGCTGGATGGTATTCGTATTGGTGCCGTTCTGCATGTTACCA
AAGAAACCGCAGTTCTGGTTGAAACCCTGAAAGCCGGTGGTGCAGAAATTGCACTGGCAGGTAGCAATCCGCTGAGCAC
CCAGGATGATGTTGCAGCAGGTCTGGCAAAAAATGGTATTCATGTTTATGCATGGCGTGGCGAAACCGAGAAAGATTATT
ATGATAACATTCGCGAAATCCTGAAATATGAACCGCATGTGATTATGGATGATGGTGGTGATCTGCATGCCTATGTGCAT
GAAAATAATCTGACCAGCAAAATTGTTGGTGGCACCGAAGAAACCACCACCGGTGTTATTCGTCTGAAAGCCATGGAAGA
AGAGAAAGTTCTGAAATATCCGGTGATTGCCGTGAATAATGCCTTTACCAAATACCTGTTTGATAACCGTATTGGCACCGG
TCAGAGCACCATTGATGGCATTCTGCGTGCAACCAATATTCTGATTGCAGGTAAAGTTGCCGTGGTGATTGGTTATGGTTG
GGTTGGTCGTGGTATTGCAAGCCGTTTTAAAGGTATGGGTGCACGTGTTATTGTTGTTGAAAGCAGCCCGTTTCGTGCACT
GGAAGCCCTGATGGATGGTTTTGATGTTATGACCATGAATCGTGCAAGCGAAATTGGCGATATTTTTGTTACCGCAACCG
GTAATCTGAATGTTGTTAGCCGTGATCATATTCTGCGTATGAAAGATGGTGCGGTTCTGGCAAATAGCGGTCACTTTAATG
TTGAGATTGATGTGAAAGGCCTGAAAGAAATTAGCGTGGAAACCCGTGAAGTTCGTCAGAATCTGGAAGAATATAAACT
GCGTAATGGCAAACGCATTTATCTGCTGGCAGATGGTCGTCTGGTTAATCTGGTTGCAGCCGAAGGTCATCCGAGCGAAG
TTATGGATCTGAGCTTTTGTAATCAGGCACTGAGCGTTGAACATCTGATTAAAAACAAAGGCAAACTGGAAAACAAAGTG
TACAACGTGCCGATCGAAATTGATGAACAGGTTGCACGTCTGAAACTGAAAGCACTGGGTATTGAAATTGAAGAACTGAC
CATCGAGCAGAAAGAATACATCAAACAGTGGAAATACGGCACCTAA

>*Sa*SAHH

ATGGCAAGCGCCCAGCAGCACGACTTCAAGGTCGCCGACCTCTCCCTCGCGGAGTTCGGCCGCAAGGAGATCACCCTCGC
CGAGCACGAGATGCCCGGCCTGATGTCGATCCGCGAGGAGTACGCCGCGTCCCAGCCGCTGGCCGGCGCCCGCGTCACC

GGCTCGCTGCACATGACCGTCCAGACGGCCGTCCTCATCGAGACGCTCACCGCCCTGGGCGCGGAGGTCCGCTGGGCCTC
CTGCAACATCTTCTCCACCCAGGACCACGCCGCCGCCGCCATCGCGGTCGGCCCCAACGGCACCCCGGACAACCCGCAGG
GCGTCCCGGTCTTCGCCTGGAAGGGCGAGAGCCTGGAGGAGTACTGGTGGTGCACCGAGCAGGCGCTGACCTGGCCGA
ACACCCCCACCGGCGGCCCCAACATGATCCTGGACGACGGCGGTGACGCCACCCTCCTCGTCCACAACGGCGTCCAGTAC
GAGAAGGACGGCAAGGTCCCCGACCCGGTCACCGCCGAGTCCGACGAGCACCGCGTCATCCTCCAGCTGCTGACCCGCA
CCCTCGGCGAGAACCCGCAGAAGTGGACCCAGCTCGCCTCGGAGATCCGCGGCGTCACCGAGGAGACCACCACCGGCGT
CCACCGCCTCTACGAGATGCAGCGCGACGGCCAGCTGCTCTTCCCGGCGATCAACGTCAACGACGCGGTCACCAAGTCGA
AGTTCGACAACAAGTACGGCTGCCGCCACTCCCTGATCGACGGCATCAACCGTGCCACCGACGTCCTCATCGGCGGCAAG
ACCGCCGTCGTCTGCGGCTACGGCGACGTCGGCAAGGGCTGCGCCGAGTCCCTGCGCGGCCAGGGCGCCCGCGTCATCG
TCACCGAGATCGACCCGATCTGCGCCCTCCAGGCGGCGATGGACGGCTACCAGGTCGCCACCCTCGACGACGTGATCGG
CCAGGCCGACATCTTCGTCACCACGACCGGCAACAAGGACATCATCATGGCCGCCGACATGGCCAAGATGAAGCACCAG
GCCATCGTGGGGAACATCGGCCACTTCGACAACGAGATCGACATGGCCGGCCTCGCCGCCATCCCCGGCATCGTCAAGG
ACGAGGTCAAGCCGCAGGTCCACACCTGGACCTTCCCCGACGGCAAGGTCATCATCGTGCTGTCCGAGGGCCGCCTGCTC
AACCTGGGCAACGCCACCGGCCACCCGTCCTTCGTGATGTCCAACAGCTTCGCGGACCAGACGCTGGCCCAGATCGAGCT
GTTCACCAAGCCCGACGAGTACCCGACCGACGTCTACGTGCTGCCCAAGCACCTCGACGAGAAGGTCGCCCGCCTCCACC
TCGACGCCCTCGGCGTCAGGCTGACGACCCTCCGCCCGGAGCAGGCCGCGTACATCGGCGTCTCCGTCGAAGGCCCGTTC
AAGCCGGACCACTACCGGTACTGA

>*Sf*SAHH

ATGACGACGACCTCCACGACCGGCCATGACTTCAAGGTCGCGGACCTCTCTCTGGCCGCTTTCGGCCGCAAGGAGATCAC
GCTGGCCGAGCACGAGATGCCCGGCCTGATGGCGATCCGCAAGGAGTACTCCGCGGAGAAGCCGCTGGCCGGAGCGCG
CATCACGGGCTCCCTGCACATGACGGTGCAGACGGCCGTGCTCATCGAAACCCTCGTCGCCCTCGGCGCCGAGGTCCGCT
GGGCCTCCTGCAACATCTTCTCCACCCAGGACCACGCCGCCGCGGCCATCGCCGTCGGCCCCGACGGCACCCCGGACAAC
CCGCGGGGCGTCCCGGTCTTCGCCTGGAAGGGCGAGACCCTGGAGGAGTACTGGTGGTGCACGGAACAGGCCCTCACCT
GGCCGAACACGCCCACCGGCGGCCCCAACATGATTCTCGACGACGGTGGTGACGCCACCCTCCTCGTCCACAAGGGCGTC
GAGTACGAGAAGGCCGGTGCCGCCCCCTCGGTCGACACCGCCGAGAACGACGAGCACCGCGTCATCCTCCAGCTCCTCA
ACCGCACCCTCGCCGAGAGCCCGCAGAAGTGGACGCAGCCCGCGTCGGAGATCCGCGGCGTCACCGAGGAGACCACCAC
CGGCGTCCACCGCCTCTACGAGATGCAGCAGGCCGGCACGCTGCTCTTCCCGGCGATCAACGTCAACGACGCCGTCACCA
AGTCGAAGTTCGACAACAAGTACGGCTGCCGCCACTCCCTGATCGACGGCATCAACCGCGCCACCGACGTCCTCATCGGC
GGCAAGACCGCCGTCGTCTGCGGCTACGGCGATGTCGGCAAGGGCTGCGCCGAGTCCCTGCGCGGCCAGGGCGCCCGG
GTCATGATCACTGAGATCGACCCGATCTGCGCCCTCCAGGCGGCGATGGACGGCTACCAGGTGGTGCGGCTGGACGATG
TCGTCGAGACCGCCGACATCTTCATCACCACCACGGGCAACAAGGACATCATCATGGCCTCGGACATGGCCAAGATGAAG
CACCAGGCCATCGTCGGCAACATCGGCCACTTCGACAACGAGATCGACATGGCCGGTCTCGCCGCCATCGACGGCATCGT
CAAGGACGAGGTCAAGCCGCAGGTCCACACCTGGACCTGGCCGGACGGCAAGAGCATCATCGTCCTGTCCGAGGGCCGC
CTGCTGAACCTGGGCAACGCCACCGGGCACCCCTCGTTCGTGATGTCGAACAGCTTCGCGAACCAGACGATCGCCCAGAT
CGAACTGTTCACCAAGCCGGAGTCGTACCCGACCGACGTCTACGTGCTGCCCAAGCACCTCGACGAGAAGGTCGCCCGCC
TCCACCTCGACGCCCTCGGCGCCAAGCTGACCACGCTCCGCCCGGAGCAGGCGGCGTACATCGGCGTCCCGGTCGAGGG
TCCCTACAAGCCGGACCACTACCGCTACTGA

>*Sso*SAHH (codon-optimised for *E. coli*)

ATGAGCTACAAAATCAAAGATCTGAGCCTGGCAAGCGAAGGTAAAAAACAAATTGAATGGGCAGAACGTCACATGCCGA
CACTGATGGAAATTCGTAAACGTTTTAAAGCCGAGAAACCGCTGAAAGGCATTAACATTAGCGCAGTTCTGCATGTTACC
AAAGAAACCGCAGCACTGGTTAAAACCCTGAAAATTGGTGGTGCAAATGTTGCACTGGCAGGTAGCAATCCGCTGAGCA
CCCAGGATGATGTTGCAGCAGCCCTGGTTGAAGAAGGTATTAGCGTTTTTGCATGGAAAGGCGAAAATGAAACCGAGTA
TTACAGCAACATTGAGAGCATCGTGAAAATCCATGAACCGAACATTGTTATGGATGATGGTGCCGATCTGCATGCCTATAT
TCATGAAAAAGTTAGCAGCAAGCTGGATATTTATGGTGGCACCGAAGAAACCACCACCGGTGTTATTCGTCTGAAAGCAA
TGGAAAAAGATGGCGTTCTGAAATATCCGCTGGTTGCAGTTAATAACGCCTATACCAAATACCTGTTCGATAATCGTTATG

GCACCGGTCAGAGCGCAATTGATGGTATTCTGCGTGCAACCAATATTCTGATTGCAGGTAAAATTGCAGTGGTTGCAGGT
TATGGTTGGGTTGGTCGTGGTATTGCAAATCGTCTGCGTGGTATGGGTGCACGTGTTATTGTTACCGAAGTTGATCCGATT
CGTGCACTGGAAGCAGTGATGGATGGTTTTGATGTTATGCCGATTGCCGAAGCAAGCAAAGTTGGTGATATTTTTGTTAC
CGCAACCGGTAATACCAAAGCCATTCGTGTTGAACACATGCTGAATATGAAAGATGGTGCCATTCTGAGCAATGCCGGTC
ACTTTAATGTTGAAGTTGATGTGAAAGGCCTGAAAGAAACAGCAGTTAAAGTGCGTAATATTCGTCCGTATGTGGATGAA
TATACCCTGCCGAATGGTAAACGTGTTTATCTGCTGGCAGATGGTCGTCTGGTTAATCTGGCAGCAGCAGAAGGTCATCC
GAGCGAAGTTATGGATATGAGCTTTGCAAATCAGGCACTGGCCGTTGAATATCTGGTGAAAAATCGTGGTAAGCTGGAA
AAAAAGGTGTACAATATGCCGATGGAACTGGATTATGAAGTGGCACGTATTAAGCTGAAAAGCATGGGTATTCAGATTG
ATGAACTGACCGAAGAACAGAAAGAATACCTGGAACAGTGGAAAAGCGGCACCTAA

>*Tk*SAHH

ATGGACTGCACGAAGGATTACTGCGTTAAGGACATCTCCCTGGCACCGAGCGGGGAGAAGAAGATAGACTGGGTCTCCC
GCTTCATGCCGGTTCTCCAGCACATCAGGAAGGACTTTGAAGAGAGGAAACCGTTTAAGGGCGTTAGGATAGCAGCGAC
TCTACACCTTGAGATGAAGACTGCCTTTCTGCTTCTGACGCTGAAAGCGGCTGGAGCCGAGGTTTCGGCAGCTGCCAGCA
ACCCACTCTCCACCCAGGACGATGTAGTTGCCGCTCTGGCAAAGGCGGGGGTCAAGGTCTACGCTATAAGGGGAGAGGA
CAGGGAGCAGTACTACGAGTTCATGCACAAGGCCCTCGACGTAAAACCGAACATCATCATAGACGACGGAGCGGATATG
GTGAGCACGGTTTTGAAGGAGAGGCAGGAGCTGATTCCCGAAATATGGGGGGCAAGCGAGGAAACCACAACCGGCGTC
ATAAGGCTCCGTGCCATGGAGAAGGATGGCGTCCTCAAGTTCCCGATCATAGCGGTCAACGATTCCTACACCAAATACCT
CTTCGACAACCGCTATGGAACCGGTCAGTCCACCTGGGACGGCATCATAAGGACTACTAACCTCCTCGTCGCTGGAAAGA
ACGTCGTTGTTGTCGGCTATGGCTGGTGCGGCAGGGGCATAGCAATGCGCGCGAGGGGACTTGGAGCGACCGTTATCGT
CGTTGAGGTTGACCCAATAAGGGCTCTAGAAGCCAGAATGGACGGATTCCTCGTCATGGACATGATGGAGGCGGCGAAG
GTAGGGGACATCTTCATAACTGCCACCGGAGACATCAACTGCATAAGGAAGGAGCACTTCGAGCTCATGAAGGACGGAG
CTATTCTCGCCAACGCCGGCCACTTCGATGTCGAGATTTCAAAGCCTGACCTTGAGGCCCTCGCAGTTGAGATAAGCGAG
CCAAGGCCGAACATCACAGAGTACAAAATGGCAGACGGGAGGAGGCTCTACCTCCTGGCTGAGGGCAGGCTTGTGAATC
TAGCCGCTGCCGACGGTCATCCAGCGGAGATAATGGACATGAGCTTCGCGCTCCAGGCGAAAGCCGCTGAGTACATCAA
GGAGAACCGCGGAAGGCTTGAGCCGAAGGTCTACGTTCTTCCGAGGGAGATAGACGAGATGGTGGCGAGGATAAAGCT
CGCCTCGATGGGGATAAAAATTGAGGAACTCACAGAAGAGCAAAAGAAATATCTGGAAAGCTGGGAGCACGGCACCTG
A

>*Tm*SAHH (codon-optimised for *E. coli*)

ATGAACACCGGTGAGATGAAGATTAATTGGGTTAGCCGTTATATGCCGCTGCTGAACAAAATTGCCGAAGAATATAGCCG
TGAAAAACCGCTGAGTGGTTTTACCGTTGGTATGAGCATTCATCTGGAAGCAAAAACCGCATATCTGGCAATTACCCTGA
GCAAACTGGGTGCAAAAGTTGTTATTACCGGTAGCAATCCGCTGAGCACCCAGGATGATGTTGCAGAAGCACTGCGTAGC
AAAGGTATTACCGTTTATGCACGTCGTACCCATGATGAAAGCATTTATCGTGAAAACCTGATGAAAGTGCTGGATGAACG
TCCGGATTTCATTATTGATGATGGTGGTGATCTGACCGTTATTAGCCATACCGAACGTGAAGAAGTTCTGGAAAATCTGAA
AGGTGTTAGCGAAGAAACCACCACCGGTGTGCGTCGTCTGAAAGCACTGGAAGAAACCGGTAAACTGCGTGTTCCGGTT
ATTGCAGTTAATGACAGCAAAATGAAATACCTGTTCGATAATCGTTATGGCACCGGTCAGAGCACCTGGGATGCAATTAT
GCGTAATACCAATCTGCTGGTTGCCGGTAAAAATGTTGTTGTTGCAGGTTATGGTTGGTGTGGTCGTGGTATTGCACTGC
GTGCAGCAGGTCTGGGTGCACGTGTTATTGTTACCGAAGTTGATCCGGTTAAAGCAGTTGAAGCAATTATGGATGGCTTT
ACCGTTATGCCGATGAAAGAAGCAGTTAAAATTGCGGATTTTGTGATTACCGCAAGCGGCAATACCGATGTTCTGAGCAA
AGAAGATATTCTGAGCCTGAAAGATGGTGCAGTTCTGGCAAATGCAGGTCATTTTAATGTTGAAATTCCGGTTCGTGTGCT
GGAAGAGATTGCAGTTGAAAAATTTGAAGCACGTCCGAATGTTACCGGTTATACCCTGGAAAATGGTAAAACCGTTTTTC
TGCTGGCAGAAGGTCGTCTGGTTAATCTGGCAGCCGGTGATGGTCATCCGGTTGAAATCATGGATCTGAGCTTTGCACTG
CAGATTTTTGCCGTTCTGTATCTGCTGGAAAACCATCGTAAAATGAGCCCGAAAGTTTATATGCTGCCGGATGAAATTGAT
GAACGTGTTGCACGTATGAAACTGGATAGCCTGGGTGTTAAAATCGATGAACTGACCGAAAAACAGCGTCGTTATCTGCG
TAGCTGGCAGTAA

### Protein sequences

His$_6$-tags are underlined.

>*Sc*HSMT

MGSSHHHHHHSSGLVPRGSHMKRIPIKELIVEHPGKVLILDGGQGTELENRGININSPVWSAAPFTSESFWEPSSQERKVVEE
MYRDFMIAGANILMTITYQANFQSISENTSIKTLAAYKRFLDKIVSFTREFIGEERYLIGSIGPWAAHVSCEYTGDYGPHPENIDYY
GFFKPQLENFNQNRDIDLIGFETIPNFHELKAILSWDEDIISKPFYIGLSVDDNSLLRDGTTLEEISVHIKGLGNKINKNLLLMGVNC
VSFNQSALILKMLHEHLPGMPLLVYPNSGEIYNPKEKTWHRPTNKLDDWETTVKKFVDNGARIIGGCCRTSPKDIAEIASAVDK
YS

>*Cg*SAHH

MGSSHHHHHHSSGLVPRGSHMAQVMDFKVADLSLAEAGRHQIRLAEYEMPGLMQLRKEFADEQPLKGARIAGSIHMTVQT
AVLIETLTALGAEVRWASCNIFSTQDEAAAAIVVGSGTVEEPAGVPVFAWKGESLEEYWWCINQIFSWGDELPNMILDDGGD
ATMAVIRGREYEQAGLVPPAEANDSDEYIAFLGMLREVLAAEPGKWGKIAEAVKGVTEETTTGVHRLYHFAEEGVLPFPAMNV
NDAVTKSKFDNKYGTRHSLIDGINRATDMLMGGKNVLVCGYGDVGKGCAEAFDGQGARVKVTEADPINALQALMDGYSVV
TVDEAIEDADIVITATGNKDIISFEQMLKMKDHALLGNIGHFDNEIDMHSLLHRDDVTRTTIKPQVDEFTFSTGRSIIVLSEGRLLN
LGNATGHPSFVMSNSFADQTIAQIELFQNEGQYENEVYRLPKVLDEKVARIHVEALGGQLTELTKEQAEYIGVDVAGPFKPEHY
RY

>*Ll*SAHH

MGSSHHHHHHSSGLVPRGSHMALLVEKTTSGREYKVKDMSQADFGRLEIELAEVEMPGLMASRSEFGPSQPFKGAKITGSLH
MTIQTAVLIETLTALGAEVRWCSCNIFSTQDHAAAAIARDSAAVFAWKGETLQEYWWCTERALDWGPGGGPDLIVDDGGDT
TLLIHEGVKAEEIYEKSGQFPDPDSTDNAEFKIVLSIIKEGLKTDPKRYHKMKDRVVGVSEETTTGVKRLYQMQANGTLLFPAINV
NDSVTKSKFDNLYGCRHSLPDGLMRATDVMIAGKVAVVAGYGDVGKGCAAALKQAGARVIVTEIDPICALQATMEGLQVLTL
EDVVSEADIFVTTTGNKDIIMLDHMKKMKNNAIVCNIGHFDNEIDMLGLETHPGVKRITIKPQTDRWVFPETNTGIIILAEGRL
MNLGCATGHPSFVMSCSFTNQVIAQLELWNEKSSGKYEKKVYVLPKHLDEKVAALHLEKLGAKLTKLSKDQADYISVPVEGPYK
PFHYRY

>*Mj*DadD

MGSSHHHHHHSSGLVPRGSHMILIKNVFVNGKRQDILIEGNKIKKIGEVKKEEIENAEIIDGKNKIAIPGLINTHTHIPMTLFRGVA
DDLPLMEWLNNYIWPMEAKLNEEIVYWGTLLGCIEMIRSGTTTFNDMYFFLEGIAKAVDESGMRAVLAYGMIDLFDEERRERE
LKNAEKYINYINSLNNSRIMPALGPHAPYTCSKELLMEVNNLAKKYNVPIHIHLNETLDEIKMVKEKTGMEPFIYLNSFGFFDDVR
AIAAHCVHLTDEEIKIMKQKNINVSHNPISNLKLASGVAPIPKLLAEGINVTLGTDGCGSNNNLNLFEEIKVSAILHKGVNLNPTVV
KAEEAFNFATKNGAKALNIKAGEIREGYLADIVLINLDKPYLYPKENIMSHLVYAFNGFVDDVIIDGNIVMRDGEILTVDEEKVYE
KAEEMYEILRS

>*Mj*SIHH

MGSSHHHHHHSSGLVPRGSHMYEVRDINLWKEGERKIQWAKQHMPVLNLIRERFKEEKPFKGITIGMALHLEAKTAVLAETL
MEGGAEIAITGCNPLSTQDDVAAACAKKGMHVYAWRGETVEEYYENLNKVLDHKPDIVIDDGCDLIFLLHTKRTELLDNIMGG
CEETTTGIIRLKAMEKEGALKFPVMDVNDAYTKHLFDNRYGTGQSALDGILRATNLLIAGKTVVVAGYGWCGRGVAMRAKGL
GAEVVVTEVNPIRALEARMDGFRVMKMEKAAEIGDIFITTTGCKDVIRKEHILKMRNGAILANAGHFDNEINKKHLEELAKSIKE
VRNCVTEYDLGNKKIYLLGEGRLVNLACADGHPCEVMDMSFANQALAAEYILKNHEKLEPRVYNIPYEQDLMIASLKLKAMGIE
IDELTKEQKKYLEDWREGT

>*Mma*SAHH

MGSSHHHHHHSSGLVPRGSHMSNVKDMSLAPSGHLKMEWAKRHMPVLCRIAEEFKNDKPFEGLTIGMALHLEAKTAILAET
LLEGGAKIVITGCNPLSTQDDVAAACVEKGMEVYAWRGETNEEYYENLNKVLDSNPDIIIDDGADLIFLIHTERTELIGKIMGGCE

ETTTGIIRLKSMAEEGALKFPVVNVNDAYTKHLFDNRYGTGQSAMDGIIRTTNLLIAGKNVVVGGYGWCGRGVASRAAGHGA
NVIITEVNPIRALEAKMDGFTVLKMEEAAKIGDIFVTTTGCKDILRMEHFLLMKDGAVLSNAGHFDNEINKNDLKELSKSVKEAR
FNIEEYDLGNKKIYLLGEGRLVNLACADGHPCEVMDMSFANQALSAKFIKENKGKLENEVYEIPYEQDFKIALLKLHSMGADIDE
LSPEQRKYLSDWKEGT

>*Mm*SAHH

MGSSHHHHHHSSGLVPRGSHMSDKLPYKVADIGLAAWGRKALDIAENEMPGLMRMREMYSASKPLKGARIAGCLHMTVET
AVLIETLVALGAEVRWSSCNIFSTQDHAAAAIAKAGIPVFAWKGETDEEYLWCIEQTLHFKDGPLNMILDDGGDLTNLIHTKYP
QLLSGIRGISEETTTGVHNLYKMMSNGILKVPAINVNDSVTKSKFDNLYGCRESLIDGIKRATDVMIAGKVAVVAGYGDVGKGC
AQALRGFGARVIITEIDPINALQAAMEGYEVTTMDEACKEGNIFVTTTGCVDIILGRHFEQMKDDAIVCNIGHFDVEIDVKWLN
ENAVEKVNIKPQVDRYWLKNGRRIILLAEGRLVNLGCAMGHPSFVMSNSFTNQVMAQIELWTHPDKYPVGVHFLPKKLDEAV
AEAHLGKLNVKLTKLTEKQAQYLGMPINGPFKPDHYRY

>*Pa*SAHH

MGSSHHHHHHSSGLVPRGSHMSAVMTPAGFTDYKVADITLAAWGRRELIIAESEMPALMGLRRKYAGQQPLKGAKILGCIH
MTIQTGVLIETLVALGAEVRWSSCNIFSTQDQAAAAIAAAGIPVFAWKGETEEEYEWCIEQTILKDGQPWDANMVLDDGGDL
TEILHKKYPQMLERIHGITEETTTGVHRLLDMLKNGTLKVPAINVNDSVTKSKNDNKYGCRHSLNDAIKRGTDHLLSGKQALVIG
YGDVGKGSSQSLRQEGMIVKVAEVDPICAMQACMDGFEVVSPYKNGINDGTEASIDAALLGKIDLIVTTTGNVNVCDANMLK
ALKKRAVVCNIGHFDNEIDTAFMRKNWAWEEVKPQVHKIHRTGKDGFDAHNDDYLILLAEGRLVNLGNATGHPSRIMDGSF
ANQVLAQIHLFEQKYADLPAAEKAKRLSVEVLPKKLDEEVALEMVKGFGGVVTQLTPKQAEYIGVSVEGPFKPDTYRY

>*Pfu*SAHH

MGSSHHHHHHSSGLVPRGSHMDCGKDYCVKDLSLAEEGWKKIDWVSRFMPVLQYIKREFEEKKPFKGVRIAATLHLEMKTAF
LLLTLKAGGAEVSAAASNPLSTQDDVVAALAKAGVKVYAIRGESREQYYEFMHKALDIRPNIIIDDGADMISLVHKERQEMLDEI
WGGSEETTTGVIRLRAMEKAGILKFPVIAVNDSYMKYLFDNRYGTGQSTWDGIMRATNLLIAGKNVVVVGYGWCGRGIAMR
ARGLGATVIVVEVDPIKALEARMDGFLVMDMKEAAKIGDIFVTATGNIKCIRREHFELMKDGAIMANAGHFDVEIWKPDLEKL
AVEINNPRPNVTEYKLKDGRRLYLLADGRLVNLVAADGHPAEIMDMSFALQAKAAEYIKDNHERLEPKVYILPREIDEMVARIKL
ESMGIKIEELTEEQKKYLESWEHGT

>*Sac*SAHH

MGSSHHHHHHSSGLVPRGSHMDYRVKDLSLAEQGRKQIEWAELHMPALMEIRKRFNAEKPLDGIRIGAVLHVTKETAVLVET
LKAGGAEIALAGSNPLSTQDDVAAGLAKNGIHVYAWRGETEKDYYDNIREILKYEPHVIMDDGGDLHAYVHENNLTSKIVGGT
EETTTGVIRLKAMEEEKVLKYPVIAVNNAFTKYLFDNRIGTGQSTIDGILRATNILIAGKVAVVIGYGWVGRGIASRFKGMGARVI
VVESSPFRALEALMDGFDVMTMNRASEIGDIFVTATGNLNVVSRDHILRMKDGAVLANSGHFNVEIDVKGLKEISVETREVRQ
NLEEYKLRNGKRIYLLADGRLVNLVAAEGHPSEVMDLSFCNQALSVEHLIKNKGKLENKVYNVPIEIDEQVARLKLKALGIEIEELTI
EQKEYIKQWKYGT

>*Sa*SAHH

MGSSHHHHHHSSGLVPRGSHMASAQQHDFKVADLSLAEFGRKEITLAEHEMPGLMSIREEYAASQPLAGARVTGSLHMTVQ
TAVLIETLTALGAEVRWASCNIFSTQDHAAAAIAVGPNGTPDNPQGVPVFAWKGESLEEYWWCTEQALTWPNTPTGGPNMI
LDDGGDATLLVHNGVQYEKDGKVPDPVTAESDEHRVILQLLTRTLGENPQKWTQLASEIRGVTEETTTGVHRLYEMQRDGQL
LFPAINVNDAVTKSKFDNKYGCRHSLIDGINRATDVLIGGKTAVVCGYGDVGKGCAESLRGQGARVIVTEIDPICALQAAMDGY
QVATLDDVIGQADIFVTTTGNKDIIMAADMAKMKHQAIVGNIGHFDNEIDMAGLAAIPGIVKDEVKPQVHTWTFPDGKVIIVL
SEGRLLNLGNATGHPSFVMSNSFADQTLAQIELFTKPDEYPTDVYVLPKHLDEKVARLHLDALGVRLTTLRPEQAAYIGVSVEGP
FKPDHYRY

>*Sf*SAHH

MGSSHHHHHHSSGLVPRGSHMTTTSTTGHDFKVADLSLAAFGRKEITLAEHEMPGLMAIRKEYSAEKPLAGARITGSLHMTVQTAVLIETLVA
LGAEVRWASCNIFSTQDHAAAAIAVGPDGTPDNPRGVPVFAWKGETLEEYWWCTEQALTWPNTPTGGPNMILDDGGDATLLVHKGVEYE
KAGAAPSVDTAENDEHRVILQLLNRTLAESPQKWTQPASEIRGVTEETTTGVHRLYEMQQAGTLLFPAINVNDAVTKSKFDNKYGCRHSLIDG
INRATDVLIGGKTAVVCGYGDVGKGCAESLRGQGARVMITEIDPICALQAAMDGYQVVRLDDVVETADIFITTTGNKDIIMASDMAKMKHQ
AIVGNIGHFDNEIDMAGLAAIDGIVKDEVKPQVHTWTWPDGKSIIVLSEGRLLNLGNATGHPSFVMSNSFANQTIAQIELFTKPESYPTDVYVL
PKHLDEKVARLHLDALGAKLTTLRPEQAAYIGVPVEGPYKPDHYRY

>*Sso*SAHH

MGSSHHHHHHSSGLVPRGSHMSYKIKDLSLASEGKKQIEWAERHMPTLMEIRKRFKAEKPLKGINISAVLHVTKETAALVKTLKI
GGANVALAGSNPLSTQDDVAAALVEEGISVFAWKGENETEYYSNIESIVKIHEPNIVMDDGADLHAYIHEKVSSKLDIYGGTEET
TTGVIRLKAMEKDGVLKYPLVAVNNAYTKYLFDNRYGTGQSAIDGILRATNILIAGKIAVVAGYGWVGRGIANRLRGMGARVIV
TEVDPIRALEAVMDGFDVMPIAEASKVGDIFVTATGNTKAIRVEHMLNMKDGAILSNAGHFNVEVDVKGLKETAVKVRNIRPY
VDEYTLPNGKRVYLLADGRLVNLAAAEGHPSEVMDMSFANQALAVEYLVKNRGKLEKKVYNMPMELDYEVARIKLKSMGIQI
DELTEEQKEYLEQWKSGT

>*Tk*SAHH

MGSSHHHHHHSSGLVPRGSHMDCTKDYCVKDISLAPSGEKKIDWVSRFMPVLQHIRKDFEERKPFKGVRIAATLHLEMKTAFL
LLTLKAAGAEVSAAASNPLSTQDDVVAALAKAGVKVYAIRGEDREQYYEFMHKALDVKPNIIIDDGADMVSTVLKERQELIPEI
WGASEETTTGVIRLRAMEKDGVLKFPIIAVNDSYTKYLFDNRYGTGQSTWDGIIRTTNLLVAGKNVVVVGYGWCGRGIAMRA
RGLGATVIVVEVDPIRALEARMDGFLVMDMMEAAKVGDIFITATGDINCIRKEHFELMKDGAILANAGHFDVEISKPDLEALAV
EISEPRPNITEYKMADGRRLYLLAEGRLVNLAAADGHPAEIMDMSFALQAKAAEYIKENRGRLEPKVYVLPREIDEMVARIKLAS
MGIKIEELTEEQKKYLESWEHGT

>*Tm*SAHH

MGSSHHHHHHSSGLVPRGSHMNTGEMKINWVSRYMPLLNKIAEEYSREKPLSGFTVGMSIHLEAKTAYLAITLSKLGAKVVIT
GSNPLSTQDDVAEALRSKGITVYARRTHDESIYRENLMKVLDERPDFIIDDGGDLTVISHTEREEVLENLKGVSEETTTGVRRLKA
LEETGKLRVPVIAVNDSKMKYLFDNRYGTGQSTWDAIMRNTNLLVAGKNVVVAGYGWCGRGIALRAAGLGARVIVTEVDPV
KAVEAIMDGFTVMPMKEAVKIADFVITASGNTDVLSKEDILSLKDGAVLANAGHFNVEIPVRVLEEIAVEKFEARPNVTGYTLEN
GKTVFLLAEGRLVNLAAGDGHPVEIMDLSFALQIFAVLYLLENHRKMSPKVYMLPDEIDERVARMKLDSLGVKIDELTEKQRRYL
RSWQ
